# Brain capillary pericytes exert a substantial but slow influence on blood flow

**DOI:** 10.1101/2020.03.26.008763

**Authors:** David A. Hartmann, Andrée-Anne Berthiaume, Roger I. Grant, Sarah A. Harrill, Tegan Noonan, Jordan Costello, Taryn Tieu, Konnor McDowell, Anna Faino, Abigail Kelly, Andy Y. Shih

## Abstract

The majority of the brain’s vasculature is comprised of intricate capillary networks lined by capillary pericytes. However, it remains unclear whether capillary pericytes contribute to blood flow control. Using two-photon microscopy to observe and manipulate single capillary pericytes *in vivo*, we find their optogenetic stimulation decreases lumen diameter and blood flow, but with slower kinetics than mural cells of upstream pial and pre-capillary arterioles. This slow, optogenetically-induced vasoconstriction was inhibited by the clinically-used vasodilator fasudil, a Rho kinase inhibitor that blocks contractile machinery. Capillary pericytes were also slower to constrict back to baseline following hypercapnia-induced dilation, and relax towards baseline following optogenetically-induced vasoconstriction. In a complementary approach, optical ablation of single capillary pericytes led to sustained local dilation and a doubling of blood cell flux in capillaries lacking pericyte contact. Altogether these data indicate that capillary pericytes contribute to basal blood flow resistance and slow modulation of blood flow throughout the capillary bed.

## INTRODUCTION

The adult human brain contains more than 400 miles of vasculature, tasked with delivery of oxygen and nutrients to billions of brain cells. Understanding how the cerebrovasculature achieves this feat is critical because insufficient blood flow is involved in the pathogenesis of numerous neurologic conditions including stroke, vascular cognitive impairment, and Alzheimer’s disease.^1^ The distribution of blood relies on a dense capillary network, through which all red blood cells (RBCs) must percolate in a single-file fashion during their transit from arteries to veins.^2^ With this arrangement, it is no surprise that the capillary bed possesses the highest flow resistance within the cerebrovasculature.^2, 3, 4^ This flow resistance must be carefully tuned for proper allocation of blood to all regions of the brain. Despite their physiological and clinical importance, the factors that govern brain capillary blood flow in the living brain remain largely unknown.

Collectively referred to as vascular mural cells, smooth muscle cells (SMCs) and pericytes line the entire cerebrovasculature, but occupy distinct microvascular zones.^5, 6^ On arterioles, concentric rings of SMCs produce powerful constrictions or dilations to alter blood rapidly. These SMC dynamics are critical for blood flow control during neurovascular coupling, where fluctuating neuronal activity produces commensurate changes in blood supply within seconds.^7^ Meanwhile, in capillary networks, pericytes are the predominant mural cell form. Pericytes are characterized by protruding ovoid cell bodies and elongated processes that partially cover the capillary surface.^8^ Their shared embryology with SMCs, as well as their anatomical location within the capillary basement membrane, have led many to hypothesize that pericytes modulate blood flow at the capillary level. However, this hypothesis has remained controversial since pericytes were discovered by Rouget and Eberth in the 1870s.^9, 10^ Seminal *ex vivo* rodent brain slice studies first demonstrated that brain pericytes could alter capillary diameter in response to electrical stimulation, neurotransmitters, and ischemia.^11^ Yet, studies seeking to corroborate these responses *in vivo* have resulted in mixed outcomes, with some studies supporting modulation of capillary blood flow by brain pericytes^12, 13, 14^, and others suggesting the contrary.^7, 15, 16^

Two salient issues underlie the conflicting reports from *in vivo* studies. First, at least three unique mural cell types reside on brain arterioles, pre-capillary arterioles, and capillaries, respectively.^5, 17^ SMCs on arterioles and ensheathing pericytes on pre-capillary arterioles clearly control blood flow.^7^ Both cell types express α-SMA, a protein that is associated with SMC contractility. However, “ensheathing pericytes” are more elongated and have an ovoid cell body that protrudes from the vessel wall, a trait that resembles classically-described pericytes.^5^ In contrast, capillary pericytes exist deeper in the microvascular network (capillary bed) and their α-SMA expression is absent or lower than upstream mural cells. Capillary pericytes have long, thin processes that incompletely cover the endothelium.^5^ Many prior studies have not semantically distinguished between ensheathing and capillary pericytes, fueling some uncertainty about whether capillary pericytes regulate blood flow.^12, 14, 18, 19^ This is critical to clarify because capillary pericytes contact the vast majority of the length of cerebrovasculature and can therefore profoundly influence blood flow. Further, the loss or dysfunction of capillary pericytes has been implicated in blood flow abnormalities during stroke^7, 12, 20, 21^, dementia^22, 23, 24^, epilepsy^25^, and other neurological diseases.^25, 26^

The second issue in studying the role of pericytes *in vivo* is the inherent connectivity of the cerebrovasculature. Because capillary blood flow dynamics are so greatly affected by upstream arterioles, the autonomous ability of capillary pericytes to control flow is difficult to determine using observational approaches. Several previous studies correlated local changes in capillary diameter with the presence (or absence) of pericyte somata.^12, 13, 15, 16^. Not only is it difficult to disentangle the influence of upstream flow changes with this strategy, but this approach is based on the assumption that only the soma of pericytes can exert tone on capillaries. Furthermore, even in animal models that can genetically target capillary pericytes, the structural and functional connectivity between pericytes, astrocytes, and endothelial cells means that deleting large numbers of pericytes will affect multiple cell types. In one recent study, specific deletion of large numbers of pericytes in the adult brain caused substantial blood-brain barrier breakdown and edema, which was believed to be the primary cause of reduced cerebral blood flow in these animals, rather than the loss of pericyte control of blood flow.^27^ Other interventions such as microvascular occlusion produce capillary constriction that slows blood flow and contributes to infarct growth. However, it remained unclear whether pericytes or other factors such as edema were the basis of this pathological capillary constriction.^28^ Therefore, “cause-and-effect” studies that manipulate capillary pericytes in a cell-specific manner are needed to fully understand how capillary pericytes contribute to blood flow control. To better isolate the role of pericytes in capillary constriction, Hill *et al.* used optogenetics to selectively probe the contractility of mural cells in different microvascular zones.^7^ Their studies concluded that capillary pericytes lacked the structural and biochemical profile (α-SMA expression) necessary to manipulate capillary diameter *in vivo.* However, their optogenetic manipulations did not test whether the cells were sufficiently stimulated to unveil contractile ability. Further, their experiments focused on the rapid vasodynamics relevant to neurovascular coupling, rather than on the basal contractile tone that might help create enduring flow resistance and flow patterns within the capillary bed.

In this study, we examined whether capillary pericytes provide vascular tone and flow resistance *in vivo*. We overcome prior issues of pericyte identification by making careful distinctions between microvascular zones of adult mouse cortex, and by using cell morphology and vascular branch order criteria to unambiguously identify capillary pericytes *in vivo*. Further, we study capillary pericytes in a cause-and effect manner, by using spatially localized two-photon optical manipulations to either stimulate or ablate individual capillary pericytes. This disentangles their local influence on capillary diameter from the influence of flow in upstream vessels.

## RESULTS

### Distinguishing among the different mural cells of pre-capillary and capillary zones

We bred PDGFRβ-Cre mice^29^ with fluorescent reporter mice^30^ to achieve robust fluorescent labeling of >99% of all mural cells throughout the cerebrovascular tree, as previously reported (**Fig. 1a**).^5, 6^ To understand the extent of cortical vasculature that was contacted by pericytes with low or absent α-SMA, *i.e.*, capillary pericytes, we immunostained 0.5-1 mm thick coronal brain slices for α-SMA (**Fig. 1b**). We then performed SeeDB tissue clearing^31^ followed by volumetric two-photon imaging of cleared slices of sensory cortex. The thick tissue sections preserved microvascular architecture and allowed us to assess α-SMA expression in mural cells along ~116 mm of total vascular length in three dimensions. This revealed that the vast majority of vascular length (96 ± 2%) was covered by mural cells with undetectable levels of α-SMA, which are nearly all capillary pericytes, but also include a small number of cells on venules that have not yet been defined (**Fig. 1c**).

**Figure 1:**
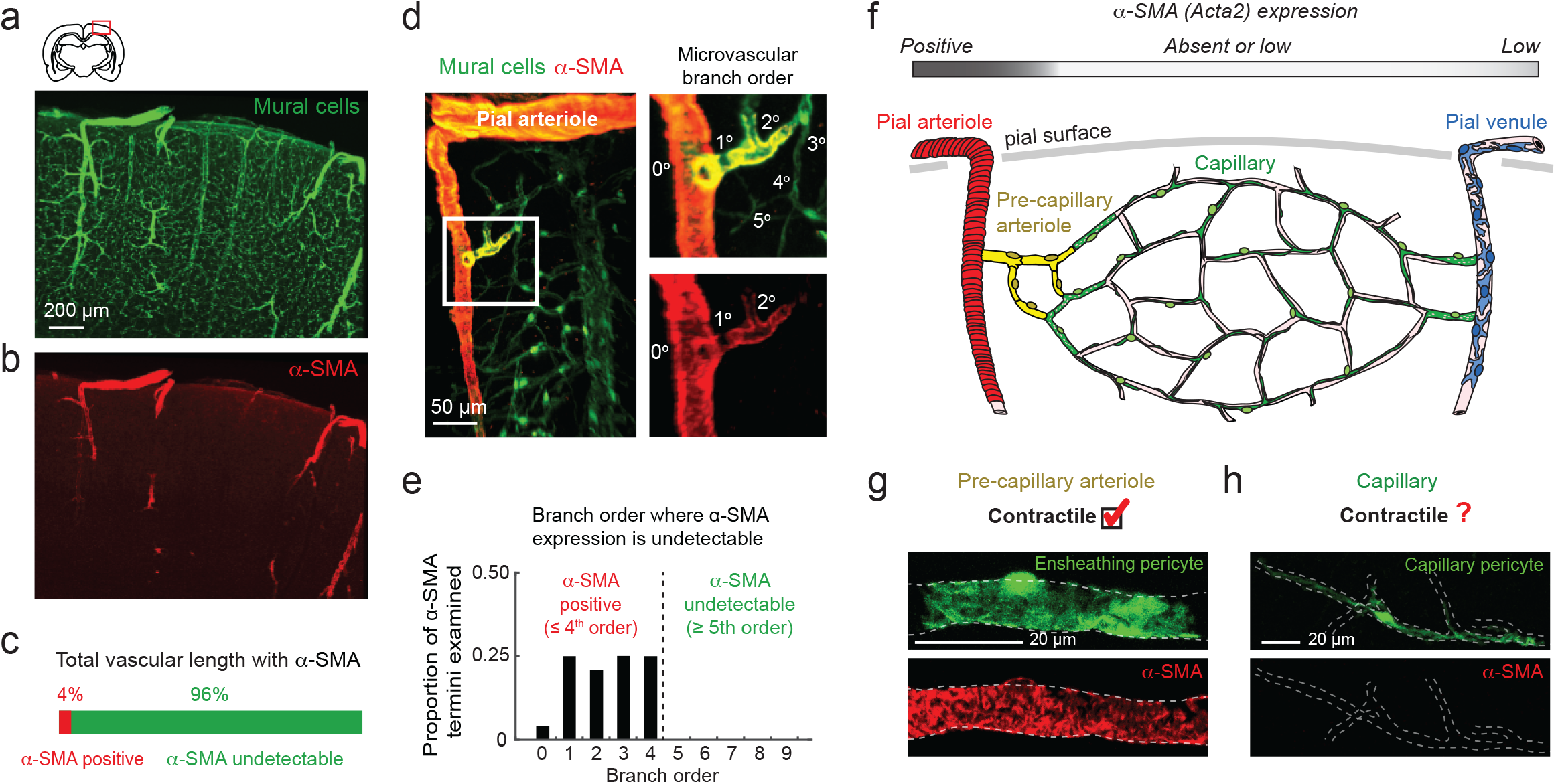
Capillary pericytes occupy the vast majority of mouse cerebrovasculature. (**a,b**) A thick section of the mouse cerebral cortex, optically cleared and imaged with two-photon microscopy, shows genetically-labeled mural cells (PDGFRβ-tdTomato) in green and α-SMA immunostaining in red. (**c**) Proportion of vascular length associated with α-SMA-positive mural cells (smooth muscle cells and ensheathing pericytes) versus α-SMA-negative or low mural cells (predominantly capillary pericytes, but a small proportion of venular SMCs). A total of 116 mm of cortical vascular length was quantified across 4 volumetric data sets from 2 mice. Each data set contained a central penetrating arteriole and surrounding capillary networks, as seen in panel (d). (**d**) Drop in α-SMA expression as pre-capillary arterioles transition into capillaries. The example shows a penetrating arteriole offshoot with α-SMA termination occurring at the second order branch. (**e**) Histogram showing proportion of branch orders at which α-SMA decrease occurs in the upper 350μm of cortex. N = 24 α-SMA terminations (2 mice, 11 penetrating arteriole networks). (**f**) Schematic depicting zones in cortical microvasculature. Red shows SMCs on pial and penetrating arterioles, yellow shows ensheathing pericytes on pre-capillary arterioles, green shows capillary pericytes, and blue shows SMCs of venules. Expression of α-SMA decreases sharply between pre-capillary arteriole and capillary zones. (**g,h**) Contractility is established in previous literature for mural cells of pre-capillary arterioles (called ensheathing pericytes), but remains uncertain for capillary pericytes, which reside in the capillary zone.

The expression level of α-SMA drops abruptly as pre-capillary arterioles transition into capillaries (**Fig. 1d**).^5, 7, 16^ To ensure that our subsequent *in vivo* imaging experiments were targeting the capillary zone, which we define as vessels contacted by capillary pericytes^5, 7^, we related mural cell α-SMA content to microvascular branch order. We considered penetrating arterioles as 0^th^ branch order, and the first offshoot from the penetrating arteriole as 1^st^ order. Each successive vessel bifurcation then increased branch order by 1. To determine the maximum branch order at which α-SMA expression was detectable, we followed 24 penetrating arteriole offshoots. These offshoots were examined in the upper 350 μm of cortex, which was most accessible to subsequent *in vivo* imaging studies. This revealed that α-SMA expression decreased at or before the 4^th^ branch order (**Fig. 1e**), a finding consistent with our past work and two other independent groups using similar tissue fixation techniques.^7, 16^ Thus, targeting microvessels of 5^th^ branch order or greater during *in vivo* imaging studies ensures the examination of capillary pericytes with undetectable levels of α-SMA. This targets roughly the center of the capillary bed, where the majority of oxygen delivery to the tissue occurs.^32, 33^

**Figure 1f-h** illustrates the central question of our study: Do capillary pericytes influence local blood flow *in vivo*? Compared to mural cells on pre-capillary arterioles, which we refer to as “ensheathing pericytes^5^”, capillary pericytes (i) individually contact a greater length of the microvasculature, (ii) are distinct in morphology and in transcriptional profile^17^, and (iii) express little to no α-SMA. These cells are the focus of our studies because they occupy the vast majority of the vasculature, and relatively little is known about their role in controlling blood flow.

### Concurrent blood flow imaging and two-photon optogenetic stimulation of capillary pericytes

To directly assess the contractile ability of capillary pericytes *in vivo*, we crossed PDGFRβ-Cre mice with reporter mice for the light-gated ion channel, ChR2-YFP^34^ (**Supplementary Fig. 1a,c**). To confirm that hemodynamic changes were the result of ChR2 activation, rather than the non-specific effects of laser irradiation, we also bred PDGFRβ-Cre mice with reporter mice for either cytosolic YFP or membrane-bound GFP (mT/mG) (**Supplementary Fig. 1b,d**). The mT/mG-expressing mice allowed us to assess whether pericytes with membrane localization of fluorophore could respond to light differently than pericytes with a cytosolic fluorophore. Ultimately, we pooled the data from control mice into a single group, as no differences were observed between YFP and mT/mG groups (**Supplementary Fig. 2**).

In adult offspring from these crosses, we generated acute, skull-removed cranial windows over sensory cortex for two-photon imaging, and injected fluorescent dextran dye retro-orbitally to visualize the vasculature (i.v. dye; Texas red-dextran, 70 kDa)(**Fig. 2a**). Unless otherwise noted, experiments were conducted using light (0.8% MAC) isoflurane anesthesia delivered in air. Capillary pericytes in PDGFRβ-ChR2-YFP mice covered the endothelium to a greater extent than did pericytes in PDGFRβ-YFP mice, suggesting that capillary pericytes may be more structurally equipped to control capillary diameter than previously illustrated by cytosolic fluorophores such as YFP (**Supplementary Fig. 1e,f**). The morphology shown in ChR2-YFP pericytes is consistent with 3-dimensional ultrastructural studies that have revealed thin sheet-like outcroppings from the central axis of pericyte processes.^35, 36^

**Figure 2:**
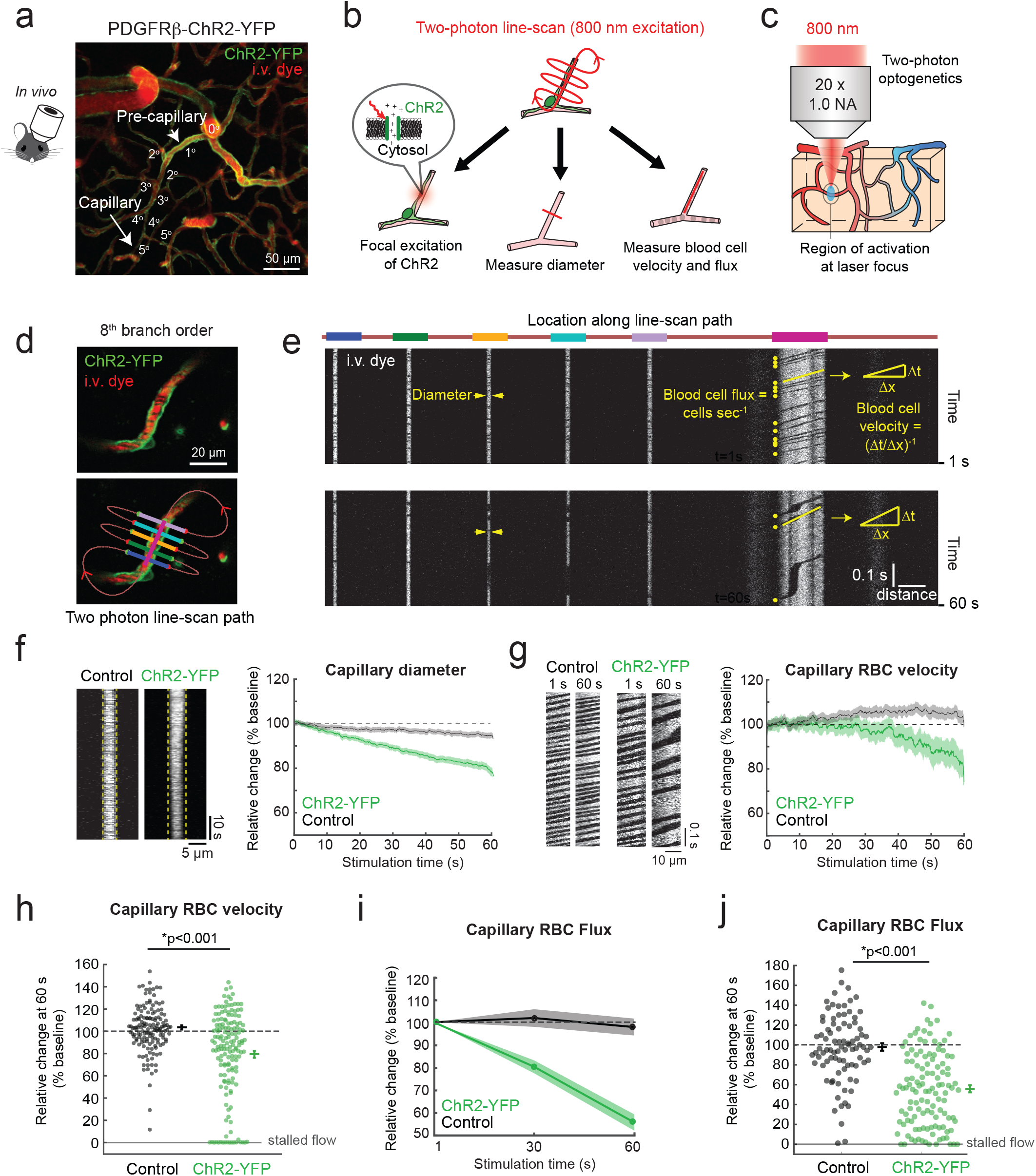
Optogenetic activation of capillary pericytes constricts capillaries and reduces blood flow. **(a)** *In vivo* two-photon image of cortical vasculature in a PDGFRβ-ChR2-YFP mouse. Branch order from the penetrating arteriole (0^th^ order) is used to verify examination of the capillary zone (≥ 5 order). The image is a projection over 150 μm of cortical depth. **(b)** Specialized two-photon line-scans (red) used to simultaneously stimulate ChR2-YFP in capillary pericytes and measure capillary vasodynamics. **(c)** Two-photon excitation of ChR2-YFP with focused 800 nm light allows focal activation of pericytes below the brain surface. **(d)** Example of an 8^th^ branch order capillary pericyte in ChR2-YFP mouse with the line-scan path traveled by the two-photon laser. Each colored line in panel (d) corresponds to the same colored line at the top of panel (e), which corresponds to regions of constant scan velocity used for data extraction. **(e)** Example of raw line-scan data, collected by repeatedly scanning the path shown in panel (d). The lateral dimension is distance along the line-scan path, and the vertical dimension is time. The upper image shows data at the beginning of the line-scan, and the lower image shows the data at the end of a continuous 60 second line scan. “Distance” scale is 20 μm. A decrease in RBC velocity manifests as increased time to travel the same distance, and hence a greater slope of the RBC shadow. A decrease in RBC flux manifests as fewer dark streaks within the line-scan. **(f)** *Left:* Example i.v. dye image showing diameter portion of line-scan over 60 seconds of stimulation. A comparison is provided for capillaries in control mice (pericytes expressing YFP or mGFP) versus ChR2-YFP mice; baseline diameter shown as vertical dotted yellow line. *Right:* Aggregate data showing greater average diameter decrease during two-photon stimulation of capillaries in ChR2-YFP mice, compared to control mice. N = 155 (10 mice) capillaries for ChR2-YFP group; n = 145 (9 mice) capillaries for control group). **(g)** *Left:* Example i.v. dye image showing RBC velocity (slope of blood cell shadow) and flux (number of blood cell shadows) in the control animals does not change over duration of scan, whereas a decrease is observed during stimulation in ChR2-YFP animals. *Right:* Aggregate data for RBC velocity change during stimulation of capillaries in ChR2-YFP and control mice. N = 160 (10 mice) capillaries for ChR2-YFP group; n = 152 (9 mice) capillaries for control group). **(h)** Bee swarm plot of RBC velocity at 60 seconds relative to baseline values in ChR2-YFP and control groups. *F(1,290)=26.8, p<0.001, repeated-measures ANOVA. N = 157 (10 mice) capillaries for ChR2-YFP genotype; n = 152 (9 mice) capillaries for control genotype. **(i)** Relative RBC flux decreases during stimulation of capillaries in ChR2-YFP mice, but not control mice. **(j)** Bee swarm plot of blood cell flux at 60 seconds relative to baseline values in ChR2-YFP and control groups. *F(1,186)=42.48, p<0.001 by repeated measures ANOVA, n = 118 (10 mice) capillaries for ChR2-YFP group; n = 87 (9 mice) capillaries for control group. Mean ± S.E.M. is shown for all time-course data and swarm plots.

We found that the mere observation of cortical capillaries in PDGFRβ-ChR2-YFP mice using 800 nm two-photon excitation was sufficient to induce capillary constriction. However, an excitation wavelength of 900 nm did not lead to appreciable constrictions (**Supplementary Fig. 3**),in line with previous reports.^7^ We therefore navigated through our sample using 900 nm excitation during tracking of capillary branch order, and only switched to 800 nm for optogenetic activation once a target capillary pericyte was identified.

To concurrently activate ChR2-YFP and measure blood flow dynamics, we used multi-segmented two-photon laser line-scan paths that performed three functions (**Fig. 2b**). First, two-photon irradiation at 800 nm activated pericyte ChR2 primarily within the imaging plane, leading to spatially restricted pericyte activation while avoiding coincident activation of upstream SMCs (**Fig. 2c**).^7^ Second, traversing the capillary lumen 5 times illuminated the i.v. dye, and provided multiple measurements of vessel diameter. Third, the line-scan bisected the central axis of the capillary lumen to track the movement of blood cell shadows, from which we extracted the RBC flow velocity and RBC flux (blood cells per second). An *in vivo* example of this line-scan being applied to a capillary is shown in **Fig. 2d**, and the resulting space-time plot of the entire scan is provided in **Fig. 2e** with annotation of where lumen diameter and RBC flow metrics were collected (**Supplementary Movies 1 and 2**).

### Capillary pericytes contract and reduce capillary flow in response to optogenetic activation

When we irradiated capillaries (5^th^ to 9^th^ branch order) for 60 seconds with two-photon line-scans, we observed a gradual decrease in capillary diameter in the ChR2-YFP group (~20% below baseline). This was primarily mediated by ChR2-YFP activation, and not by laser irradiation, because the constriction was substantially smaller in controls (~5%) (**Fig. 2f**). A clear deviation in diameter was evident within 10 to 20 seconds of stimulation. Laser intensity as a function of cortical depth was well-matched between ChR2-YFP and control groups, eliminating the possibility that the groups received different levels of irradiation (**Supplementary Fig. 4**). Critically, vasoconstriction was associated with a decrease in RBC velocity and flux in the same capillaries, showing that the induced levels of constriction were sufficient to alter blood flow. It also confirmed that the diameter changes were not simply due to the capillary shifting from imaging focal plane (**Fig. 2g-j**). This critical result was independent of the anesthesia used, as we obtained a similar outcome with animals lightly sedated by chlorprothixene (**Supplementary Fig. 5a,b**).

Our data support the idea that capillary flow reduction was due to locally-induced pericyte contraction, as opposed to a passive downstream deflation that might occur if stimulation had conducted to upstream arterioles.^37^ Capillary diameter deviated from baseline earlier than did blood cell velocity, *i.e.* within 10 seconds for lumen diameter (**Fig. 2f**), vs. 30 seconds for RBC velocity (**Fig. 2g**). If upstream arteriole constriction had triggered a downstream deflation of capillaries, one would instead expect RBC velocity change to precede diameter change.^38^

During optogenetic stimulation with line-scans, we observed complete cessation of blood flow in 12% of capillaries from ChR2-YFP mice, but none from control mice. To better understand the persistence of these flow stalls, we also collected full-field image stacks at 900 nm 1-3 minutes after the 60 second stimulation period for all vessels imaged. We observed sustained stalling of RBCs in ~30% of capillaries from ChR2-YFP mice, but in only 3% of capillaries from control mice (**Supplementary Fig. 6**). Capillaries that had smaller basal diameter and lower RBC velocity and flux were more likely to stall (**Supplementary Fig. 7a-c**). Capillaries with flow stalls were not appreciably different with respect to branch order from the penetrating arteriole (**Supplementary Fig. 7d**). Collectively, these data support the idea that capillary pericytes are sufficient to constrict capillaries and maintain this state of contraction on a protracted time-scale.

Previous studies have investigated pericyte contractility by comparing diameter changes in vessel segments beneath the soma to those beneath the processes, finding that contractility occurred preferentially at the pericyte soma.^12, 13, 15^ To test whether this effect was recapitulated using an optogenetic approach, we divided our ChR2-YFP data set into two groups, “soma” or “no soma”, corresponding to whether any portion of the multi-segment line-scan transected the pericyte soma or not (**Supplementary Fig. 8a,b**). Somata were easily detected using three-dimensional image stacks collected before and after each scan. These two groups showed no detectable difference in optogenetically-induced decrease in capillary diameter, RBC velocity or RBC flux (**Supplementary Fig. 8c-e**), suggesting that capillary pericytes can be stimulated to exert force on capillaries at their somata and their far-reaching processes. Considering that 93% of the endothelial contact made by capillary processes is through their extensive processes, this suggests a broad potential impact on capillary flow by pericytes.^39^

In 30% of optogenetically stimulated capillary pericytes, we observed localized blebbing (small membrane protrusions) along the pericyte wall, possibly due to reorganization of the actin cytoskeleton that lines the plasma membrane (**Supplementary Fig. 9a-e**). These local protrusions were reminiscent of blebs described in the process of contraction to endothelin^40^ and contractility-driven cell migration^41^. The observed blebbing occurred within one minute of stimulation and is thus unlikely to be an indication of migration. It is also unlikely that the blebbing was the result of photodamage because it was not observed in control mice that were exposed to the same laser intensities. Critically, pericyte blebbing did not drive the genotype effects on vasodynamics, as omitting these cells from the data set had no effect on significance of capillary flow reduction in ChR2-YFP expressing mice (**Supplementary Fig. 9f,g**).

### Fasudil blocks optogenetically-induced capillary constriction

We next asked whether capillary constriction by pericytes relied upon cellular contractile machinery. The Rho/ROCK inhibitor fasudil inhibits mural cell contraction by blocking actomyosin cross-bridging. It also acts by inhibiting α-SMA-independent smooth muscle cell contraction that occurs via polymerization of cytoskeletal actins, which might be relevant for capillary pericytes that have little to no α-SMA.^42^ Fasudil does not alter conductance of L-type calcium channels in mural cells^43^, which are essential mediators of mural cell membrane voltage changes induced by ChR2 activation.^44^ Thus, fasudil is expected to inhibit mural cell contractile machinery without influencing their optogenetic activation.^43^

Fasudil was incorporated into an agarose layer overlying the cortical surface during preparation of an acute cranial window, with concentrations 10-fold (1 mM) and 100-fold (10 mM) higher than a commonly-used *ex vivo* concentration.^45^ This allowed for direct access to the brain and accounted for drug dilution as it crossed the intact dura and entered the brain interstitial fluid.^46^ Under basal conditions, fasudil caused dilation of pial arterioles and capillaries, but without any effects on capillary RBC velocity (**Supplementary Fig. 10**). However, fasudil increased baseline capillary RBC flux, which can pack RBCs more tightly, creating a ‘traffic jam’ that limits RBC velocity (**Supplementary Fig. 11**).^47^ During optogenetic stimulation of capillary pericytes, fasudil attenuated relative and absolute capillary constriction in a dose-dependent manner (**Fig. 3a, Supplementary Fig. 10e,f**). Furthermore, fasudil abolished the reduction of RBC velocity and flux caused by optogenetic pericyte stimulation (**Fig. 3b,c**). Accordingly, blockade of constriction also led to a dose-dependent alleviation of persistent flow stalls (**Supplementary Fig. 12**). The incidence of pericyte blebbing was also markedly reduced in line with the idea that the membrane protrusions were a Rho/ROCK-dependent process involving actin cytoskeleton reorganization (**Supplementary Fig. 13**).^48^ Collectively, these data support the idea that optogenetic stimulation of capillary pericytes engages a form of cellular contractile machinery that can be pharmacologically modulated.

**Figure 3:**
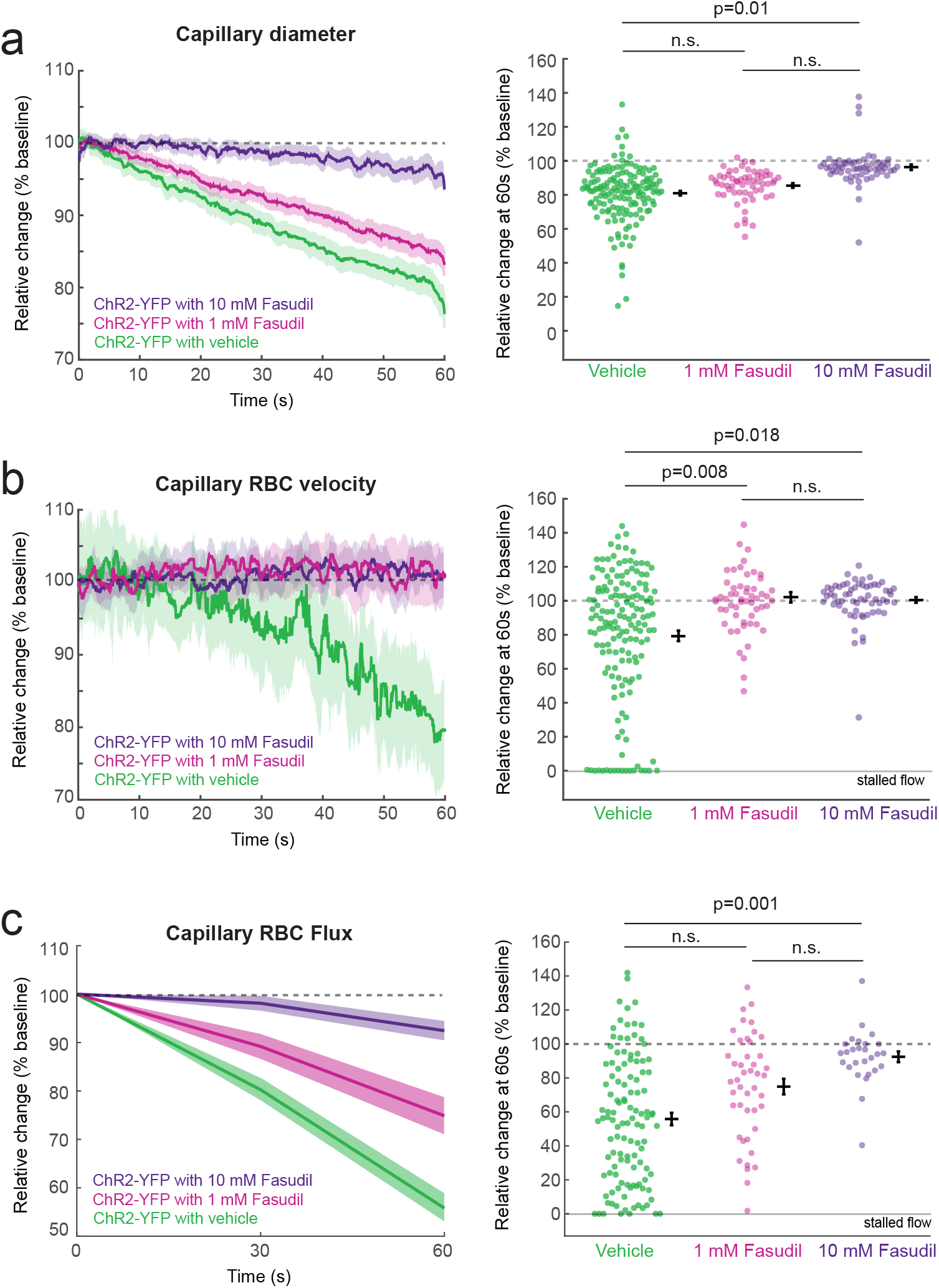
Topical administration of fasudil prevents hemodynamic changes induced by optogenetic stimulation of capillary pericytes. **(a)** *Left:* Fasudil dose-dependently abrogated reduction in capillary diameter during ChR2-YFP stimulation in capillary pericytes. *Right:* Bee swarm plot of relative diameter change from baseline with and without fasudil. N = 154 (10 mice), 60 (3 mice), 60 (3 mice) for vehicle, 1 mM fasudil, 10 mM fasudil, respectively. **(b)** *Left:* Fasudil abrogated decrease of capillary RBC velocity during ChR2-YFP stimulation in capillary pericytes. *Right:* Bee swarm plot of relative RBC velocity change from baseline with and without fasudil. N = 157 (10 mice), 57 (3 mice), 61 (3 mice) for vehicle, 1 mM fasudil, 10 mM fasudil, respectively. **(c)** *Left:* Fasudil dose-dependently abrogated decrease in capillary RBC flux during ChR2-YFP stimulation in capillary pericytes. *Right:* Bee swarm plot of relative RBC flux change from baseline with and without fasudil. N = 118 (10 mice), 45 (3 mice), 27 (3 mice) for vehicle, 1 mM fasudil, 10 mM fasudil respectively. Mean ± S.E.M. is shown for all time-course data and swarm plots. P values shown in bee swarm plots from repeated measures ANOVA adjusted by Tukey *post hoc* test (overall ANOVA test p values all <0.015).

### Capillary pericytes constrict more slowly than upstream mural cells with high α-SMA content

Although optogenetic stimulation of capillary pericytes produced hemodynamic changes, the kinetics were slow. In the same mice, we compared the contractile dynamics of capillary pericytes against mural cells of other microvascular zones by applying a similar line-scan pattern to other vessel types. We found that stimulation of pial arteriole SMCs in ChR2-YFP mice led to a more robust and rapid vasoconstriction than that seen in capillary pericytes. We observed a maximal arteriolar constriction of 40% within the first 25 seconds of stimulation, followed by gradual relaxation back to baseline despite continued stimulation (**Fig 4a**). This rebound may be due to a negative feedback mechanism induced by strong vessel contraction.^49^ Stimulation of pial arterioles in control animals exhibited a smaller and slower change, reaching only 15% constriction by the end of a full 60 seconds of stimulation, confirming that the rapid constriction response was driven by ChR2 activation (**Fig 4a**). No significant diameter change was observed with stimulation of mural cells on pial venules in either ChR2-YFP or control animals (**Fig. 4d,e**).

**Figure 4:**
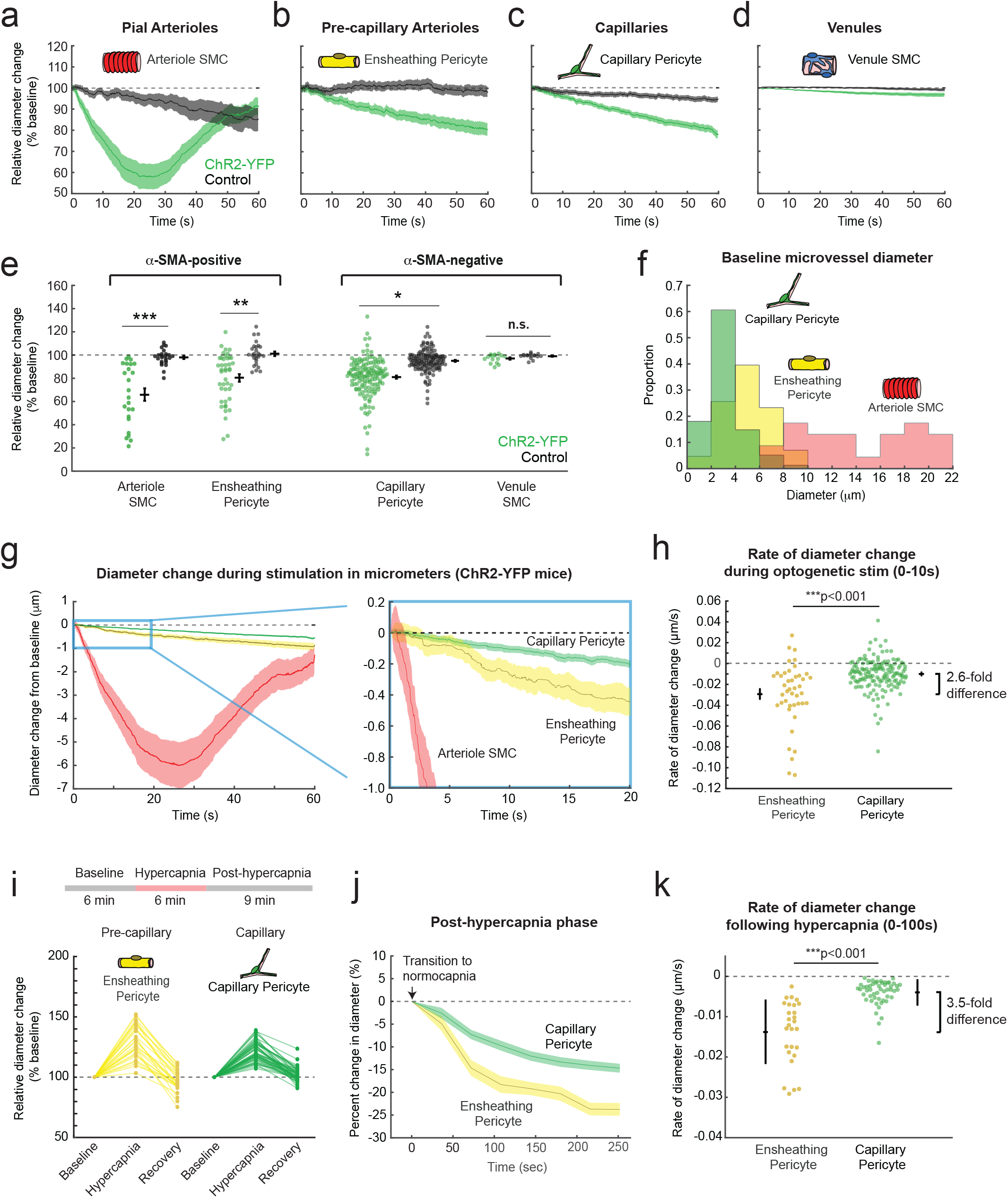
Mural cell types of different microvascular zones have distinct effects on vessel diameter. **(a-d)** Relative change in diameter over time for surface arterioles (a), pre-capillary arterioles (b), capillaries (c), and surface venules (d). Vessel types were identified based on pial angioarchitecture, sub-surface vessel branch order, and morphological identification of mural cells. All vessels were then line-scanned as described in Fig. 2b. N = 25 (5 mice) vs. 26 (4 mice) arterioles, 43 (10 mice) vs. 33 (9 mice) pre-capillary arterioles, 155 (10 mice) vs. 145 (9 mice) capillaries, 15 (4 mice) vs. 16 (3 mice) venules, for ChR2-YFP vs Control respectively. **(e)** Bee swarm plots showing relative diameter change in ChR2-YFP vessels, compared with diameter change in the same vessel type from control animals. The peak constriction of arterioles occurred at ~25 seconds of stimulation and was used in analysis; all other vessel diameters were measured at 60 seconds of stimulation and compared to baseline. Two-way repeated measures ANOVA showed overall effect and interaction of vessel type and genotype. For vessel type F(3,429)=16.51, overall p<0.0001; for genotype F(1,429)=21.53, p<0.0001, interaction F(3,429)=11.92, p<0.0001. Tukey *post hoc* analyses comparing genotypes at each branch order showed ***p<0.001, **p<0.005, *p<0.03, and n.s is p>0.1. N values same as in (a). **(f)** Histogram showing normalized distribution of baseline diameters for different vessel types in ChR2-YFP mice. N = 155 (10 mice), 43 (10 mice), 25 (5 mice), for capillaries (green), pre-capillary arterioles (yellow), and arterioles (red), respectively. **(g)** *Left:* Absolute diameter change from baseline for capillaries (green), pre-capillary arterioles (yellow), and arterioles (red) during optogenetic stimulation of vessels in ChR2-YFP mice. *Right*: Magnified view of differences in constriction between penetrating arterioles, pre-capillary arterioles and capillaries. **(h)** Bee swarm plot showing rate of diameter change during first 10 seconds of optogenetic stimulation of pre-capillary arterioles (covered by ensheathing pericytes) versus capillaries (covered by capillary pericytes). Repeated measures ANOVA showed F(1,184), p<0.001, n=43 (9 mice) pre-capillary arterioles and 155 (10 mice) capillaries. **(i)** *Top:* Experimental time-course for hypercapnia studies. *Bottom:* Relative change in pre-capillary arteriole and capillary diameter at the end of the hypercapnic phase (hypercapnia), and 9 min after return to normocapnia (recovery). Each line represents one vessel over 6 PDGFRβ-tdTomato mice. **(j)** Change in diameter for each vessel type following transition from hypercapnia to normocapnia. **(k)** The rate of absolute diameter decrease was significantly different between pre-capillary arterioles and capillaries, p<0.001 by linear mixed effects modeling (n=51 capillaries, n=27 pre-capillary arterioles in 5 mice). All data shown are mean ± S.E.M.

Below the pial surface, ensheathing pericytes and capillary pericytes contract when ChR2 is stimulated (**Fig 4b,c,e**), but were both slower relative to SMCs of pial arterioles (**Fig. 4b,c**). When comparing relative diameter change after 60 seconds of stimulation, there appeared to be no substantive difference between ensheathing and capillary pericytes within the ChR2 animals (**Fig 4e**), despite clear differences in α-SMA content (**Fig. 4e**). However, since pre-capillary arterioles were slightly larger than capillaries at baseline (**Fig. 4f**), we also compared absolute rate changes in vessel diameter during optogenetic stimulation. This revealed that stimulation of ensheathing pericytes led to ~2.6-fold faster initial contractions than did similar activation of capillary pericytes (**Fig. 4g,h**). Of note, we targeted cells from 1^st^ to 3^rd^ branch orders with ensheathing morphology, *i.e.,* circumferential processes, because this morphology further increases the likelihood of α-SMA expression.^5^

A recent study used specialized fixation procedures to show that α-SMA expression extends further into the capillary bed than previously believed in retinal vasculature.^50^ This raised the possibility that α-SMA terminations could also occur at higher branch orders in cerebral cortex. We examined this further by plotting capillary responses across all branch orders within the capillary group. This revealed similar magnitudes of constriction and reduction in RBC flow among all branch orders during optogenetic stimulation (**Supplementary Fig S14a-d)**. We sub-grouped these data for statistical analysis and found no difference between responses in 5^th^ to 7^th^ order capillaries versus 8^th^ to 9^th^ order capillaries (**Supplementary Fig S14e-g**).

On average, constriction appeared nearly linear through the 60 second stimulation for pre-capillary arterioles and capillaries (**Fig 4b,c**). However, a subset of vessels had reached a maximum constriction prior to the end of the scan, as evident from a plateau in diameter (**Supplementary Fig 15a-d**). Capillaries that plateaued did not differ from those that continued to constrict in any vasodynamic metric examined (**Supplementary Fig 15e-g**). We calculated the time to half-maximum constriction for this vessel subset, and observed a graded increase from arteriolar SMCs to ensheathing pericytes to capillary pericytes (**Supplementary Fig 16a-d**). As expected, the contraction magnitude across mural cell types exhibited a graded decrease going from arterioles to pre-capillary arterioles to capillaries (**Supplementary Fig 16e**). Collectively, these data indicate that capillary pericytes can contract *in vivo*, but with slower dynamics compared to the α-SMA-expressing mural cells of pre-capillary and pial arterioles.

### Capillaries are slower to contract back to baseline following hypercapnic dilation

To investigate the constrictive rate of microvessels in a more physiologic context, we measured the diameter of pre-capillary arterioles and capillaries following transient hypercapnic challenge (5% CO_2_ in air for 6 minutes). Hypercapnia induced robust and reliable dilation of both capillaries and pre-capillary arterioles, and constriction to baseline upon reinstatement of normocapnia (**Fig. 4i, Supplementary Fig. S17 and Supplementary Movie 3**).^51, 52, 53^ The rate of constriction to baseline after hypercapnia was significantly slower in capillaries than in pre-capillary arterioles, particularly in the first 100 seconds after transition to normocapnia (**Fig. 4j**). It is worth noting that the difference in constrictive rate between these microvascular zones was similar between optogenetic stimulation and recovery after hypercapnia. That is, pre-capillary arterioles constricted with a ~2.5-fold and 3.5-fold faster rate than capillaries during ChR2 activation and after hypercapnia, respectively (**Fig. 4k**). While this experiment clearly shows that capillaries constrict at a slower rate than pre-capillary arterioles, the experiment is correlative because upstream arteries and arterioles are also dilated by hypercapnia. However, when interpreted in the context of our optogenetic studies, it provides strong evidence that there are differences in kinetics of ensheathing pericytes and capillary pericytes during physiologically-relevant blood flow modulation.

### Pre-capillary arterioles dilate over seconds, but capillaries dilate over minutes when recovering from ChR2 activation

Intravascular pressure competes against the process of vasoconstriction, but facilitates the process of vasodilation. The kinetics of dilation may therefore be quite distinct from constriction. To further investigate this idea, we imaged the relaxation of pre-capillary arterioles or capillaries following optogenetically-induced constriction. We used 2-D movie scans to gain a better view of the constrictive and dilatory process. Capillary fields contained no pre-capillary arterioles to ensure no indirect influence on upstream flow. We again found that 800 nm laser light stimulated ChR2 and induced contraction of both ensheathing and capillary pericytes (**Fig. 5c**), which is similar to our results obtained from line scanning at this same wavelength (**Figs. 2, 4**). After 60 seconds of optogenetic stimulation, we then switched the imaging to 900 nm, which does not strongly activate mural cell ChR2 **(Supplementary Fig. 3**).^7^ Pre-capillary arterioles exhibited a rapid phase of dilation, returning to baseline within 50 seconds, on average, and ultimately settling just above baseline diameters (**Fig. 5a,c,e,f**). In some cases when maximal constriction had been achieved in pre-capillary arterioles, the pre-capillary lumen opened first proximal to the penetrating arteriole and then gradually along the vessel length, suggesting influence of upstream intravascular pressure (**Fig. 5a and Supplementary Movie 4**). In one instance, we observed delayed dilation in a distal region of the pre-capillary arteriole, despite rapid dilation upstream, suggesting the presence of a sphincter (**Supplementary Movie 5**). In contrast, capillaries dilated considerably slower, requiring on average ~300 seconds to regain baseline levels (**Fig. 5b,c,e,f, and Supplementary Movie 6,7**). When considering absolute diameter, this corresponds to a 25-fold faster rate of dilation in pre-capillary arterioles, compared to capillaries. This slow dilatory response was seen in all capillaries examined and not just a subset of cells with blebbing. Control mice expressing only YFP showed no vasoconstriction during imaging with 800 nm light (**Fig. 5d, and Supplementary Movie 8,9**). However, prolonged scanning of ensheathing pericytes in YFP mice led to a slight vasodilation suggesting that part of the dilatory response in ChR2-YFP mice was driven by light itself.^54^ In contrast, the laser light had no effect on capillary diameter in either group.

**Figure 5:**
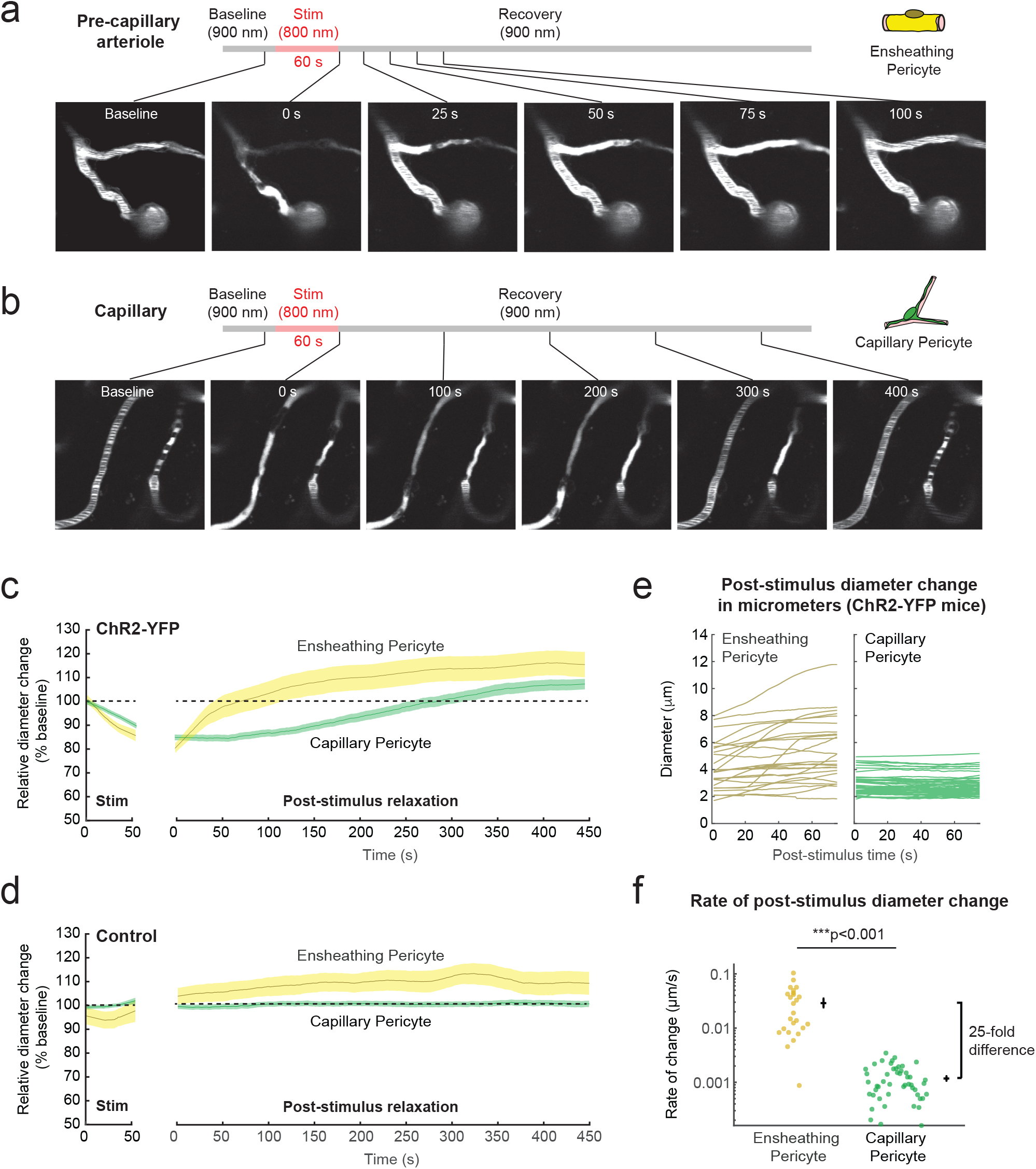
Dilation back to baseline after ChR2-mediated vasoconstriction is faster in pre-capillary arterioles than in capillaries. **(a)** Pre-capillary arterioles before and after activation of ensheathing pericyte with 800 nm in a PDGFRβ-ChR2-YFP mouse. Imaging with 800 nm excitation produces substantial constriction and cessation of RBC flow by 60 seconds of stimulation. Rapid vessel relaxation can then be observed when imaging with 900 nm light, which does not activate ChR2 to the same extent. **(b)** Same as (a), but with selective excitation of capillary pericytes. Note the slower relaxation to baseline for capillaries. **(c)** Relative change in pre-capillary arteriole and capillary diameter for stimulation (800 nm) and post-stimulus relaxation (900 nm) phases in PDGFRβ-ChR2-YFP mice. N=20 ensheathing pericytes, 49 capillary pericytes stimulated across 6 mice. Equivalent laser powers were applied for both 800 nm and 900 nm imaging. **(d)** Relative change in vessel diameter when imaging control PDGFRβ-YFP mice with the same laser power and cortical depth. N=13 ensheathing pericytes, 61 capillary pericytes irradiated across 3 PDGFRβ-YFP mice. **(e)** Absolute vessel diameters of pre-capillary arterioles and capillaries, covered by ensheathing and capillary pericytes respectively, in the first 80 seconds of post-stimulus relaxation in ChR2-expressing mice. **(f)** Rate of absolute diameter change after ChR2 stimulation is significantly different after excitation of ensheathing pericytes (yellow) versus capillary pericytes (green). N=20 ensheathing pericytes, 49 capillary pericytes across 6 PDGFRβ-ChR2-YFP mice. F(72)=197.73, p<0.001 by a linear mixed effects modeling. All time-course data shown as mean ± S.E.M.

Thus, relaxation of ensheathing pericytes combined with high intravascular pressure at the pre-capillary arteriole enables rapid vasodilation, which is not observed at the capillary level. This result is in line with recent studies showing that ensheathing pericytes on low order vessels are an early and robust responder to vasoactive stimuli and neural activity.^12, 14, 15, 38^

### Selective optical ablation of capillary pericytes increases capillary flow

Having established that contraction of capillary pericytes is sufficient to reduce capillary flow on a slow time-scale, we hypothesized that a physiological role of capillary pericytes is to maintain long-term basal capillary tone and blood flow resistance. This is likely important for establishing basal set-points of blood flow, and could occur independent of the rapid vessel diameter changes necessary for neurovascular coupling. To test this, we used two-photon optical ablation to eliminate individual pericytes in mouse cortex, as described in our previous work.^55^ In this procedure, precise line-scans with a shorter wavelength and higher power than routine imaging were used to specifically irradiate the somata of pericytes protruding laterally from the capillary wall, killing the cell by photo- and thermal toxicity (**Fig. 6a**). Death of the targeted pericyte was confirmed by loss of cellular fluorescence which persisted in the days after the ablation; cells regaining fluorescence were omitted from the data set. As a control for the effects of laser irradiation itself, we implemented a sham control where line-scans were placed with similar proximity to the capillary wall but not overlying any pericyte somata (**Fig. 6b**). All other parameters of the sham irradiation, including laser intensity, wavelength, and scan duration, matched the ablation conditions.

**Figure 6.**
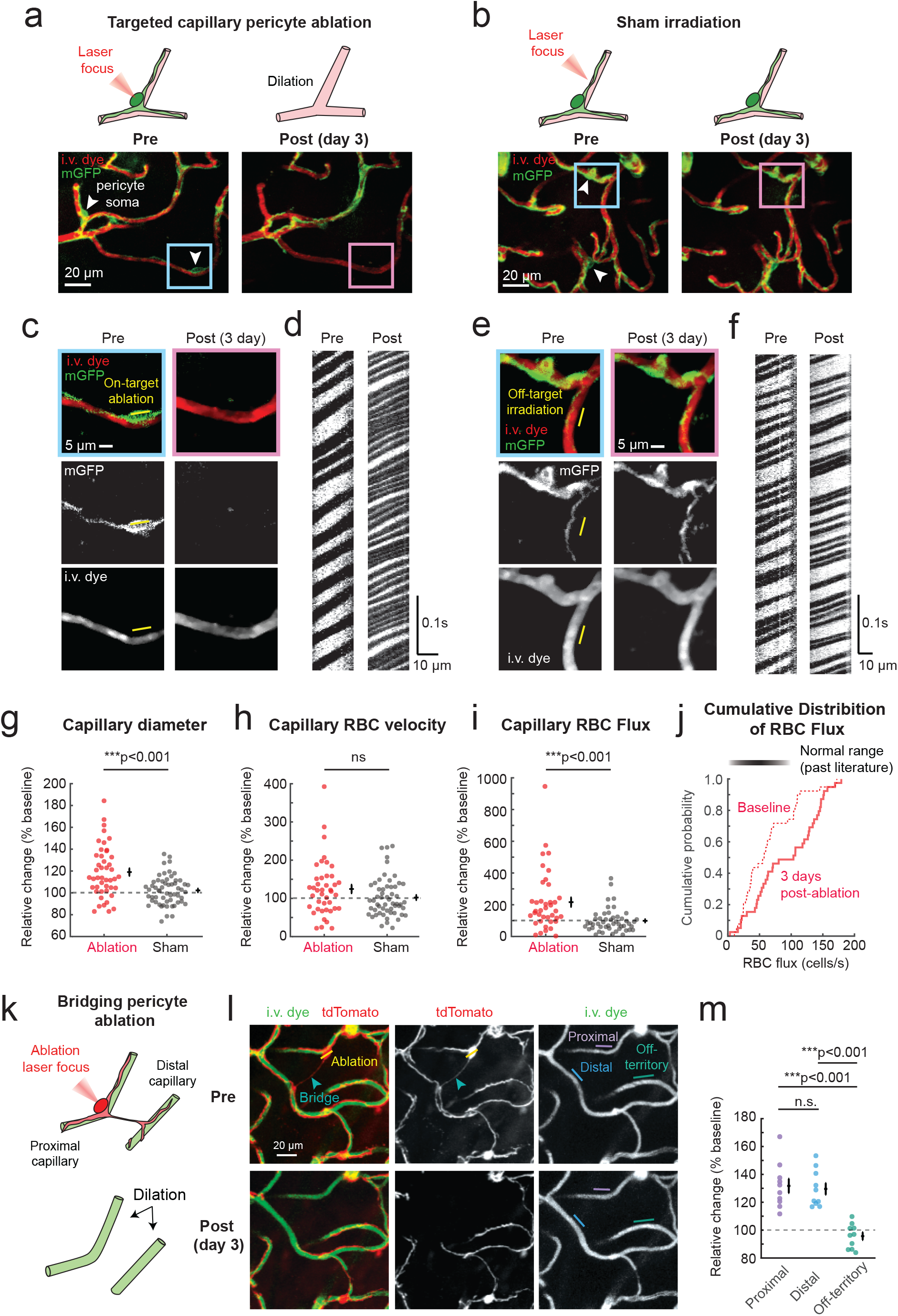
Ablation of individual capillary pericytes produces local capillary dilation and increased blood cell flux. **(a)** Two capillary pericytes were ablated in a PDGFRβ-mT/mG mouse by focusing a 725 nm line-scan only on the pericyte soma. White arrowheads mark the somata of the targeted cells. The same area of the capillary bed was then re-imaged 3 days post-ablation, which showed regions devoid of mGFP fluorescence where the ablated pericytes once covered. **(b)** A sham control involving identical scan parameters as for cell ablation. The location of the laser focus is away from the pericyte soma, but of similar distance to the capillary wall as in ablation conditions. **(c,d)** Magnified image of the boxed region in panel (a), showing the precise location of laser irradiation during ablative scanning (yellow line). Line-scans (900 nm excitation) were used to measure RBC velocity and flux from capillary segments affected by pericyte ablation. Note the marked increase in RBC flux 3 days after pericyte loss. **(e,f)** Magnified image of boxed region in sham irradiation example (panel b), and corresponding line-scan to sample RBC flow. **(g-i)** Fold change in capillary diameter (g), RBC velocity (h), and RBC flux (i) 3 days following ablation of a capillary pericyte, compared to outcome after sham irradiations in the same mice. For panel (g), F(1,95)=21.46, p < 0.001 by repeated measures ANOVA; n = 45 (6 mice) ablation and 57 (6 mice) control capillaries. For panel (h), p > 0.2 by repeated measures ANOVA; n = 41 (6 mice) ablation and 54 (6 mice) control capillaries. For panel (i), F(1,81)=9.79, p = 0.001 by repeated measures ANOVA; n = 39 (6 mice) ablation and 49 (6 mice) control capillaries. **(j)** Cumulative distribution plot of RBC flux at baseline and 3-days post-ablation. A typical range for capillary RBC flux based on past literature is provided for comparison (see text for references). **(k)** Schematic showing the ablation of a bridging pericyte to disentangle the potential effects of local light and heat damage from the effects of losing pericyte-endothelial contact. **(i)** *Top:* A pericyte targeted for ablation (yellow line across soma) with bridging processes reaching across the parenchyma to contact a distant vessel (arrowhead). *Bottom:* After ablation, pericyte contact is lost on two vessels, termed “proximal” for its proximity to the ablated soma, and the “distal” vessel. Note dilation of the proximal and distal, but not the off-territory vessel, which does not lose pericyte contact. **(m)** Dilation of both proximal and distal capillaries is a consequence of bridging pericyte ablation, but not off-territory capillaries. Kruskal-Wallis test: X^2^(2)=19.45 overall, with adjusted p values shown after Tukey post hoc test. n = 10 bridging pericyte ablations across 7 PDGFRβ-tdTomato animals.

When capillary diameter was re-examined 3 days after pericyte ablation, we observed sustained dilation averaging ~20% over baseline, with some vessels showing up to 80% dilation, which corresponds with a ~2 μm increase in absolute diameter (**Fig. 6c,g**). While this local dilation was not associated with a significant increase in RBC velocity (**Fig. 6d, h**), it produced a 2-fold increase of RBC flux compared to the sham group (**Fig. 6d, i**), as predicted by recent capillary flow modeling studies.^47^ In contrast, we found no change in diameter, RBC velocity or RBC flux over time for the sham irradiation group (**Fig. 6e-i**). Interestingly, our post-ablation flux values exceeded the RBC flux ranges found in previous studies of normal isoflurane-anesthetized mouse cortex (~45 cells/s^56^ to 62 cells/s^57^ on average). In one study, only ~10% of the capillaries exceeded RBC flux rates of 100 cell/s^57^, whereas ~50% of our post-ablation capillary population exceeded 100 cells/s, suggesting that abnormally high rates of RBC flux occur with acute pericyte loss (**Fig. 6j**).

To further ensure that the observed dilations were not the result of unintended damage to the surrounding neurovascular structures (astrocyte endfeet, endothelium), we ablated ‘bridging’ pericytes that extend their processes across the parenchyma to contact more distant capillary segments (**Fig 6k,l,m, Supplementary Fig. 18a**). These pericytes are thought to be remnants of developmental vascular pruning.^58^ Ablation of a bridging pericyte soma produced dilation of capillary segments adjacent to the ablated pericyte soma (“proximal” segment), as well as segments contacted by the pericyte bridge far from the site of irradiation (“distal” segment). Importantly, segments still contacted by neighboring pericytes did not change in diameter (“off-territory” segment), even though these segments were of similar distance from the ablation site as the distal segments (**Fig 6l,m, Supplementary Fig. 18b**). In some cases, a bridging pericyte soma was located within the parenchyma, tethered between two primary processes that extended to contact two capillaries. Ablation of these pericytes also led to dilation of both capillaries contacted by the ablated cell’s processes (**Supplementary Fig. 18c**). These data strongly suggest that loss of pericyte-endothelial contact is the reason for capillary dilation, and that laser damage or damage to other cells is unlikely to be the basis for dilation.

In the acute time-frame post-ablation, small but consistent lumen dilations could be measured within minutes following capillary pericyte ablation, suggesting immediate loss of capillary tone as opposed to long-term endothelial remodeling (**Supplementary Fig. 19**). In the chronic time-frame, neighboring pericytes extended their processes to re-cover the exposed endothelium, allowing capillaries to regain their normal diameter, consistent with our prior studies (**Supplementary Fig 20**).^55^ No extravasation of a 70kDa fluorescent dextran dye was observed after pericyte ablation, again consistent with our past work showing no BBB leak after single pericyte ablation.^55^

Collectively, these data support the idea that capillary pericytes actively maintain flow resistance throughout the capillary network, and that loss of pericytes leads to local capillary dilation and augmentation of RBC flux.

### Greater dilation of capillaries with smaller baseline diameter after pericyte ablation

Brain capillaries are heterogeneous in diameter and flow under normal basal conditions.^59, 60^ This heterogeneity is thought to create a blood flow “reserve” by providing room for the homogenization of flow across the capillary bed during functional hyperemia, and the augmentation of tissue oxygen extraction.^57, 61, 62^ Consistent with this idea, we observed that capillary diameter and blood cell flux were highly heterogeneous, even within small regions of tissue (**Fig. 7a-d**). Interestingly, these heterogeneous hemodynamics were quite stable over a 3-day interval (**Fig 7b-d**). To better understand the origins of this stable heterogeneity, we first asked whether capillary diameter influenced blood flow at baseline. Indeed, we found a weak but significant positive correlation between capillary diameter and RBC velocity and flux, in line with a previous study (**Supplementary Fig. 21a,b**).^63^

**Figure 7.**
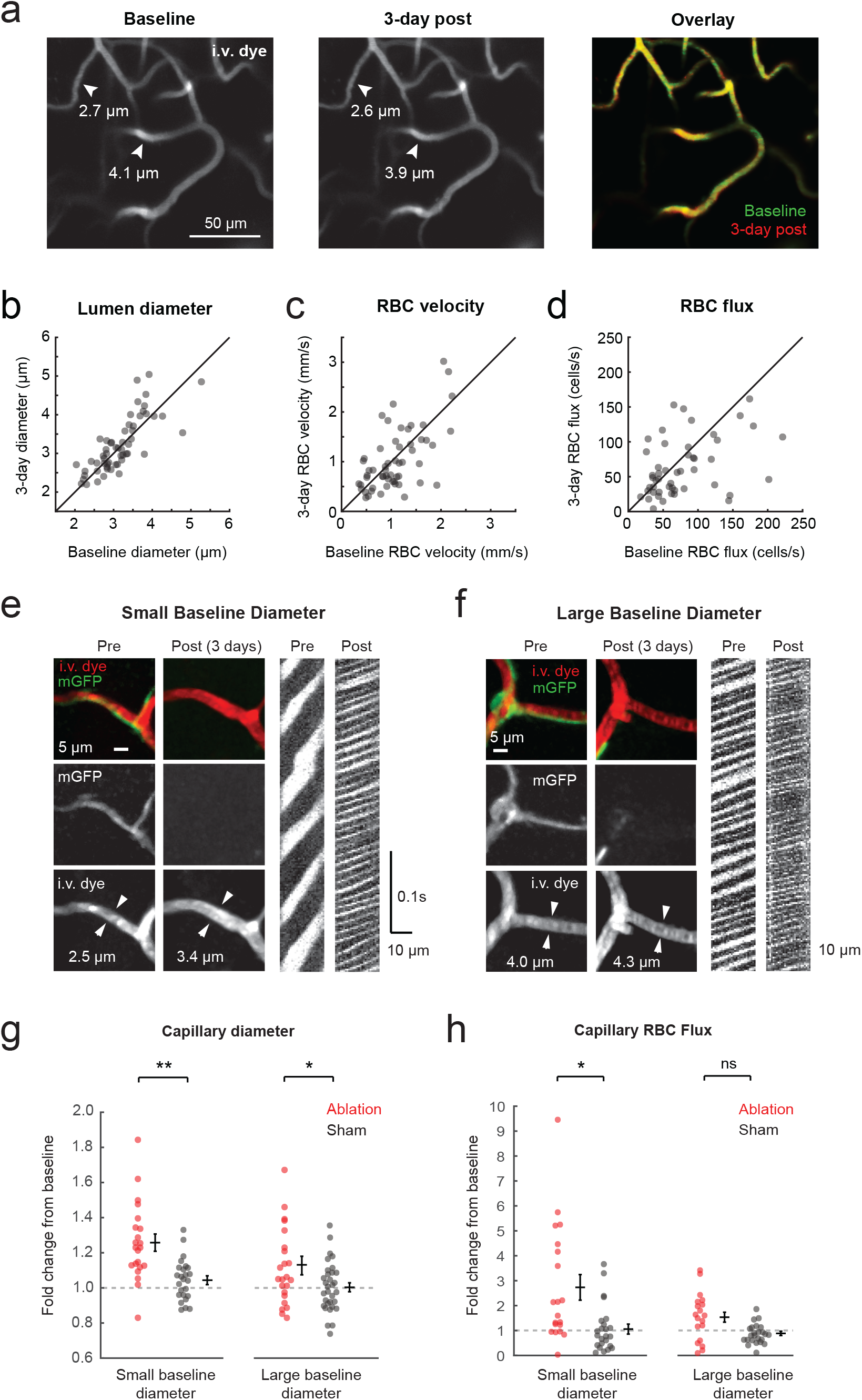
Heterogeneity of basal capillary vasodynamics. **(a)** Images of i.v. dye collected 3 days apart from the same cortical region, showing stability of capillary diameters. The image highlights one small diameter and one larger diameter capillary. Overlay images are co-registered. **(b-d)** Scatterplots and correlation analysis of capillary vasodynamic parameters collected at baseline vs. 3 days post showing that flow is relatively stable over time for the sham irradiation condition (such as Fig 6b). Pearson’s correlation: Lumen diameter: R^2^ = 0.62, p<0.001; n = 57 capillaries, RBC velocity: R^2^ = 0.39, p<0.001; n = 56 capillaries, and RBC flux: R^2^ = 0.17, p<0.01; n = 49 capillaries. **(e,f)** Example images and line-scan data collected before and 3 days following capillary pericyte ablation. A small baseline diameter capillary (e) and large baseline diameter capillary (f) is shown. **(g,h)** Bee swarm plots showing fold change in capillary diameter (g) and blood cell flux (h) in small and large diameter capillary groups. For (g), **p=0.0015, F(1,44)=11.51; * p=0.0160, F(1,44)=6.28. For (h), * p=0.0124, F(1,42)=6.83; and n.s. indicates p>0.1 by repeated measures ANOVA.

If pericytes are involved in creating this heterogeneous landscape of capillary blood flow, one would expect that some vessels are thin because pericytes are consistently exerting high amounts of tone on them, and other vessels are dilated because their associated pericytes are relatively relaxed. In line with this idea, we found that ablation of pericytes on small-diameter vessels tended to produce a greater dilation than did ablation of pericytes on larger-diameter vessels (**Fig. 7e,f, Supplementary Fig. 21c**). To permit more rigorous analyses, we split our data set at the median diameter of 3 μm to generate "small" versus "large" baseline diameter groups (**Fig. 7g,h, Supplementary Fig. 21e**). Small (<3 μm) and large (> 3 μm) vessels both dilated significantly more after ablation when compared to sham, but small-diameter capillaries dilated 2-fold more (~24% dilation) than large-diameter capillaries (~12% dilation)(**Fig. 7g**). Further, only the small capillary group showed a significant increase in RBC flux compared to the sham group (**Fig. 7h, Supplementary Fig. 21d**). Together, these data suggest that pericytes throughout the capillary bed exert variable levels of tone, which contribute to durable capillary flow heterogeneity.

## DISCUSSION

Capillary pericytes contact more than 90% of the total vascular length in the cerebral cortex, yet their ability to regulate blood flow *in vivo* has remained elusive. Using single cell optical manipulations *in vivo*, we show that capillary pericyte tone is sufficient for modulating vessel diameter, and is necessary for establishing normal baseline diameter *in vivo* (**Fig 8)**. This bidirectional control of capillary diameter occurs with relatively slow kinetics compared to upstream α-SMA positive mural cells. Further, capillary pericytes appeared to exert variable amounts of tone, as ablation of pericytes of small diameter capillaries produced proportionally greater hemodynamic effects. Altogether, the slow contractile nature of capillary pericytes is consistent with a role in establishing basal capillary flow resistance and heterogeneity at rest. The data also supports the possibility that capillary pericytes modulate late phases of functional hyperemia, or other slower physiologic phenomena. This is in contrast to the rapid kinetics required for the initial phases of neurovascular coupling, which appears to be a role better played by ensheathing pericytes and SMCs on upstream vessels.

**Figure 8.**
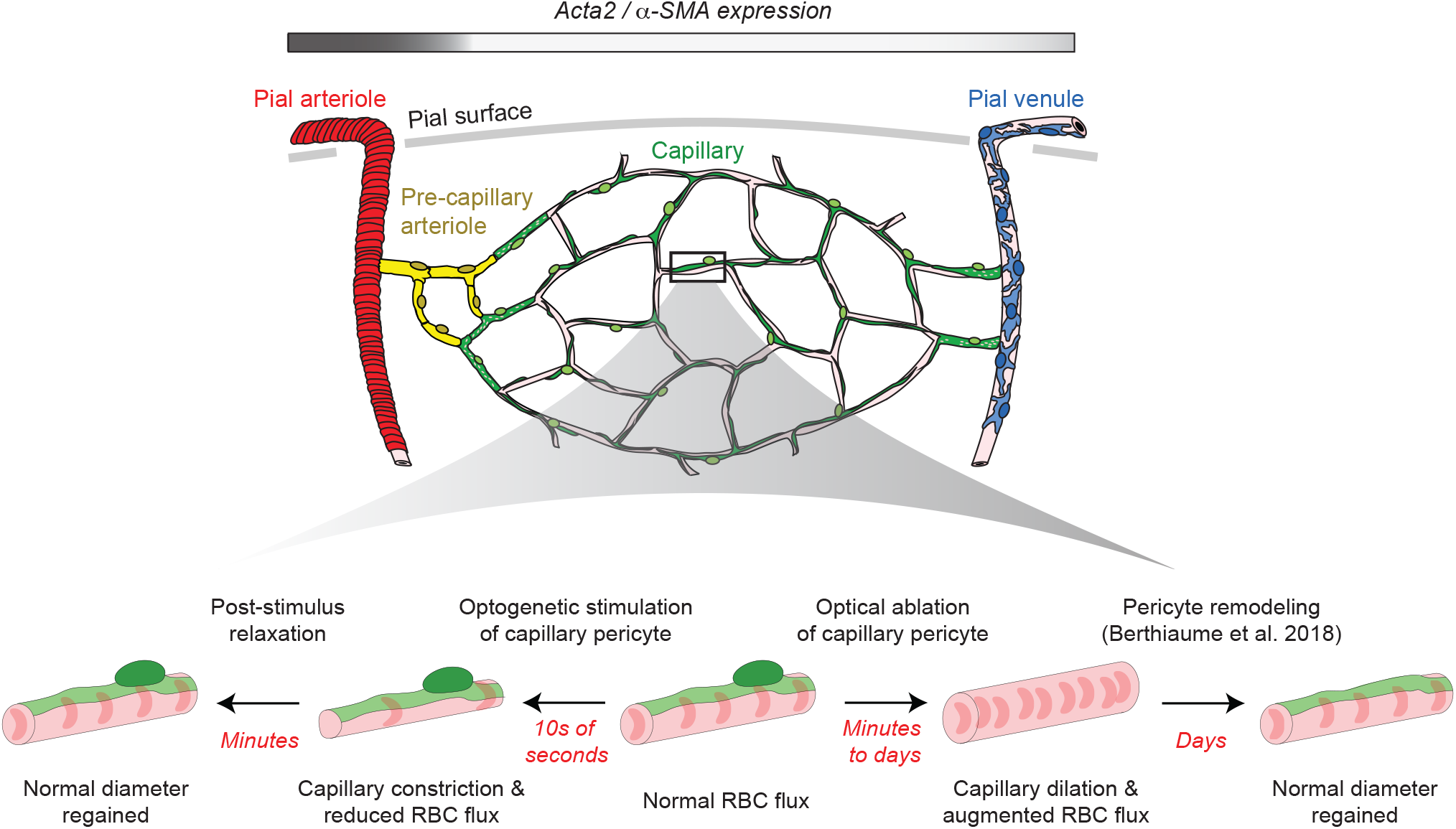
Optical cause-and-effect studies show that pericyte exert slow influence on capillary diameter in vivo. *Top:* The majority of the cerebrovasculature is ined by capillary pericytes that express little to no α-SMA. *Bottom middle:* Under basal conditions, a capillary pericyte establishes capillary tone and maintains normal RBC flux in the associated capillary. *Bottom left:* Selective activation of ChR2 in capillary pericytes leads to slow vasoconstriction and reduction of RBC flux. Slow vasodilation is observed after cessation of ChR2 activation. *Bottom right:* Selective ablation of capillary pericytes causes vasodilation and a substantial increase in RBC flux in territories left uncovered by pericytes. Diameter returns to baseline days later after neighboring pericytes regain contact with capillary regions exposed by pericyte ablation.

A past study by Hill *et al.* used a similar optogenetic approach, and concluded that capillary pericytes cannot constrict around capillaries.^7^ However, this study may have overlooked their contractile ability by examining relatively short periods of stimulation relevant to neurovascular coupling (up to 20 seconds). The laser powers used for optogenetic stimulation may also have been below the threshold needed to discern the effect of capillary pericytes. Indeed, comparative studies in non-opsin expressing mice are necessary to define the regime in which laser-induced effects dissociate from optogenetic effects, and these controls were not included in their study. Here, we challenged capillary pericytes with prolonged stimulation (up to 60 seconds) at higher laser powers, and observed the capacity for substantial capillary pericyte contraction, oftentimes involving prolonged stalls in blood flow. Importantly, the regime of laser powers we used had minimal effect on capillaries in non-opsin expressing mice, indicating that our observations are not simply the result of photo-damage or other effects of light itself.

With respect to potential contractile mechanisms, recent single cell transcriptomic studies have revealed that capillary pericytes express myosin (*myh11, myl9, myh9*) and regulators of myosin phosphorylation state (*mylk* and *Ppp1r12b*).^17^ However, they express little to no smooth muscle actin transcripts (*acta1*, *acta2, actg2, actc1*), making the binding partner and purpose of these myosin proteins unclear.^17^ It is possible that lower levels of smooth muscle actin are sufficient to support slow actomyosin contractile dynamics in capillary pericytes, and that specialized fixation procedures are necessary to detect this expression immunohistochemically.^50^ An alternate possibility is that capillary pericytes contract using a separate mechanism involving polymerization of cytoskeletal actins, a process known to be important for vascular SMC contractility.^64, 65^ In this mechanism, stimuli such as increased intravascular pressure induce the polymerization of a submembranous actin network through Rho kinase-dependent inhibition of actin depolymerization.^66, 67^ The actin cytoskeleton is tethered to focal adhesion complexes, anchoring it to the plasma membrane and creating force needed to alter cell shape. Indeed, electron microscopy (EM) in rodent brain has revealed instances where the endothelium directly underlying the pericyte becomes rippled or corrugated, as if the pericyte had pulled together only a portion of the endothelium to tighten the capillary lumen.^68^ EM studies have shown that capillary pericyte processes carry bundles of actin and myosin-like filaments that abut the endothelium, which may provide the physical substrate for force generation.^69^ Finally, the blebbing seen in our work may represent local rupture or reorganization of the actin network, caused by excessive mechanical tension when pericytes are stimulated to supraphysiologic levels, as likely occured during a subset of our optogenetic stimulation experiments.

The ability of fasudil to inhibit capillary pericyte contraction in our study has two key implications. First, as an inhibitor of Rho kinase, this drug blocks actomyosin cross-bridge cycling as well as the slower contractile mechanism of actin cytoskeleton polymerization, both of which are known to be important mechanisms in SMC contractility. In capillaries, fasudil was able to block constriction, RBC stalling, and membrane blebbing. This supports the idea that stimulation of capillary pericytes drives contractile machinery in the cell, and argues against a non-specific mechanism of capillary constriction such as pericyte swelling. Second, fasudil is clinically-used as a treatment to alleviate cerebral vasospasm after subarachnoid hemorrhage in Japan.^70^ Studies also suggest that fasudil treatment improves stroke outcomes in humans^71^ and animals.^72^ Our data indicate that one of the many salutary actions of fasudil in these diseases may be prevention of aberrant pericyte contraction and improvement of capillary blood flow.

In addition to maintenance of constant flow resistance, capillary pericytes may also contribute to slow aspects of functional hyperemia. Pioneering studies in brain slice showed that stimulation of pericytes on capillary-sized vessels elicited pericyte contraction and dilation on the time-scale of tens of seconds to minutes.^11, 73^ While the physiological relevance of these kinetics were hard to establish *ex vivo*, some recent *in vivo* studies may have identified a correlate of this slow capillary modulation. Small, delayed dilations of capillaries have been reported during sensory stimulation (1-5% increase from baseline), which can persist even after upstream vessels rapidly constrict toward baseline.^7, 38^ These capillary dilations were considered “passive” distensions resulting from upstream pressure change. However, our findings raise the possibility that capillary pericytes actively contribute to this distension. In line with this idea, capillary dilations are accompanied by decreases in pericyte calcium levels, suggesting engagement of signaling cascades within the cell.^38^ Further, a recent study showed that specific deletion of capillary pericytes in adulthood reduces functional hyperemia.^74^ In our pericyte ablation studies, average capillary dilations of 10-20% were sufficient to produce 1.5 to 3-fold increases in local RBC flux (**Fig. 7g,h**). In concert with other factors that reduce flow resistance (change in compliance of erythrocytes or leukocytes), smaller dilations of 1-5% could contribute to phasic modulation of capillary RBC flow.^16, 75^ If broadly distributed, these delayed capillary resistance changes may facilitate the even distribution of blood across cortical regions and layers after its rapid blood supply by arterioles.

Our observation that not all capillaries respond equally after pericyte ablation is consistent with a role in the creation of basal capillary flow heterogeneity.^76^ Capillary pericytes may exert different levels of tone on different capillary segments, which helps to create a mixture of low-flux and high-flux vessels within the same capillary bed. Recent studies have shown that blood flow among brain capillaries homogenizes during functional hyperemia^77^, with capillaries exhibiting low flux at rest experiencing the greatest flow increase during stimulation.^33, 57, 61, 62^ This process is thought to be important in distributing the increased blood (and oxygen) that is unleashed by the dilation of upstream arterioles.^76^ It may also be important in distributing oxygen according to basal metabolic needs, as cortical layer IV has more homogeneous flow than other layers at rest.^33^ Increased tone from a subset of capillary pericytes may thus provide a basal set-point of flow heterogeneity that balances the metabolic demands of the tissue at rest versus the demands that occur during local neural activation. In addition to playing a role in heterogeneity, the increased pericyte tone on certain vessels may be part of an overall design to prioritize blood flow to certain regions at baseline. Preferred routes for blood flow, or thoroughfare channels, through the capillary system have previously been reported in the brain^4, 78^, but their functional significance remains poorly understood. One can speculate that thoroughfare channels are shaped by regional energetic demands on protracted time scales.

Though many recent studies have detected links between pericyte dysfunction, blood flow deficits, and brain dysfunction, the mechanisms driving these relationships are unclear. Animal studies have shown that genetically-induced global pericyte loss/dysfunction leads to hypoperfusion and BBB disruption, followed by neurodegeneration.^13, 23, 27, 79^ However, the numerous deleterious effects of widespread pericyte loss make it difficult to discern exactly how the absence of capillary pericytes alters cerebral blood flow. By ablating individual pericytes in a manner that does not affect the BBB^55^, we isolated the effect of pericyte loss on local blood flow, finding that flow in fact increases in uncovered capillaries (**Fig 6**). This finding is in line with our recent study that detected increased blood flow through pre-capillary arterioles in a mouse model with mild pericyte deficiency.^53^ How might pericyte loss in chronic diseases, such as Alzheimer’s disease^80, 81, 82, 83^ and CADASIL^84^, then lead to cerebral hypoperfusion that is found in these diseases? Since RBC flux must be conserved at capillary branches, we hypothesize that pericyte loss can cause capillary “steal”, wherein flow is diverted away from active tissues down the path of least resistance, which would be the capillaries that are dilated from a lack of pericytes. In support of this idea, global but gradual pericyte loss leads to microscopic pockets of hypoxia in the cortex.^13^ Future studies could investigate this idea by examining the effects of individual pericyte ablation on flow and oxygen distribution in the broader capillary network. These abnormal dilations could co-exist within the same networks as aberrant pericyte constriction caused by amyloid β exposure and oxidative injury, further exacerbating flow patterns in capillary network.^19^

There are some limitations to the current study. First, our study focuses on mural cells, but the role of pericyte-endothelial signaling in the kinetics of basal and neurally-evoked vasodynamics is an open question.^37, 85^ Second, our imaging experiments primarily utilized isoflurane anesthesia, which is a known vasodilator. However, it is clear that capillaries were not maximally dilated by isoflurane in our preparations, as we could elicit both constrictions and dilations. Additionally, although we carefully controlled incident laser power, it is unclear if the effects of optogenetic activation were equal across the different mural cells, as we did not measure calcium influx or cellular depolarization in these studies. Finally, the topical application of fasudil dilates upstream arterioles and may have impeded pericyte contraction due to increased intravascular pressure from upstream flow. This is the inherent difficulty of using pharmacology to target specific microvascular zones, though our single cell optogenetic approach circumvented some of this concern by probing pericyte contractility one cell at a time.

In summary, we have shown that capillary pericytes are associated with the vast majority of the brain’s microvasculature, and they are necessary and sufficient to establish capillary diameter *in vivo*. Capillary pericytes exhibit slow contractile dynamics that are well-equipped to maintain basal cerebrovascular resistance, create capillary flow heterogeneity, and contribute to slow modulation of blood flow in the capillary bed.

## Abbreviations

α-SMA: α-smooth muscle actin
ACSF: Artificial Cerebral Spinal Fluid
ChR2: Channelrhodopsin-2
PDGFRβ: Platelet-derived growth factor receptor β
RBC: Red blood cell
SMC: Smooth muscle cells
YFP: Yellow fluorescent protein

## AUTHOR CONTRIBUTION

Experiments were designed by AYS, DAH, and AAB. Experiments were conducted by DAH, AAB, RIG, SAH, TN, and AYS. Data analysis was performed by DAH, AAB, SAH, TN, TT, JC, KM, and AYS. Statistics were performed by AK, AF, and DAH. The manuscript was written by AYS and DAH with contributions from all authors.

## ACKNOWLEDGMENTS

We appreciate the helpful comments and discussion of Manuel Levy, David Kleinfeld, Arthur Riegel, Prakash Kara, and Narayan Bhat.

## ONLINE METHODS

### Animals

The Institutional Animal Care and Use Committee at the Medical University of South Carolina and the Seattle Children’s Research Institute approved the procedures used in this study. The institutions have accreditation from AAALAC and all experiments were performed within its guidelines.

To express ChR2-YFP (H134R-eYFP) in mural cells for optogenetic studies, we cross-bred heterozygous PDGFRβ-Cre^29^ males (FVB/NJ background) with Ai32 females (floxed ChR2-YFP; Jax ID: 024109).^34^ To generate controls for optogenetic studies and for pericyte labeling in ablation studies, we bred heterozygous PDGFRβ-Cre males with Ai3 females (floxed YFP; Jax ID: 007903)^30^, or with mT/mG females (floxed membrane-bound eGFP; Jax ID: 007676).^86^ For SeeDB (**Fig. 1**), auxiliary ablation studies (**Fig 6k-m, Supplementary Figs. 18,19**), and hypercapnia studies (**Fig 4i-k, Supplementary Fig. 17**), we used PDGFRβ-tdTomato animals that were generated crossing heterozygous PDGFRβ-Cre males with Ai14 females (Jax ID: 007914). We identified successfully crossed mice by examining for green or red fluorescence in tail tip samples using a fluorescent stereoscope. For all studies we used a roughly equal mixture of both male and female offspring, aged 3-12 months. The mice were housed on a 12 hours light (6am-6pm) – dark cycle, with *ad libitum* access to chow and water. We did not employ randomized selection strategies for allocation of animals to different experimental groups. However, the use of ChR2-YFP and control mice, and fasudil treated and control mice, were interleaved over the entire period of study.

### Surgery

We generated acute, skull-removed cranial windows for optogenetic experiments. Under 4% isoflurane anesthesia, we first injected 50 μL of 0.06 mg/mL buprenorphine (0.1 mg/kg for a 30 g mouse) intraperitoneally. Once the animal was in the surgical plane of anesthesia (4% MAC induction, 1-2% during surgery) the scalp was excised and periosteum cleaned from the skull surface. C&B MetaBond quick adhesive cement (Parkell; S380) was then applied to the skull surface to affix a custom-made metal flange to the right half of the skull. This metal flange could later be screwed into a custom holding post for head-fixation during imaging. A 3 mm diameter circular craniotomy (dura intact) was created over the left hemisphere, and centered over 1.5 mm posterior and 3 mm lateral to bregma, which encompasses the barrel and other regions of the somatosensory cortex. The cortical surface was cleaned of any blood and covered with a drop of warm 1.5% agarose (Sigma; A9793; w/v dissolved in modified artificial cerebral spinal fluid^87^), and immediately overlaid with a 4 mm diameter glass coverslip (Warner Instruments; 64-0724 (CS-4R)). Care was taken not to compress the cortex during this process, which could affect cortical microvascular flow. MetaBond was then used to seal the edges of the coverslip, and to cover any remaining exposed skull surface or skin. We opted to use topical fasudil application, rather than systemic injection. This was done to circumvent fasudil’s inability to enter the healthy brain from the bloodstream^88^, as well as to avoid confounding reductions in cerebral perfusion pressure that can occur with systemic dosing.^89^ Fasudil was dissolved directly into the warm 1.5% agarose solution used in the cranial window procedure, providing direct access to the brain through the cortical surface. The final fasudil concentrations in the agarose (1 mM and 10 mM) were chosen based on: (1) previous studies showing that 0.1 mM fasudil prevented constriction of isolated aorta^45^, and (2) a study showing that ~10% of similarly-sized molecules at the meninges enters the brain parenchyma.^46^ This was the basis for a 1 mM fasudil concentration. We further included a 10 mM concentration to account for dilution of the fasudil by the brain interstitial fluid.^90^ Data collection began roughly 60 minutes after fasudil/vehicle loaded agarose application to allow for time to diffuse through the cortex.

For ablation experiments, we generated chronic, skull-removed windows using methods previously described.^55^ Chronic cranial windows were necessary because these experiments required multiple days of follow up imaging. This is in contrast to the optogenetic studies which could be performed acutely and required topical drug application in some studies. Chronic window implantation was similar to procedures for acute cranial windows described above, but with the following modifications. A surgical plane of anesthesia was induced with a cocktail consisting of fentanyl citrate (0.05 mg/kg), midazolam (5 mg/kg) and dexmedetomidine hydrochloride (0.5 mg/kg)(Patterson Veterinary). Dexamethasone (40 μL, Patterson Veterinary) was also given 4-6 hours prior to surgery, which helped to further reduce brain swelling during the craniotomy. Surgeries were performed under sterile conditions. Craniotomies were sealed with a glass coverslip consisting of a round 3 mm glass coverslip (Warner Instruments; 64-0720 (CS-3R)) glued to a round 4 mm coverslip (Warner Instruments; 64-0724 (CS-4R)) with UV-cured optical glue (Norland Products; 7110). The coverslip was positioned with the 3 mm side placed directly over the craniotomy, while the 4 mm coverslip laid on the skull surface at the edges of the craniotomy. An instant adhesive (Loctite Instant Adhesive 495) was carefully dispensed along the edge of the 4 mm coverslip to secure it to the skull, taking care not to allow any spillover onto the brain. Lastly, the area around the cranial window was sealed with dental cement. This two-coverslip “plug” fits precisely into the craniotomy and helps to inhibit skull regrowth, thereby preserving the optical clarity of the window over months. Imaging was initiated after a 4-week recovery period, when inflammation had subsided.

For hypercapnia experiments, we generated PoRTs windows using previously described methods.^91, 92^ This window type does not breaching the intracranial space, and therefore avoids potential herniation caused by brain volume change during hypercapnia. Briefly, the skull overlying the sensory cortex was carefully thinned to translucency using a hand-held electrical drill (Osada; EXL-M40) fitted with a 0.5 mm drill burr (Fine science tools; 19007-05). The thinned region was cleaned with ACSF and allowed to dry. A drop of Loctite 401 instant adhesive applied to the dried bone, and immediately overlayed with a 4 mm circular cover slip, taking care to avoid producing bubbles under the glass. The glue was allowed to dry for 15 minutes, and then the edges of the window were sealed with Metabond. Mice were used for hypercapnia experiment immediately after window implantation.

Distinct cranial window types were used to serve different needs in this study. To determine whether window type itself affected cerebral blood flow, we compared basal capillary RBC flux collected from acute and chronic skull-removed cranial windows. RBC flux measured between these two window types was not statistically different, with an average of 69.6 ± 34.9 cells per second for acute versus 74.9 ± 47.7 cells per second for chronic (mean ± SD; p>0.1 by Wilcoxon rank-sum test, n=87 capillaries from 9 YFP or mT/mG mice with acute windows; n=94 capillaries from 6 YFP or mT/mG mice with chronic windows).

### Two-photon imaging

#### Optogenetic experiments

Mice were imaged immediately after cranial window construction for optogenetic studies. To label the vasculature, 50 μL of 2.5% (w/v in saline) 70 kDa Texas red-dextran (Invitrogen; D1830) was injected through the retro-orbital vein under deep isoflurane anesthesia (2% MAC in medical air). Texas red-dextran injections were repeated as needed, roughly every 3 hours. Isoflurane was reduced to ~0.8% MAC in medical air, which leaves the mouse anesthetized but reactive to light toe pinch during imaging. The cortical microvasculature was imaged with a Sutter Moveable Objective Microscope coupled to a Coherent Ultra II Ti:Sapphire laser source. In some studies (relaxation after ChR2 activation and hypercapnia), a different two-photon imaging system was used, consisting of a Bruker Investigator coupled to a Spectra-Physics Insight X3. With both systems, green and red fluorescence emission was collected through 515/30 (Chroma ET510/50m-2P) and 615/60 (Chroma ET605/70m-2P) filters, respectively, and detected by photomultiplier tubes (PMTs). Low-resolution maps of the cranial window were first collected for navigational purposes using a 4-X (0.16 NA) objective (Olympus; UPlanSAPO). We then switched to a 20-X (1.0 NA) water-immersion objective (Olympus; XLUMPLFLN) and imaged with 900 nm excitation to collect volumetric data of individual penetrating arterioles in a way that would not activate ChR2 (lateral resolution 0.63 μm/pixel, axial resolution 1 μm). All imaging with the water-immersion lens was done with room temperature distilled water. The green channel (corresponding to YFP) was used to discern vascular branch order, and to pre-select specific pericytes for optogenetic stimulation based on the following criteria: (1) the branch order could be unambiguously discerned, (2) a considerable length of the pericyte was contained within one imaging plane to make line-scanning possible, (3) the pericyte was not downstream, or within 1 branch, of a previously-stimulated pericyte. We ordered pericyte stimulation from highest (capillaries) to lowest (pre-capillary arterioles and lastly pial arterioles and venules) branch order. Up to 15 pericytes within the domain of one penetrating arteriole were targeted for optogenetic activation. We only selected penetrating arterioles that extended a primary branch within the upper 300 μm of cortex. Some higher order capillaries could be traced to multiple penetrating arterioles as a source. In these cases, the target capillary was categorized based on the lowest branch order from any of the penetrating arterioles. For each mouse, we examined the branching network of 1 to 3 penetrating arterioles.

After selecting mural cells for optogenetic stimulation, we used 900 nm excitation (non-stimulating wavelength) to navigate to each of the cells (**Supplementary Fig. 3**). For each target cell, we first collected a ‘pre’ image stack (lateral resolution 0.25 μm/pixel, axial resolution 1 μm) and a brief movie at 900 nm. Images from this movie were then used to draw a multi-segmented line-scan path that transected the vessel five times, and bisected the vessel once. This was to acquire multiple measurements of lumen diameter and one measurement of RBC velocity, respectively, as described previously.^93^ The pixel dwell time for these line-scan segments was 1.5 μs. The line-scan was accelerated between these segments to further increase sampling rate. With these parameters we routinely achieved a ~600 Hz sampling rate. We measured the proportion of the line-scan that was fluorescent in the YFP channel, which represent regions where activation of ChR2-YFP could occur. Approximately 8% of the line-scan intersected with ChR2-YFP in capillary scans, *i.e.* 8% duty cycle, and 11% for both pre-capillary arteriole and pial arteriole scans due to greater vessel coverage of ensheathing pericytes and SMCs. This slight difference in parameters likely had no effect on outcome, as there was no correlation between duty cycle and the change in vessel diameter when examining each vessel type separately (data not shown), or with all branch orders combined (1^st^ to 9^th^) (R=−0.05, p=0.5 by Pearson’s correlation of proportion of scan with YFP vs. fold change diameter, n=197 vessels). For pial arterioles and many pre-capillary arterioles, a 600 Hz rate could not capture RBC velocity, but could be used to determine vessel diameter. During stimulation, we maintained line-scanning of the target vessel for 60 seconds at 800 nm excitation, followed by collection of a ‘post-stimulation’ stack of the same region using 900 nm excitation. Every 2 hours of imaging, we injected 200 μL of lactated ringer’s solution subcutaneously (Patterson Veterinary; 07-869-6319). At the conclusion of each experiment, mice were still reactive to a light toe pinch.

The laser powers utilized for the 800 nm and 900 nm excitation line-scans were kept consistent across experimental groups by adhering to a power-vs-cortical depth chart established from preliminary studies on 4 PDGFRβ-ChR2-YFP mice. Data collected throughout the study were then added to this data set, so that comparable levels of laser irradiation at certain depths were applied for all animals (**Supplementary Fig. 4**). Laser powers were determined at the output of the 20-X objective using a laser power meter (Thor Labs; PM100D), with galvanometric mirrors engaged in a full field, 512 × 512 pixel scan covering 16,600 μm^2^. Comparable PMT gain values were used across all animals.

For studies using chlorprothixene sedation (**Supplementary Fig. 5, Fig. 4i-k**), we injected 30 μL of 1 mg/mL chlorprothixene solution intramuscularly (thigh muscle), immediately after cranial window surgery with isoflurane (Sigma C1671). We then injected 70 kDa Texas-red dextran retro-orbitally under 2% isoflurane, turned off the isoflurane, and waited 15-30 minutes for chlorprothixene to take effect. Some mice required more than one injection per imaging session. Basal capillary diameter did not differ significantly between isoflurane and chlorprothixene sedation. However, a modest increase in RBC velocity was soon, likely due to upstream arteriole dilation (**Supplementary Fig. 5c-f**).

#### Relaxation after ChR2 activation

Using PDGFRβ-ChR2-YFP mice anesthetized with 0.8% MAC isoflurane (separately established at the Seattle Children’s Research Institute), we collected full field movies before, during and after optical stimulation. Movies were collected in regions that *(i)* contained a penetrating arteriole with pre-capillary arteriole offshoot, or *(ii)* contained only capillaries. In the latter, we carefully selected capillary regions with no penetrating or pre-capillary arterioles within the imaging field to avoid perturbing flow upstream of the imaged capillaries. During continuous movie collection, we started with imaging at 900 nm, then switched to 800 nm to stimulate ChR2 for 60 seconds, and then returned to 900 nm for an additional 450 seconds to observe vessel relaxation. Movies were collected at ~1 frame per second, and imaging resolution was 0.38 μm/pixel. Imaging was performed with an Insight X3 laser where power at the output was nearly identical between 800 nm and 900 nm. Thus, the same laser power (~50 mW) was applied for both stimulation and observation periods for vessels imaged between 50 and 150 μm below the pial surface.

#### Hypercapnia experiments

PDGFRβ-tdTomato mice were sedated with chlorprothixene to ensure full capacity for vasodilation, and imaged through a thinned-skull window. Vasculature was labeled with retro-orbital injection of 2 MDa FITC-dextran. To ensure the same vessel regions were observed despite shifting due to brain volume change, we collected continuous 25 μm thick image stacks at a sampling rate of 1 stack every 36 seconds. Lateral resolution was 0.38 μm/pixel and axial resolution of 1 μm/pixel. During continuous stack collection, we established a 6 min baseline (10 stacks) during inhalation of air, then 6 minutes of 5% CO_2_ with air balance, delivered through a nose cone placed ~5 mm from the snout. Finally, we returned to air inhalation for 9 minutes to observe post-hypercapnic constriction. Movement artifacts were common during the hypercapnic phase, precluding our ability to measure rate of vasodilation. However, the mice were calm during the post-hypercapnic phase, allowing quality measurements of vessel diameter. Only vessels without movement artifact at baseline normocapnia and during the entire recovery phase were considered for analysis. Diameters were analyzed as described below using an ImageJ macro.

#### Ablation experiments

Mice with chronic windows were anesthetized with isoflurane and 70 kDa Texas red-dextran was injected to label the vasculature. Isoflurane was maintained at 0.8% MAC during imaging. On imaging day 1, we collected volumetric data of penetrating arterioles at 900 nm excitation with the 20-X, 1.0 NA objective. We used these as maps to select pericytes for ablation. Good candidates for ablation exhibited: (1) unambiguous capillary pericyte morphology, and (2) had somata protruding abluminally that could be easily targeted by the laser without collateral damage to the capillary wall. We then collected high-resolution ‘pre-ablation’ image stacks of each target capillary (lateral resolution 0.37 μm/pixel, axial resolution 1 μm) and line-scans to gather RBC velocity and flux. After collecting baseline data, we flipped a coin to decide whether the targeted pericytes would be part of the ablation group, or sham irradiation control group. We stratified this randomization process so that ablation and control data were collected at comparable depths, and so that the same number of ablation and control cells would be collected from each animal. Further, we ensured that each cell targeted for ablation was more than two capillary branches away from other cells that were targeted for sham or ablation.

To perform pericyte ablation, we used techniques previously described.^55^ Briefly, at our highest digital zoom (10X; 0.06 μm/pixel), we applied a small line-scan restricted to only the soma of the target pericyte (1-2 μm away from the vessel wall), and scanned this region for 1-3 minutes using 725 nm excitation at 20-60 mW, depending upon cortical depth. The ablative scan was periodically interrupted to check for successful ablation, which was apparent as loss of fluorescence from the entire cell. Successful ablations resulted in prolonged absence of fluorescence from the targeted pericyte, but not in neighboring pericytes, when examined 3 days after ablation. For sham irradiation data, we used the same parameters of laser irradiation, but applied the line-scan 1-2 μm away from the vessel wall at a region lacking pericyte somata. This is the same approximate distance from the vessel wall as when targeting a pericyte soma. This procedure was similarly applied to ablation of bridging pericytes.

During post-ablation imaging, we relocated the targeted vessel using vascular angioarchitecture for guidance. Image stacks and line-scans of the same target capillary were collected with identical parameters as baseline. Importantly, the head stage of the animal remained fixed throughout all imaging sessions, so vessel orientation would not be different between days. In 4 mice, we also ablated (or sham-irradiated) 2 adjacent pericytes to examine whether the amount of uncovered capillary length influenced the extent of hemodynamic change post-ablation (**Fig 6a,b**). However, we found no difference in capillary hemodynamics whether one or two pericytes were ablated (data not shown), and the ablation data were therefore combined.

### Analysis of in vivo imaging data

#### Determining vessel depth

We calculated the cortical depth of all stimulated vessels by browsing the initial 300 μm of cortical depth in image stacks of the penetrating arteriole network using ImageJ. We measured the distance between the bottom of the dura mater and the plane in which the line-scan for a single vessel was performed. This was facilitated by YFP or GFP labeling of meningeal fibroblasts in the dura of our PDGFRβ mice.

#### Automated analysis of vessel diameter and RBC velocity from line-scan data

For each line-scan in our optogenetic studies, we calculated the diameter from all 5 diameter lines by measuring the full-width at half maximum of the vessel’s intensity profile, as previously described.^87, 93^ The spatial resolution of the diameter scans were ~ 0.5 μm/pixel. Linear interpolation was used to add subpixel accuracy to the diameter measurement. We calculated RBC velocity from the single bisecting line-scan using a radon transform method, as previously described.^94^ The window of data used for analysis of both diameter and RBC velocity was 40 ms (average of ~24 lines). Time-course data was acquired by running this window along the length of the line-scan, with 20 ms overlap between windows, providing 50 measurements per second. We then processed these raw diameter and RBC velocity values in batches using custom MATLAB scripts. During this process, we filtered out extreme RBC velocity values by converting values greater than 3-times the standard deviation into the mean of the entire trace. Then, for unbiased quality control, the mean of the diameter trace had to be greater than the standard deviation for the data to be analyzed using automation. Typically, 4 of the 5 collected diameter traces passed this criterion, and these traces were averaged together. The occasional presence of a stalled RBC prevented the ability to obtain a diameter calculation, and these traces were excluded. In instances where a “fold-change” in diameter or RBC velocity was provided, this was calculated by dividing the averaged value from 58-60 seconds in the line-scan by the average of 0-3 seconds, the latter being considered our baseline. The time course traces shown in the figures using line-scan are median filtered over 1 second, but median-filtered data were not used to calculate fold changes used in statistical comparisons. For the ablation experiments, we analyzed the RBC velocity in the same manner as for optogenetic studies, explained above. RBC velocities for ablation experiments were reported as the average of a 60 second line-scan collected at 900 nm excitation.

#### Automated analysis of vessel diameter in full field or maximally projected images

Lumen diameter was measured in full field images for ablation, post-optogenetic relaxation, and hypercapnic studies. The 2-D images or movies were anonymized to blind the rater (D.A.H., T.T., or A.Y.S.) to the experimental group. To reduce bias of measurement location, we used a custom ImageJ macro, as previously described^53^, to analyze lumen diameter at multiple, equidistant locations along each vessel segment of interest. The user first draws a line that bisects the vessel segment to be measured. The macro then creates perpendicular cross-lines (spaced ~3 μm) to obtain fluorescence intensity across the lumen width. It then calculates the full-width at half maximum of the intensity profile for each equidistant location. These values were then averaged together, and converted to micrometer units, to obtain a single diameter value per vessel segment. In cases where RBC flow measurements hade also been collected for the same vessel, the initial bisecting line was drawn to match where RBC flow line-scan measurements were collected.

#### Rate of diameter change

Continuous diameter data from individual vessels were linearly regressed in MATLAB software to calculate the rate of absolute diameter change. We focused on periods where the kinetics between ensheathing versus capillary pericytes were clearly distinct and in a roughly linear phase of change. In optogenetic constriction data, this was the first 10 seconds after initiation of optogenetic stimulation for both cell types. In hypercapnia data, this was the first ~144 seconds of data after transition to normocapnia for both cell types. In data collected for relaxation after ChR2 activation, the time range examined was different for ensheathing and capillary pericytes due to their very distinct kinetics. Rate was calculated in the first 50 seconds following the optogenetic stimulation phase for ensheathing pericytes, and from 50-400 seconds for capillary pericytes.

#### Analysis of flow-no flow and membrane blebbing in post-stacks

We visually examined all post-ablation image stacks in a blinded and randomized manner. Image stacks from all experimental groups were gathered and randomized for analysis using a random number generator within custom MATLAB code. We scrolled through each image stack and categorized them for blood flow through the stimulated capillary segment. We denoted capillaries as “flowing” if there was any movement of RBC shadows between frames of the image stack. In a blinded manner, we also tallied the presence of pericyte membrane ‘blebbing’, defined as focal, bubble-like protrusions from the stimulated pericyte that were not present prior to the stimulation period.

#### Manual analysis of RBC velocity

Automated analysis of RBC velocity was inaccurate for cases that slowed greatly or stopped flowing altogether.^94^ Therefore, whenever the standard deviation exceeded the mean RBC velocity, we manually analyzed the RBC velocity (44/175 ChR2-YFP vs. 2/161 control, for branch orders 5-9). To do this, we compared the slope of RBC streaks in one frame at the beginning of the line-scan, and one frame at the end. We reduced bias by blinding the rater (D.A.H) to the timing of the presented frames (beginning or end of scan). We calculated: (1) the distance traveled by RBCs (x-dimension) using a pixel/μm conversion for the line-scan, and (2) time of travel (y-dimension) using the line sampling rate, and divided these values to derive RBC velocity. Of note, manual RBC velocity analyses were only used in calculating fold change in order to increase accuracy for statistics. Continuous RBC velocity traces presented in the figures did not exclude any data based on high standard deviation.

#### Quantification of RBC flux

The line-scan data from optogenetic or ablation studies were cropped to show only the part of the scan where velocity was measured. We then manually counted the number of blood cell streaks occurring in frames at 1, 30, and 60 seconds of the line-scan using a custom-made MATLAB script. These frames were presented to the rater (D.A.H.) in random order, and blind to the group, to reduce bias. Some streaks appeared thicker and likely corresponded to a rouleaux of RBCs. These streaks were counted as single RBCs since it was not possible to distinguish the number of cells, which may lead to a slight underestimation of the RBC count. Line-scans were excluded from analysis if the RBC flux was too high for reliable cell counting (**Supplementary Fig. 11**), or if the RBCs were stuck together for more than ~20% of the presented frame (50/168 ChR2-YFP scans, and 61/148 YFP scans were excluded under these criterion). For optogenetic data, RBC flux counts at 30 and 60 seconds were then normalized to counts at 1 second to provide a fold-change from baseline. For pericyte ablation data, RBC flux values at 1, 30 and 60 seconds were averaged together to provide a single value for each capillary in a given day.

#### Image Processing

Only raw images were utilized in data analysis, viewed with ImageJ or MATLAB software. For presentation purposes, images were contrasted and cropped in Adobe Photoshop, one color channel at a time, in a similar manner across all conditions.

#### Exclusion critera

For optogenetic experiments, we omitted data from any capillaries with visible Texas red-dextran extravasation in the post-stimulation image stack. Diameter data were excluded if the baseline diameter was less than 1 μm. If precise depth could not be calculated for a capillary, we did not include that capillary in plotting laser power versus cortical depth. In ablation data sets, we excluded cells that were not successfully ablated, or if neighboring processes had already grown into the previously uncovered region, at 3-days post-ablation. For hypercapnia experiments, we excluded vessels that dilated less than 5% above baseline levels, as the magnitude of constriction was too small for reliable rate calculation.

### Histology

#### Thin sections

Isolated pericyte were examine in **Fig 1g,h**. As previously described^5^, NG2-Ai14 mice were deeply anesthetized with euthasol (Patterson Veterinary; 07-805-9296), and then transcardially perfused with phosphate-buffered saline (PBS), followed by 4% paraformaldehyde (PFA) in PBS. After perfusion, the brain was extracted and incubated in 4% PFA in PBS overnight. Brains were then transferred to PBS with 0.01% sodium azide, where they were stored until sliced into 100 μm sections on a Vibratome Series 1000. We performed antigen retrieval by placing sections into a vial of 0.125% trypsin (Sigma; T4049) in PBS, and incubating the vial at 37°C for 1 hour. Afterwards, FITC-conjugated tomato lectin (1:250 dilution; Vector Laboratories FL-1171) and anti-α-SMA antibody (1:200 dilution; mouse host, Sigma A5228) were added to an incubation solution composed of 2% TritonX-100 (v/v, Sigma-Aldrich; X100), 10% goat serum (v/v, Vector Laboratories; S1000), and 0.1% sodium azide (w/v, Sigma-Aldrich; S2002) in PBS. After overnight incubation, the slices were washed in PBS, and then the slices were added to incubation solution containing anti-mouse Alexa 647 secondary antibody (1:500 dilution; ThermoFisher; A31626) for 2 hours. Slices were washed in PBS, then mounted onto a slide, dried, and sealed with a cover slip using Fluoromount G (Southern Biotech). We collected images using an Olympus BX61 microscope with an UPLSAPO10×2 objective lens (0.345 pixels per μm lateral resolution, 1 μm Z step size), and UPLSAPO 60xO objective lens (8.8 pixels per μm lateral resolution, 1 μm Z step size), with laser and PMT parameters kept comparable for all cells.

#### Optical clearing and two-photon imaging of thick brain sections

To measure capillary pericyte territory in the cortical vasculature, we used the same optical clearing and imaging methods described previously.^5^ Briefly, a vibrotome was used to cut 500 μm thick coronal sections of PDGFRβ-tdTomato brains. These sections and our images encompassed the somatosensory cortex based on visible landmarks and a mouse brain atlas. Trypsin antigen retrieval was performed for 1 hour, as described for thin sections. The tissues were then incubated in 1:200 anti-α-SMA-FITC antibody (Sigma; F3777) in the same incubation solution described above for 1 week, and then rinsed in PBS for 1 h. The See Deep Brain (SeeDB) optical tissue clearing method was then performed.^31^ After the 5 day SeeDB protocol, the sections were submerged in SeeDB solution and imaged with two-photon microscopy. Four large image stacks were collected, each encompassing a volume of 0.055 mm^3^, with dimensions 500 μm in anterior-posterior direction (corresponding to coronal slice thickness), 312 μm in medial-lateral direction, and 350 μm of depth from the pial surface. Only the upper 350 μm of cortex was analyzed, as it corresponded with our *in vivo* imaging capabilities. The image stacks were stitched together with the XuvTools^95^ (www.xuvtools.org) and further examined in Imaris software (Bitplane). The total length of microvasculature was then manually traced in 3-D using Imaris, and the proportion exhibiting tdTomato or anti-α-SMA-FITC fluorescence was determined. For calculations of branch order of α-SMA termination, we analyzed the data sets as described previously, but reported α-SMA termination of each vessel branch, rather than averaging branch orders per penetrating arteriole offshoot.^5^

### Statistics

Calculations and statistical tests were performed in either MATLAB (R2018a), SAS version 9.4 or R version 3.5.1.

#### Main figures

For optogenetic studies, fold changes in diameter and RBC velocity were calculated by taking the average value from 58-60 seconds of the 800 nm scan, and dividing by the baseline, defined as the average value of the first 3 seconds of same scan. For RBC flux in optogenetic studies, the fold change was the number of cells in a frame corresponding to the 60 second timepoint, divided by the number of cells in the frame corresponding to the 1 second timepoint. Fold change values of vessel diameter, RBC velocity, and RBC flux were compared across groups using either a repeated measures analysis of variance (ANOVA) model or a linear mixed effect model (LMEM), with genotype or ablation/sham group as co-variables in the model. A random intercept was included in all models to account for differences across animals. This strategy accounts for the nested nature of the data, *i.e.,* that multiple vessels come from one animal and are thus not independent of one another. A repeated measures ANOVA was used in comparing the effect of pericyte ablation on small and large vessels. In addition, LMEM was used in comparing rate of constriction after hypercapnia and rate of vessel dilation after optogenetic stimulation on small and large vessels. A two-way repeated measures ANOVA was used when comparing relative diameter change on both vessel type and genotype. All post-hoc comparisons were made using a Tukey post hoc adjustment, where applicable. Data for the fold change in RBC velocity and flux after ablation, as well as rate of diameter change after optogenetic stimulation and hypercapnia, were right-skewed and log-transformed prior to running analyses.

#### Supplemental Figures

Details of statistical tests used are described within the figure legends.

### Data Availability Statement

Data and figures originating from this study are available upon request.

## REFERENCES

1. Kisler K, Nelson AR, Montagne A, Zlokovic BV. Cerebral blood flow regulation and neurovascular dysfunction in Alzheimer disease. Nature Review Neuroscience 18, 419–434 (2017).

2. Blinder P, Tsai PS, Kaufhold JP, Knutsen PM, Suhl H, Kleinfeld D. The cortical angiome: An interconnected vascular network with noncolumnar patterns of blood flow. Nature Neuroscience 16, 889–897 (2013).

3. Gould IG, Tsai PS, Kleinfeld D, Linninger A. The capillary bed offers the largest hemodynamic resistance to the cortical blood supply. Journal of Cerebral Blood Flow & Metabolism 37, 52–68 (2016).

4. Schmid F, Tsai PS, Kleinfeld D, Jenny P, Weber B. Depth-dependent flow and pressure characteristics in cortical microvascular networks. PloS Computational Biology 13, e1005392 (2017).

5. Grant RI, Hartmann DA, Underly RG, Berthiaume A-A, Bhat NR, Shih AY. Organizational hierarchy and structural diversity of microvascular pericytes in adult mouse cortex. Journal of Cerebral Blood Flow & Metabolism 39, 411–425 (2017).

6. Hartmann DA, Underly RG, Grant RI, Watson AN, Lindner V, Shih AY. Pericyte structure and distribution in the cerebral cortex revealed by high-resolution imaging of transgenic mice. Neurophotonics, 041402 (2015).

7. Hill RA, Tong L, Yuan P, Murikinati S, Gupta S, Grutzendler J. Regional Blood Flow in the Normal and Ischemic Brain Is Controlled by Arteriolar Smooth Muscle Cell Contractility and Not by Capillary Pericytes. Neuron 87, 95–110 (2015).

8. Armulik A, Genove G, Betsholtz C. Pericytes: Developmental, physiological, and pathological perspectives, problems and promises. Developmental Cell 21, 193–215 (2011).

9. Rouget C. Memoire sur le developpement la structure et les proprietes physiologiques des capillaires sanguins et lymphatiques. Arch Physiol Norm Path 5, 603–663. (1873).

10. Eberth CJ. Handbuch der Lehre von der Gewegen des Menschen und der Tiere Leipzig: Engelmann (1871).

11. Peppiatt CM, Howarth C, Mobbs P, Attwell D. Bidirectional control of CNS capillary diameter by pericytes. Nature 443, 642–643 (2006).

12. Hall CN, et al. Capillary pericytes regulate cerebral blood flow in health and disease. Nature 508, 55–60 (2014).

13. Kisler K, et al. Pericyte degeneration leads to neurovascular uncoupling and limits oxygen supply to brain. Nature Neuroscience 20, 406–416 (2017).

14. Cai C, et al. Stimulation-induced increases in cerebral blood flow and local capillary vasoconstriction depend on conducted vascular responses. Proceedings of the National Academy of Sciences 15, E5796–E5804 (2018).

15. Fernández-Klett F, Offenhauser N, Dirnagl U, Priller J, Lindauer U. Pericytes in capillaries are contractile *in vivo*, but arterioles mediate functional hyperemia in the mouse brain. Proceedings of the National Academy of Sciences USA 107, 22290–22295 (2010).

16. Wei HS, et al. Erythrocytes Are Oxygen-Sensing Regulators of the Cerebral Microcirculation. Neuron 91, 851–862 (2016).

17. Vanlandewijck M, et al. A molecular atlas of cell types and zonation in the brain vasculature. Nature 554, 475–480 (2018).

18. Mishra A, Reynolds JP, Chen Y, Gourine AV, Rusakov DA, Attwell D. Astrocytes mediate neurovascular signaling to capillary pericytes but not to arterioles. Nature Neuroscience 19, 1619–1627 (2016).

19. Nortley R, et al. Amyloid β oligomers constrict human capillaries in Alzheimer’s disease via signaling to pericytes. Science 365, Epub 2019 Jun 2020 (2019).

20. Yemisci M, Gursoy-Ozdemir Y, Vural A, Can A, Topalkara K, Dalkara T. Pericyte contraction induced by oxidative-nitrative stress impairs capillary reflow despite successful opening of an occluded cerebral artery. Nature Medicine 15, 1031–1037 (2009).

21. Fernández-Klett F, et al. Early loss of pericytes and perivascular stromal cell-induced scar formation after stroke. Journal of Cerebral Blood Flow & Metabolism 33, 428–439 (2013).

22. Montagne A, et al. Pericyte degeneration causes white matter dysfunction in the mouse central nervous system. Nature Medicine 24, 326–337 (2018).

23. Sagare AP, et al. Pericyte loss influences Alzheimer-like neurodegeneration in mice. Nature Communications 4, 2932 (2013).

24. Winkler EA, Sagare AP, Zlokovic BV. The pericyte: a forgotten cell type with important implications for Alzheimer’s disease? Brain Pathology 24, 371–386 (2014).

25. Arango-Lievano M, et al. Topographic Reorganization of Cerebrovascular Mural Cells under Seizure Conditions. Cell reports 23, 1045–1059 (2018).

26. Lendahl U, Nilsson P, Betsholtz C. Emerging links between cerebrovascular and neurodegenerative diseases-a special role for pericytes. EMBO Reports 20, e48070 (2019).

27. Nikolakopoulou AM, et al. Pericyte loss leads to circulatory failure and pleiotrophin depletion causing neuron loss. Nature Neuroscience 22, 1089–1098 (2019).

28. Taylor ZJ, et al. Microvascular basis for growth of small infarcts following occlusion of single penetrating arterioles in mouse cortex. Journal of Cerebral Blood Flow & Metabolism 36, 1357–1373 (2016).

29. Cuttler AS, et al. Characterization of Pdgfrb-Cre transgenic mice reveals reduction of ROSA26 reporter activity in remodeling arteries. Genesis 49, 673–680 (2011).

30. Madisen L, et al. A robust and high-throughput Cre reporting and characterization system for the whole mouse brain. Nature Neuroscience 13, 133–140 (2010).

31. Ke MT, Fujimoto S, Imai T. SeeDB: a simple and morphology-preserving optical clearing agent for neuronal circuit reconstruction. Nature Neuroscience 16, 1154–1161 (2013).

32. Sakadžić S, et al. Large arteriolar component of oxygen delivery implies a safe margin of oxygen supply to cerebral tissue. Nature Communications 5, 5734 (2014).

33. Li B, et al. More homogeneous capillary flow and oxygenation in deeper cortical layers correlate with increased oxygen extraction. Elife 8, e42299 (2019).

34. Madisen L, et al. A toolbox of Cre-dependent optogenetic transgenic mice for light-induced activation and silencing. Nature Neuroscience 15, 793–802 (2012).

35. Mathiisen TM, Lehre KP, Danbolt NC, Ottersen OP. The perivascular astroglial sheath provides a complete covering of the brain microvessels: An electron microscopic 3D reconstruction. Glia 9, 1094–1103 (2010).

36. Haley MJ, Lawrence CB. The blood-brain barrier after stroke: Structural studies and the role of transcytotic vesicles. Journal of Cerebral Blood Flow & Metabolism 37, 456–470 (2017).

37. Longden TA, et al. Capillary K+-sensing initiates retrograde hyperpolarization to increase local cerebral blood flow. Nature Neuroscience 20, 717–726 (2017).

38. Rungta RL, Chaigneau E, Osmanski BF, Charpak S. Vascular Compartmentalization of Functional Hyperemia from the Synapse to the Pia. Neuron 99, 362–375 (2018).

39. Underly RG, Levy M, Hartmann DA, Grant RI, Watson AN, Shih AY. Pericytes as inducers of rapid, matrix metalloproteinase-9 dependent capillary damage during ischemia. Journal of Neuroscience 37, 129–140 (2017).

40. Neuhaus AA, Couch Y, Sutherland BA, Buchan AM. Novel method to study pericyte contractility and responses to ischaemia in vitro using electrical impedance. Journal of Cerebral Blood Flow & Metabolism 37, 2013–2024 (2016).

41. Paluch E, Sykes C, Prost J, Bornens M. Dynamic modes of the cortical actomyosin gel during cell locomotion and division. Trends in Cell Biology 16, 5–10 (2006).

42. Somlyo AP, Somlyo AV. Signal transduction and regulation in smooth muscle. Nature 372, 231–236 (1994).

43. Saponara S, Fusi F, Sgaragli G, Cavalli M, Hopkins B, Bova S. Effects of commonly used protein kinase inhibitors on vascular contraction and L-type Ca(2+) current. Biochemical Pharmacology 84, 1055–1061 (2012).

44. Rorsman NJG, Ta CM, Garnett H, Swietach P, Tammaro P. Defining the ionic mechanisms of optogenetic control of vascular tone by channelrhodopsin-2. British journal of pharmacology 175, 2028–2045 (2018).

45. Asano T, et al. Mechanism of action of a novel antivasospasm drug, HA1077. The Journal of pharmacology and experimental therapeutics 241, 1033–1040 (1987).

46. Roth TL, Nayak D, Atanasijevic T, Koretsky AP, Latour LL, McGavern DB. Transcranial amelioration of inflammation and cell death after brain injury. Nature 505, 223–222 (2014).

47. Schmid F, Reichold J, Weber B, Jenny P. The impact of capillary dilation on the distribution of red blood cells in artificial networks. American Journal of Physiology: Heart and Circulatory Physiology 308, H733–H742 (2015).

48. Wyckoff JB, Pinner SE, Gschmeissner S, Condeelis JS, Sahai E. ROCK- and myosin-dependent matrix deformation enables protease-independent tumor-cell invasion in vivo. Current Biology 16, 1515–1523 (2006).

49. Harraz OF, et al. Ca(V)3.2 channels and the induction of negative feedback in cerebral arteries. Circulation Research 115, 650–661.

50. Alarcon-Martinez L, et al. Capillary pericytes express α-smooth muscle actin, which requires prevention of filamentous-actin depolymerization for detection. Elife 7, e34861 (2018).

51. Hutchinson EB, Stefanovic B, Koretsky AP, Silva AC. Spatial flow-volume dissociation of the cerebral microcirculatory response to mild hypercapnia. Neuroimage 32, 520–530 (2006).

52. Gutiérrez-Jiménez E, Angleys H, Rasmussen PM, Mikkelsen IK, Mouridsen K, Østergaard L. The effects of hypercapnia on cortical capillary transit time heterogeneity (CTH) in anesthetized mice. Journal of Cerebral Blood Flow & Metabolism 38, 290–303 (2018).

53. Watson AN, et al. Mild pericyte deficiency is associated with aberrant brain microvascular flow in aged PDGFRβ+/− mice. Journal of Cerebral Blood Flow & Metabolism Epub ahead of print, (2020).

54. Rungta RL, Osmanski BF, Boido D, Tanter M, Charpak S. Light controls cerebral blood flow in naive animals. Nature Communications 8, 14191 (2017).

55. Berthiaume AA, et al. Dynamic Remodeling of Pericytes In Vivo Maintains Capillary Coverage in the Adult Mouse Brain. Cell Reports 22, 8–16 (2018).

56. Lyons DG, Parpaleix A, Roche M, Charpak S. Mapping oxygen concentration in the awake mouse brain. Elife 5, e12024 (2016).

57. Gutiérrez-Jiménez E, et al. Effect of electrical forepaw stimulation on capillary transit-time heterogeneity (CTH). Journal of Cerebral Blood Flow & Metabolism 36, 2072–2086 (2016).

58. Damisah EC, Hill RA, Tong L, Murray KN, Grutzendler J. A fluoro-Nissl dye identifies pericytes as distinct vascular mural cells during in vivo brain imaging. Nature Neuroscience 20, 1023–1032 (2017).

59. Hudetz AG. Blood flow in the cerebral capillary network: a review emphasizing observations with intravital microscopy. Microcirculation 4, 233–252 (1997).

60. Kleinfeld D, Mitra PP, Helmchen F, Denk W. Fluctuations and stimulus-induced changes in blood flow observed in individual capillaries in layers 2 through 4 of rat neocortex. Proceedings of the National Academy of Sciences USA 95, 15741–15746 (1998).

61. Li Y, Wei W, Wang RK. Capillary flow homogenization during functional activation revealed by optical coherence tomography angiography based capillary velocimetry. Scientific Reports 8, 4107 (2018).

62. Li B, Lee J, Boas DA, Lesage F. Contribution of low- and high-flux capillaries to slow hemodynamic fluctuations in the cerebral cortex of mice. Journal of Cerebral Blood Flow & Metabolism 36, 1351–1356 (2016).

63. Desjardins M, Berti R, Lefebvre J, Dubeau S, Lesage F. Aging-related differences in cerebral capillary blood flow in anesthetized rats. Neurobiology of Aging 35, 1947–1955 (2014).

64. Cipolla MJ, Gokina NI, Osol G. Pressure-induced actin polymerization in vascular smooth muscle as a mechanism underlying myogenic behavior. FASEB Journal 16, 72–76 (2002).

65. Yamin R, Morgan KG. Deciphering actin cytoskeletal function in the contractile vascular smooth muscle cell. Journal of Physiology 590, 4145–4154 (2012).

66. Moreno-Domínguez A, et al. Cytoskeletal reorganization evoked by Rho-associated kinase- and protein kinase C-catalyzed phosphorylation of cofilin and heat shock protein 27, respectively, contributes to myogenic constriction of rat cerebral arteries. Journal of Biological Chemistry 289, 20939–20952 (2014).

67. Gunst SJ, Zhang W. Actin cytoskeletal dynamics in smooth muscle: a new paradigm for the regulation of smooth muscle contraction. American journal of physiology: Cell physiology 295, C576–C587 (2008).

68. Nahirney PC, Reeson P, Brown CE. Ultrastructural analysis of blood-brain barrier breakdown in the peri-infarct zone in young and aged mice. Journal of Cerebral Blood Flow & Metabolism 36, 413–425 (2015).

69. Le Beux YJ, Willemot J. Actin- and myosin-like filaments in rat brain pericytes. The Anatomical Record 190, 811–826 (1978).

70. Tachibana E, et al. Intra-arterial infusion of fasudil hydrochloride for treating vasospasm following subarachnoid haemorrhage. Acta neurochirurgica 141, 13–19 (1999).

71. Shibuya M, Hirai S, Seto M, Satoh S, Ohtomo E. Effects of fasudil in acute ischemic stroke: results of a prospective placebo-controlled double-blind trial. Journal of Neurological Sciences 238, 31–39 (2005).

72. Vesterinen HM, et al. Systematic review and stratified meta-analysis of the efficacy of RhoA and Rho kinase inhibitors in animal models of ischaemic stroke. Systematic reviews 2, 33 (2013).

73. Cauli B, et al. Cortical GABA interneurons in neurovascular coupling: Relays for subcortical vasoactive pathways. Journal of Neuroscience 24, 8940–8949 (2004).

74. Kisler K, Nikolakopoulou AM, Sweeney MD, Lazic D, Zhao Z, Zlokovic BV. Acute Ablation of Cortical Pericytes Leads to Rapid Neurovascular Uncoupling. Frontiers in Cellular Neuroscience 14, 27 (2020).

75. Cruz Hernández JC, et al. Neutrophil adhesion in brain capillaries reduces cortical blood flow and impairs memory function in Alzheimer’s disease mouse models. Nature Neuroscience 22, 413–420 (2019).

76. Jespersen SN, Østergaard L. The roles of cerebral blood flow, capillary transit time heterogeneity, and oxygen tension in brain oxygenation and metabolism. Journal of Cerebral Blood Flow & Metabolism 32, 264–277 (2012).

77. Stefanovic B, et al. Functional reactivity of cerebral capillaries. Journal of Cerebral Blood Flow & Metabolism 28, 961–972 (2007).

78. Hudetz AG. Blood Flow in the cerebral capillary network: A review emphasizing observations with intravital microscopy. Microcirculation 4, 233–252 (1997).

79. Bell RD, et al. Pericytes control key neurovascular functions and neuronal phenotype in the adult brain and during brain aging. Neuron 68, 409–427 (2010).

80. Sengillo JD, Winkler EA, Walker CT, Sullivan JS, Johnson M, Zlokovic BV. Deficiency in mural vascular cells coincides with blood-brain barrier disruption in Alzheimer’s disease. Brain Pathology 23, 303–310 (2013).

81. Schultz N, et al. Amyloid-beta 1-40 is associated with alterations in NG2+ pericyte population ex vivo and in vitro. Aging Cell 17, e12728 (2018).

82. Miners JS, Schulz I, Love S. Differing associations between Aβ accumulation, hypoperfusion, blood-brain barrier dysfunction and loss of PDGFRB pericyte marker in the precuneus and parietal white matter in Alzheimer’s disease. Journal of Cerebral Blood Flow & Metabolism 38, 103–115 (2017).

83. Halliday MR, et al. Accelerated pericyte degeneration and blood-brain barrier breakdown in apolipoprotein E4 carriers with Alzheimer’s disease. Journal of Cerebral Blood Flow & Metabolism 36, 216–227 (2016).

84. Dziewulska D, Lewandowska E. Pericytes as a new target for pathological processes in CADASIL. Neuropathology 32, 515–521 (2012).

85. Chow BW, et al. Caveolae in CNS arterioles mediate neurovascular coupling. Nature 579, 106–110 (2020).

86. Muzumdar MD, Tasic B, Miyamichi K, Li L, Luo L. A global double-fluorescent Cre reporter mouse. Genesis 45, 593–605 (2007).

87. Shih AY, Driscoll JD, Drew PJ, Nishimura N, Schaffer CB, Kleinfeld D. Two-photon microscopy as a tool to study blood flow and neurovascular coupling in the rodent brain. Journal of Cerebral Blood Flow & Metabolism 32, 1277–1309 (2012).

88. Chen M, et al. Simply combining fasudil and lipoic acid in a novel multitargeted chemical entity potentially useful in central nervous system disorders. RSC Advances 5, 37266–37269 (2014).

89. Shin HK, et al. Rho-kinase inhibition acutely augments blood flow in focal cerebral ischemia via endothelial mechanisms. Journal of Cerebral Blood Flow & Metabolism 27, 998–1009 (2007).

90. Bedussi B, et al. Clearance from the mouse brain by convection of interstitial fluid towards the ventricular system. Fluids and barriers of the CNS 12, 23 (2015).

91. Drew PJ, et al. Chronic optical access through a polished and reinforced thinned skull. Nature Methods 7, 981–984 (2010).

92. Shih AY, Mateo C, Drew PJ, Tsai PS, Kleinfeld D. A polished and reinforced thinned skull window for long-term imaging and optical manipulation of the mouse cortex. Journal of Visualized Experiments, http://www.jove.com/video/3742 (2012).

93. Driscoll JD, Shih AY, Drew PJ, Cauwenberghs G, Kleinfeld D. Two-photon imaging of blood flow in cortex. Cold spring harbor protocols 8, 759–767 (2013).

94. Drew PJ, Blinder P, Cauwenberghs G, Shih AY, Kleinfeld D. Rapid determination of particle velocity from space-time images using the Radon transform. Journal of Computational Neuroscience 29, 5–11 (2010).

95. Emmenlauer M, et al. XuvTools: free, fast and reliable stitching of large 3D datasets. Journal of Microscopy 233, 42–60 (2009).

