## Supplementary Figures for "Brain capillary pericytes exert a substantial but slow influence on blood flow"

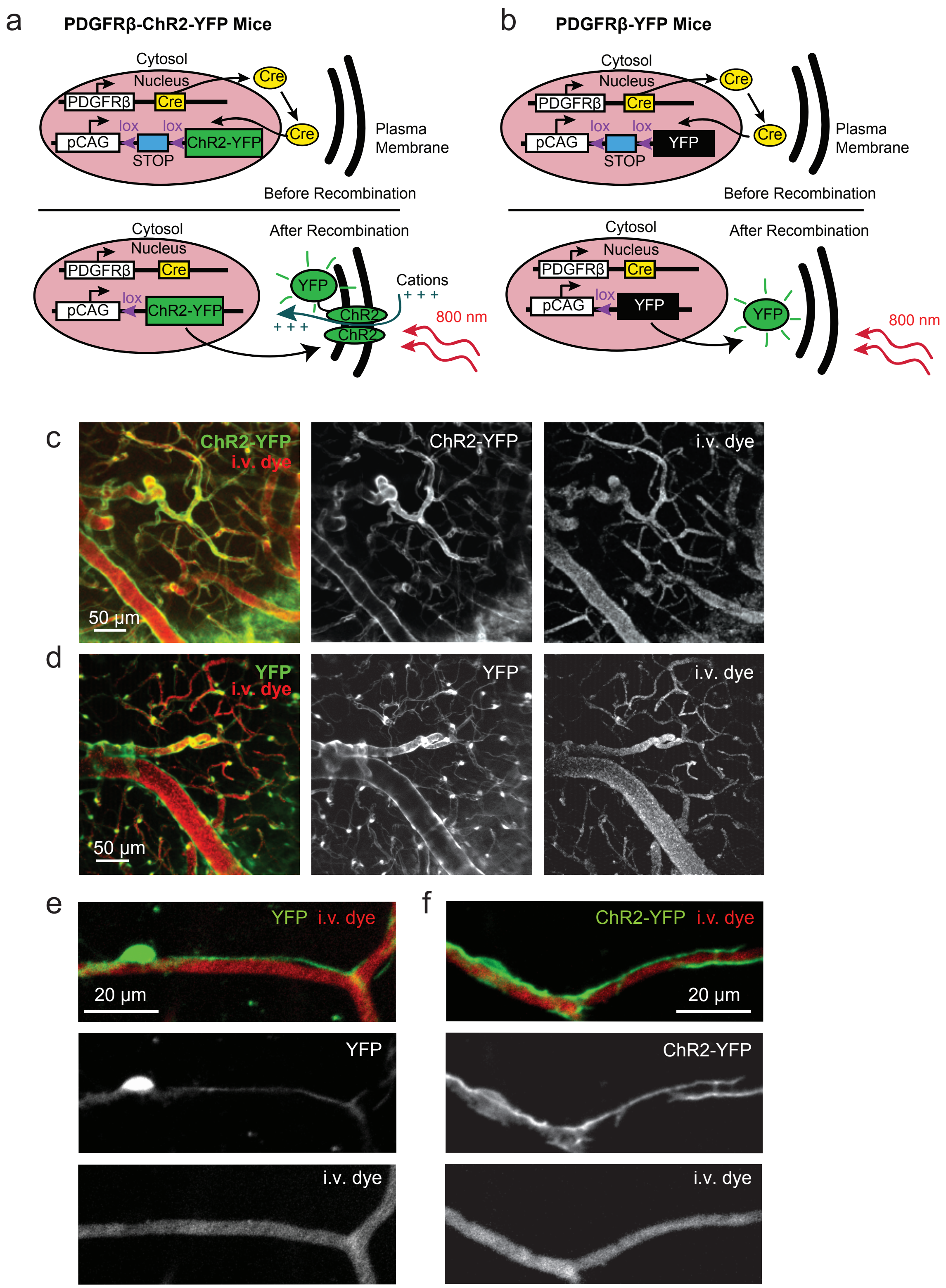


**Supplementary Figure 1. Genetic targeting of YFP and ChR2-YFP to cerebrovascular mural cells.**
**(a,b)** Cross-breeding strategy to create PDGFRβ-ChR2-YFP (a) and PDGFRβ-YFP (b) mice. **(c,d)** *In vivo* two-photon images of cortical vasculature through a cranial window for a PDGFRβ-ChR2-YFP mouse (c) and a PDGFRβ-YFP mouse (d). Maximum projection over 150 µm of cortical thickness. **(e,f)** High-magnification images of capillary pericytes expressing YFP (e), which is a cytosolic protein, versus ChR2-YFP (f), which is membrane-bound. Note how the ChR2-YFP label reveals greater endothelial coverage by capillary pericyte processes, compared to a cytosolic fluorescent protein.

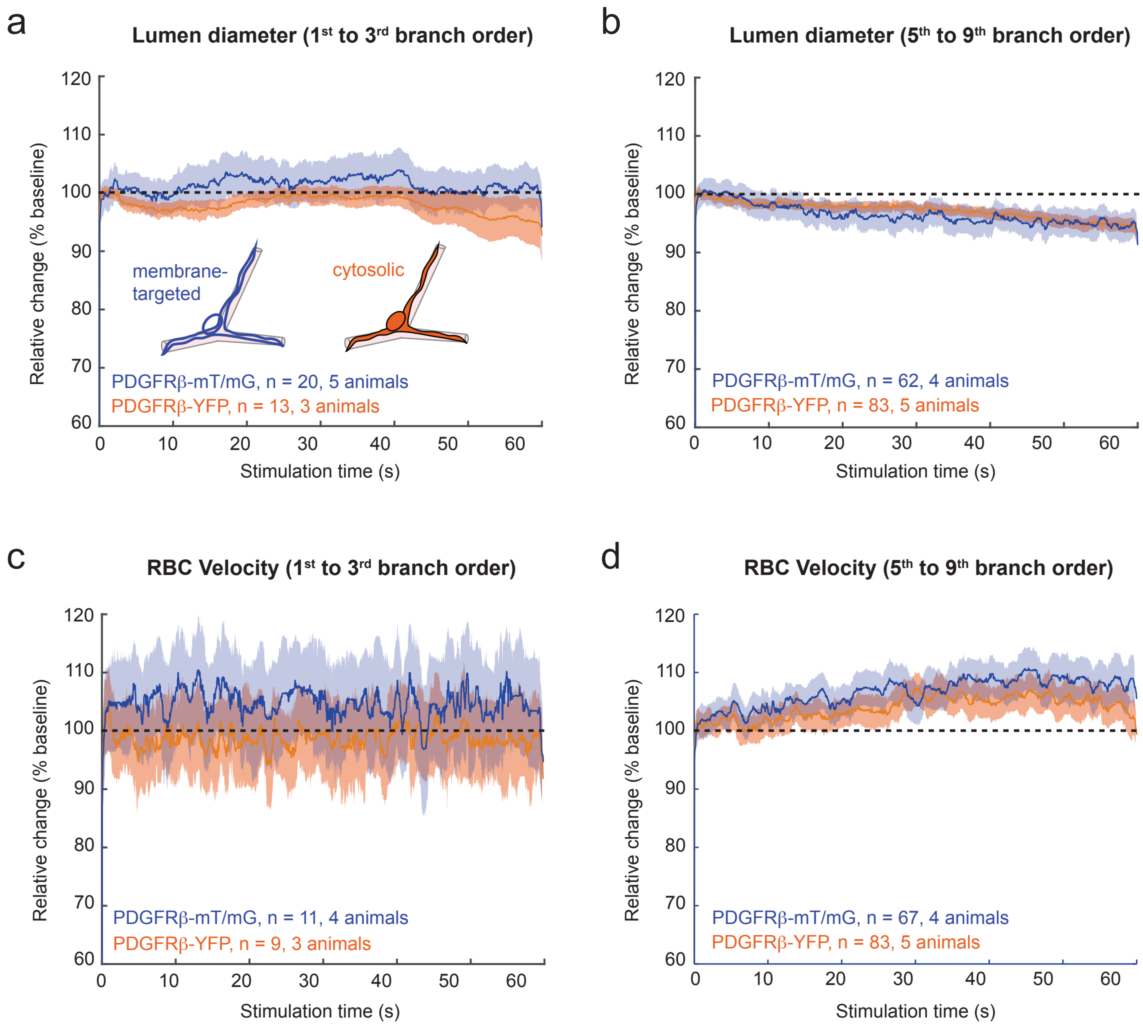


**Supplementary Figure 2. Comparison of non-opsin expressing control mice. (a,b)** Relative change in lumen diameter during two-photon laser line scanning for 1st to 3rd branch order vessels (a), and 5th to 9th branch order capillary vessels (b). Both PDGFRβ-YFP and PDGFRβ-mT/mG mice were examined. Data traces are mean ± SEM. **(c,d)** Relative change in RBC velocity during two-photon laser line scanning for the same microvascular zones

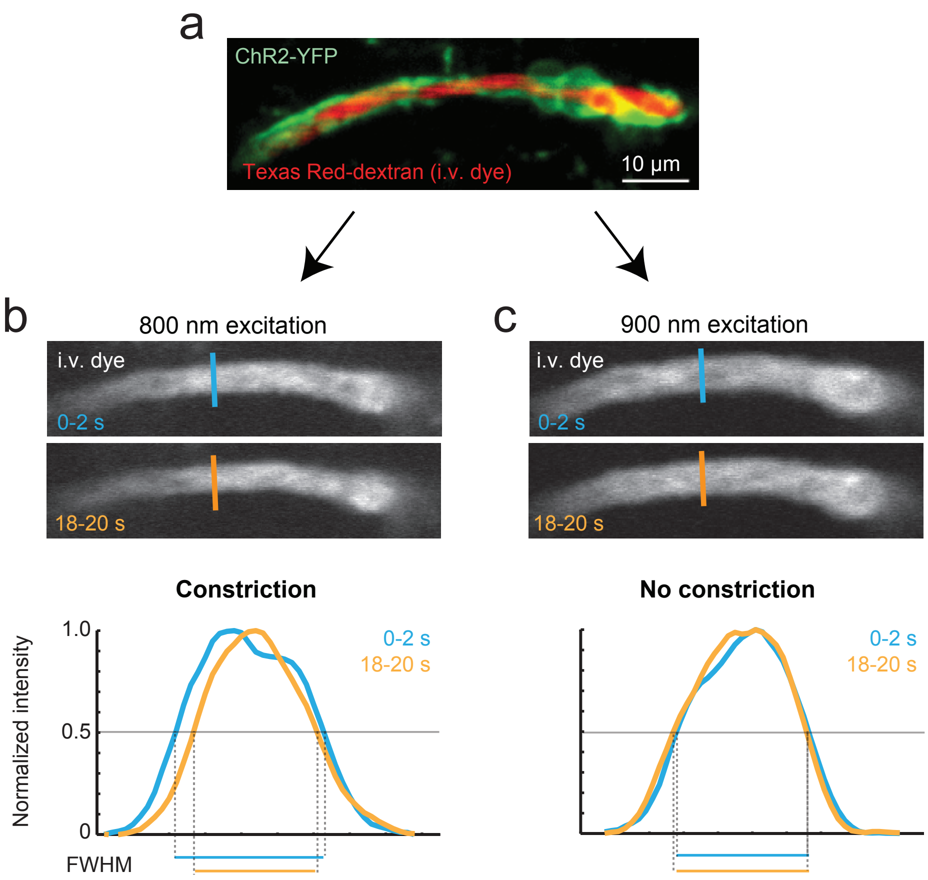


**Supplementary Figure 3. Excitation wavelength dependence of ChR2-YFP activation.**
**(a)** A capillary pericyte *in vivo* (green) covering a capillary with intraluminal space labeled by Texas Red-dextran. **(b)** Planar imaging of this vessel using 800 nm excitation leads to vasoconstriction of the lumen over a period of 20 seconds. A full-width at half max measurement of the intensity profile taken across the vessel width shows a narrower lumen at 18-20 seconds (orange) compared to 0-2 seconds (blue). **(c)** Imaging at 900 nm leads to no obvious vasoconstriction over the same time frame, and is thus used to navigate the sample without significant ChR2 activation

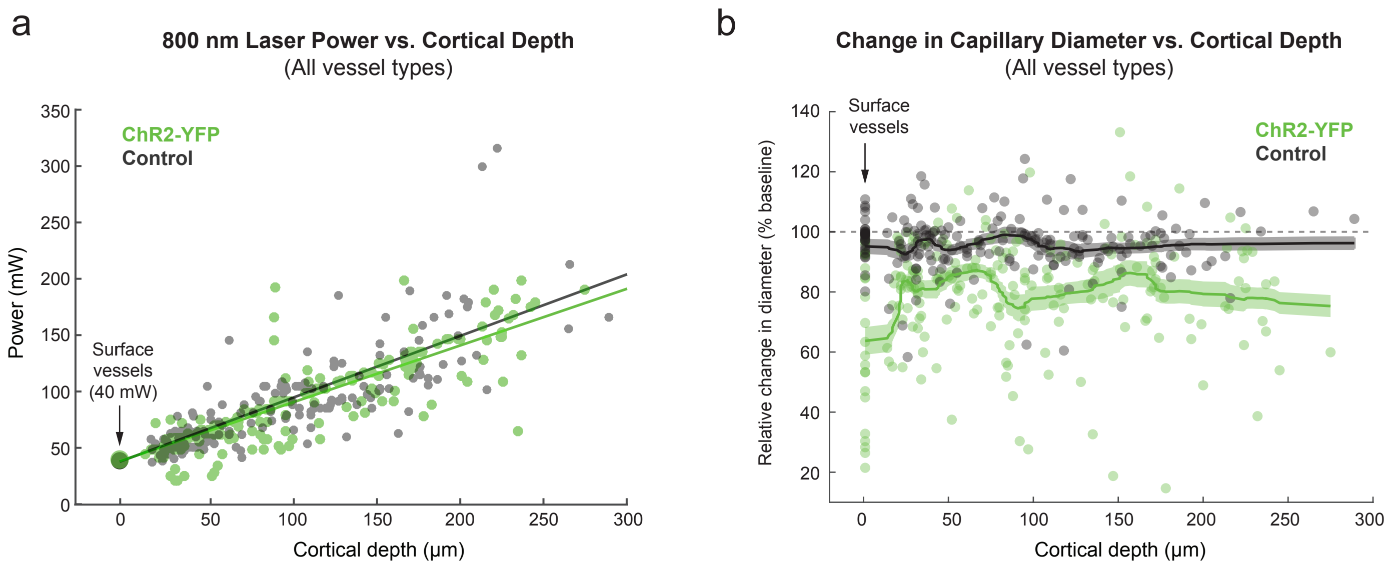


**Supplementary Figure 4. Comparison of optogenetic laser power and change in microvessel diameter as a function of cortical depth.** **(a)** Laser power (800 nm) at the microscope objective plotted as a function of depth below pial surface for the stimulated vessel. Power levels are matched well between ChR2-YFP and control groups. **(b)** Relative change in lumen diameter for target vessel as a function of cortical depth. Laser powers used at different cortical depths produces relatively similar average constrictive response of vessels. Mean ± SEM (moving average window of 25 points).

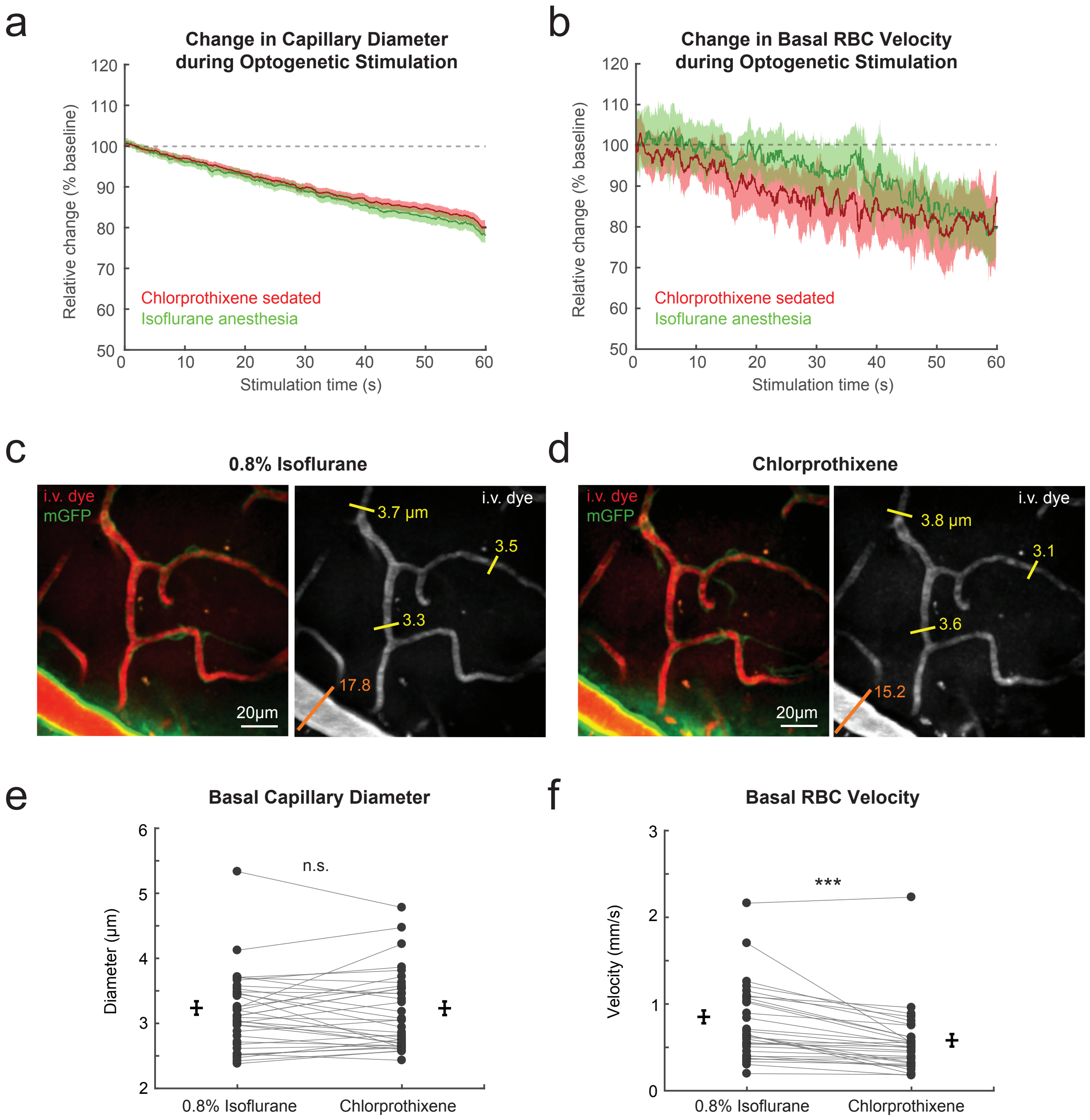


**Supplementary Figure 5. Effect of anesthetic on basal and optogenetically-modulated capillary vasodynamics.** **(a)** Change in diameter of capillaries (5th-9th branch order) during optogenetic activation. Time-course traces between isoflurane anesthetized (0.8% MAC in air) and chlorprothixene sedated PDGFRβ-CHR2-YFP mice are overlaid; n = 63 capillaries from 4 mice in the chlorprothixene group, n = 155 capillaries from 10 mice in the isoflurane group. **(b)** Change in RBC velocity of capillaries during optogenetic activation; n = 65 capillaries from 4 mice in the chlorprothixene group, n = 160 capillaries from 10 mice in the isoflurane group. **(c,d)** In separate PDGFRβ-mT/mG mice with a chronic cranial windows, capillary diameter and RBC velocity was measured under 0.8% isoflurane. Measurements were then repeated in the same capillaries between 15-90 minutes after injecting chlorprothixene (30 μL of 1 mg/mL i.m.) and cessation of isoflurane delivery. Maximum projection overlays near the cortical surface show capillaries (yellow lines, numbers are diameter in micrometers) and pial arterioles (orange line) with each anesthetic. **(e,f)** Capillaries did not differ in diameter between isoflurane anesthesia and chlorprothixene sedation; $p > 0.1$ by paired t-test, n = 35 capillaries from 3 mice. However, arterioles are more dilated during isoflurane. As a result, RBC velocity significantly decreased after isoflurane washout; ***$p<0.001$ by paired t-test, n= 31 capillaries from 3 mice. Data presented as mean ± SEM.

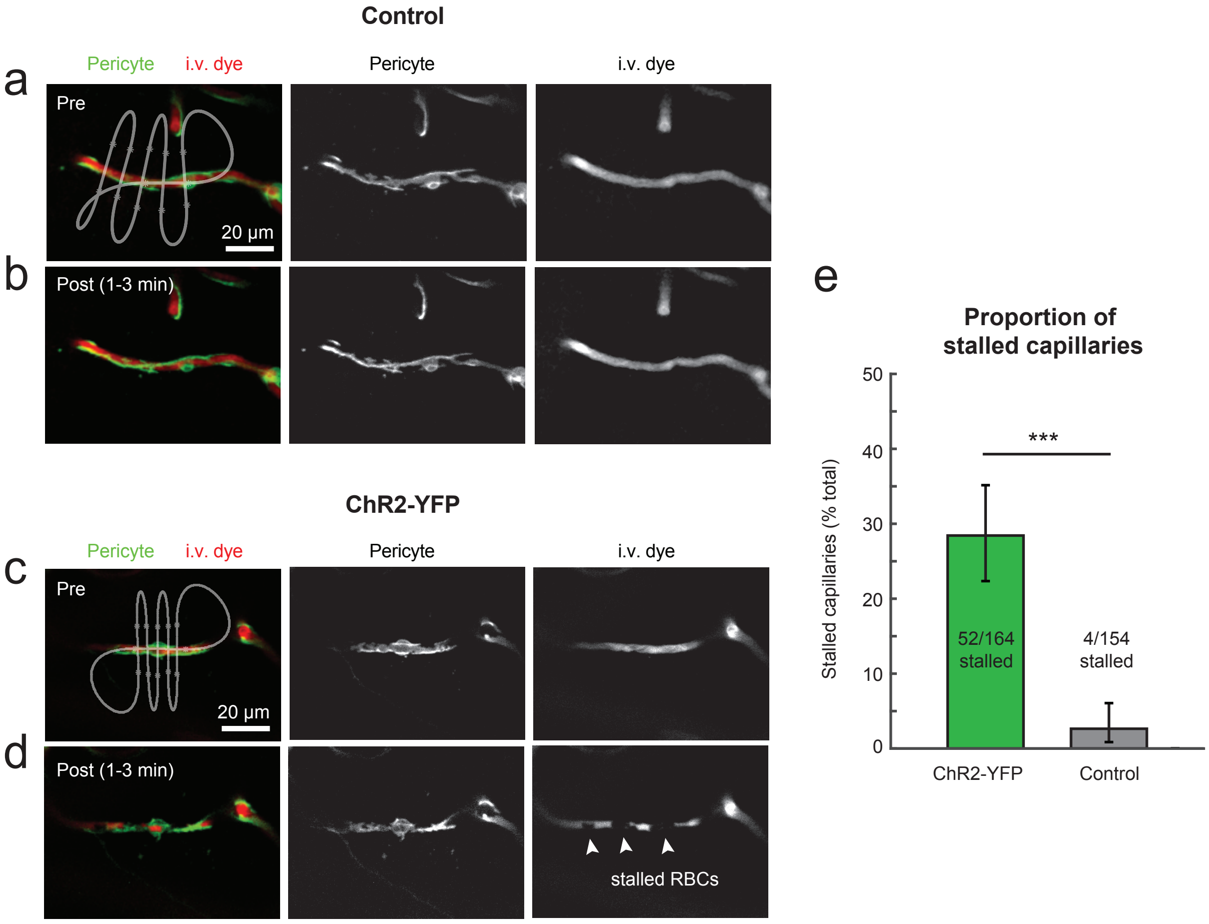


**Supplementary Figure 6. Capillary constriction and stalling induced by optogenetic activation of capillary pericytes.** **(a,b)** A 3-D image stack was collected at 900 nm before and after stimulating ChR2, which was done with a line-scan at 800 nm. Example images from a control mouse with no constriction of stalling RBCs. **(c,d)** Example of capillary constriction and stalled RBCs in the post-scan image from a ChR2-YFP mouse. **(e)** Proportion of capillaries (5th-9th branch order) stalling in ChR2-YFP and control groups. Presented as # capillaries stalled / # total capillaries analyzed. Data is mean +/- 95% confidence interval, *p <0.0001, generalized linear mixed model incorporating hierarchical data. N = 164 total vessels (10 animals) for ChR2-YFP, N= 154 total vessels (9 animals) for Control.

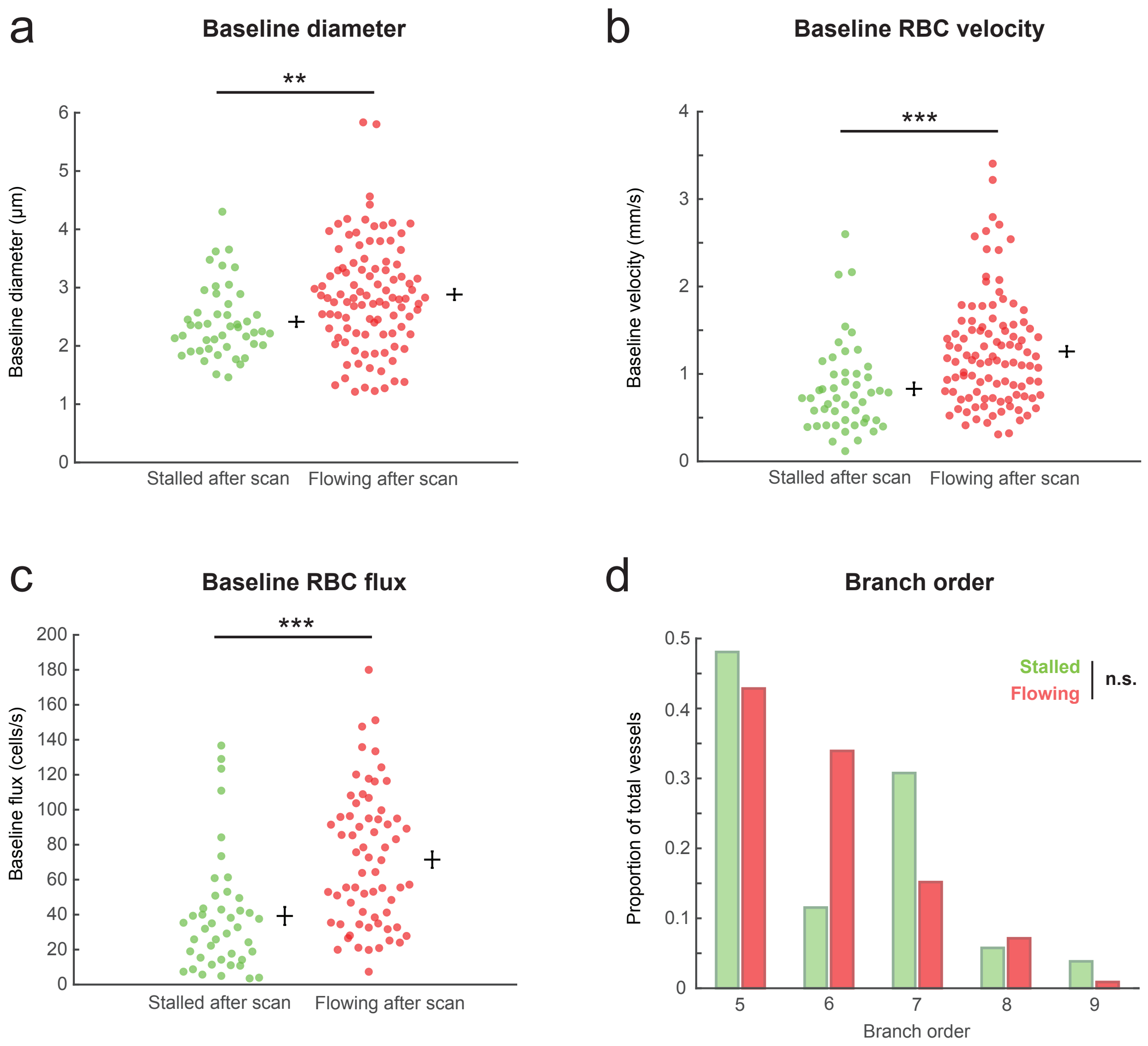


**Supplementary Figure 7: Baseline capillary characteristics predict stalling after ChR2-YFP stimulation.** In image stacks collected after 60 seconds of ChR2-YFP excitation, the flow in a subset of capillaries were stalled (green), while most were flowing (red). Vessels were not stalled when stimulating non-opsin control mice, as shown in previous figures. (a) Baseline diameter was significantly smaller for the vessels that were found stalled after ChR2 stimulation (**$p<0.005$ by Wilcoxon Rank-sum test, n=47 stalled, 103 flowing). (b) Baseline RBC velocity was significantly lower in the vessels that were found stalled after ChR2 stimulation (***$p<0.001$ by Wilcoxon Rank-sum test, n=49 stalled, 104 flowing). (c) Baseline RBC flux was significantly less for the vessels that were found stalled after ChR2 stimulation (***$p<0.001$ by Wilcoxon Rank-sum test, n=43 stalled, 66 flowing). (d) Branch order distribution was not different among vessels that were flowing or stalled after the scan ($p>0.2$ by Kolmogorov-Smirnov test). Altogether, this indicates that vessels with smaller diameter, slower flow, and lower flux, are more likely to stall during pericyte contraction.

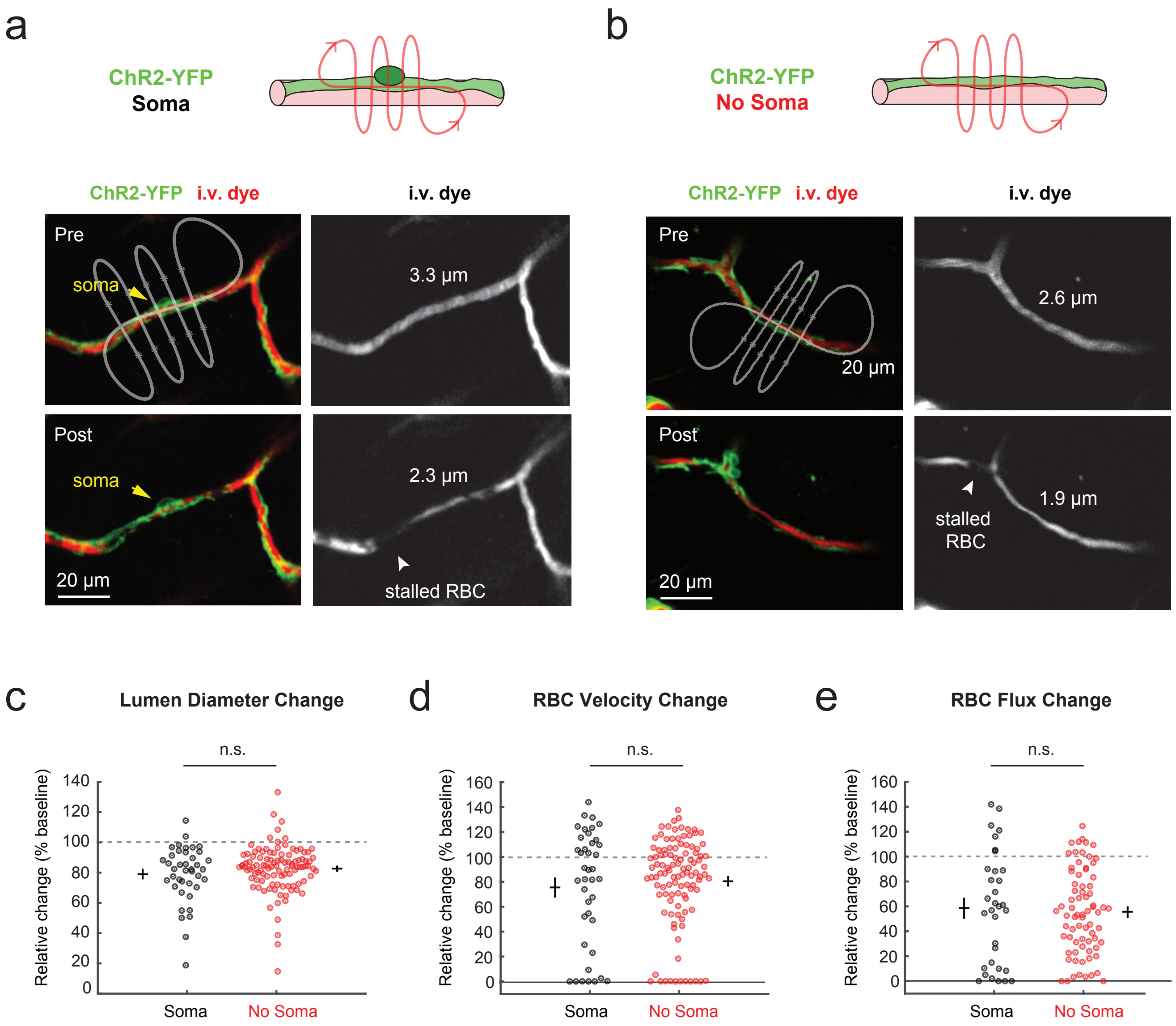


**Supplementary Figure 8: Capillary constriction occurs whether optogenetic stimulation is applied to the pericyte somata or its processes.** **(a)** A line-scan applied to the pericyte soma in a ChR2-YFP mouse leads to sustained capillary constriction and RBC stalling. **(b)** Stimulation of capillary regions devoid of pericyte somata also leads to sustained capillary constriction and RBC stalling. Image stacks in the "No soma" group were carefully examined to ensure there were no pericyte somata above or beneath the lumen, hidden away from view. **(c-e)** Change in lumen diameter, RBC velocity and RBC flux induced by optogenetic pericyte stimulation in ChR2-YFP mice does not differ whether the scan is applied at the pericyte soma (black) or away from the soma (red). N = 40, 40, 33 vessels with soma in scan path, and n = 105, 107, 74 vessels without soma in scan path for panels c,d,e, respectively. N = 10 animals for all. Comparisons made using a Wilcoxon rank-sum test, all p>0.1.

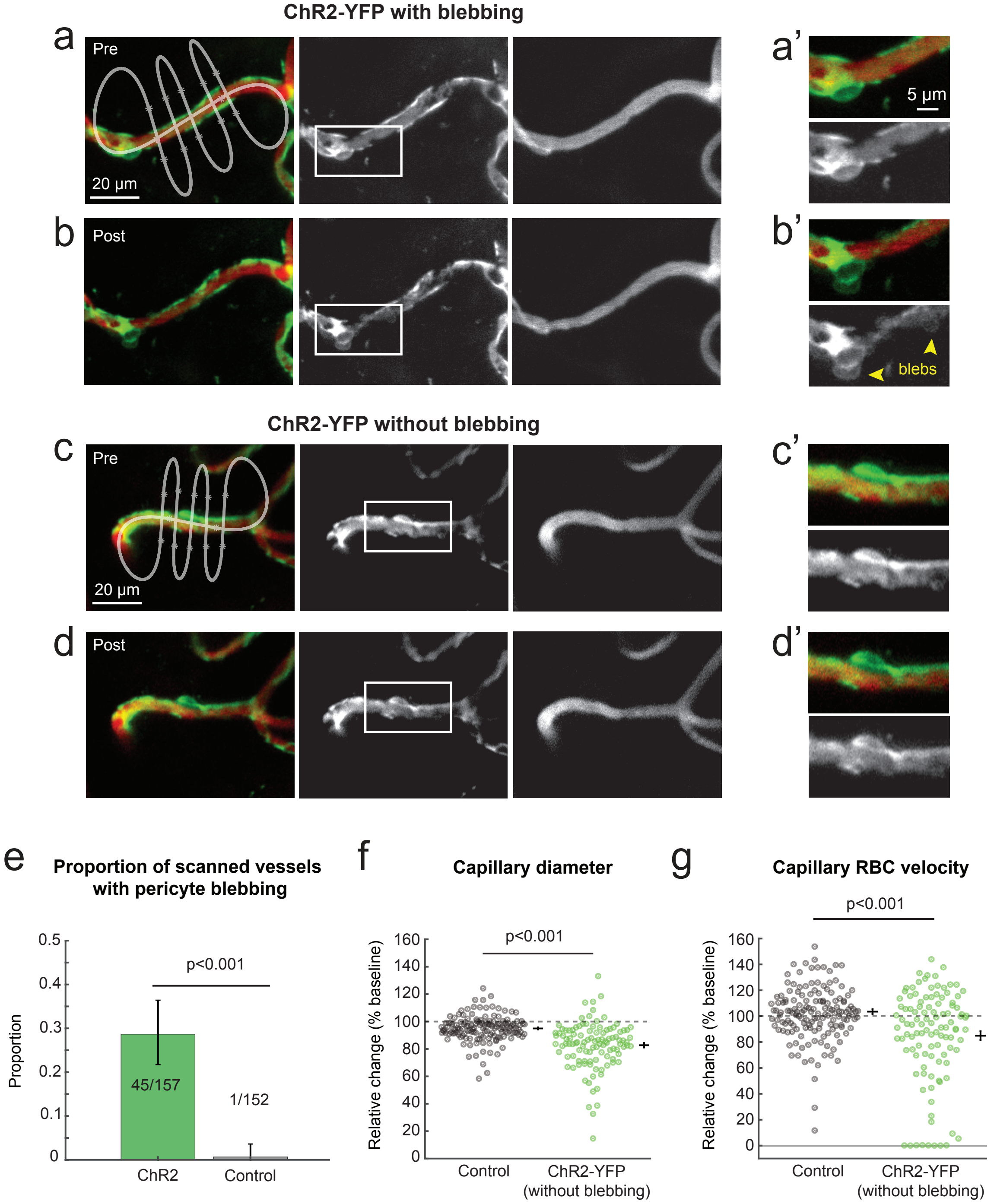


**Supplementary Fig. 9: Capillary pericyte blebbing in subset of vessels stimulated in ChR2-YFP mice.** **(a,a')** Example of a 6th order ChR2-YFP capillary pericyte that blebbed after scanning along the path shown in white. The a' image is an inset of the pericyte soma. **(b,b')** After 60 second of scanning, this pericyte showed blebs on the soma and processes (arrowheads). A 20% capillary constriction from baseline was recorded for this stimulation. **(c,c',d,d')** Example of a 5th order ChR2-YFP capillary pericyte that did not bleb, but also exhibited a 20% capillary constriction. **(e)** Proportion of pericytes with blebbing in each ChR2-YFP and control groups. Chi-Square test. **(f,g)** Removing the ChR2-YFP data with blebbing did not affect our conclusions on diameter (f) or RBC velocity (g). N=107 vessels (10 animals) for ChR2-YFP, and n=145 (9 animals) control mice in panel (f); N=108 vessels (10 animals) ChR2-YFP, and n=152 (9 animals) control mice in panel (g). Statistics were performed with Mann-Whitney U tests.

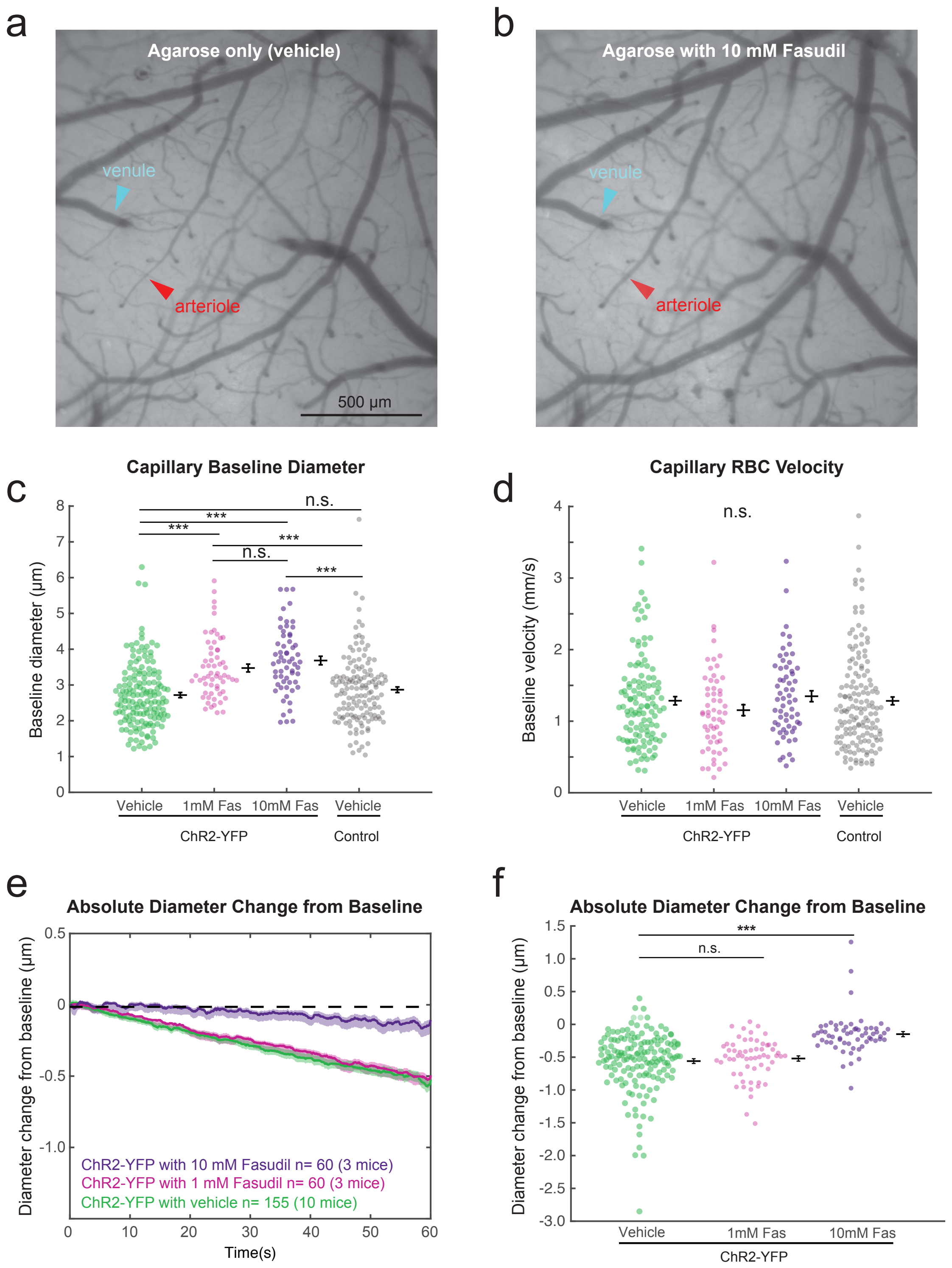


**Supplementary Figure 10. Fasudil dilates microvessels.** **(a,b)** Pial vessels through a cranial window before (a) and after (b) application of agarose containing 10 mM fasudil. Arterioles (red arrow) are dilated, but venules (blue arrow) are unchanged. **(c)** Baseline diameter of capillaries ($5^{th}$-$9^{th}$ branch order) as measured from in vivo two photon line-scans. Mean ± SEM. ***$F(3,416)=22.7$ (overall $p<0.001$) with Tukey-adjusted $p < 0.001$, by one-way ANOVA (n = 155 (10 mice), 60 (3 mice), 60 (3 mice), and 145 (9 mice) capillaries for ChR2-YFP vehicle, ChR2-YFP 1mM fasudil, ChR2-YFP 10mM fasudil, and control mice with vehicle, respectively. **(d)** Baseline capillary RBC velocity, mean ± SEM. n = 117 (10 mice), 57 (3 mice), 61 (3 mice), and 150 (9 mice) capillaries for ChR2-YFP vehicle, ChR2-YFP 1mM fasudil, ChR2-YFP 10mM fasudil, and control mice with vehicle, respectively. Not significant by one-way ANOVA. $F(3,381)=0.95$, overall $p>0.1$. **(e)** Absolute diameter change from baseline. N values shown on graph. **(f)** Absolute diameter change at 60 seconds shows a significant difference between vehicle and 10mM fasudil. Kruskal-Wallis test, df 272, Chi-square 59, ***$p<0.0001$. Same n values as (e)

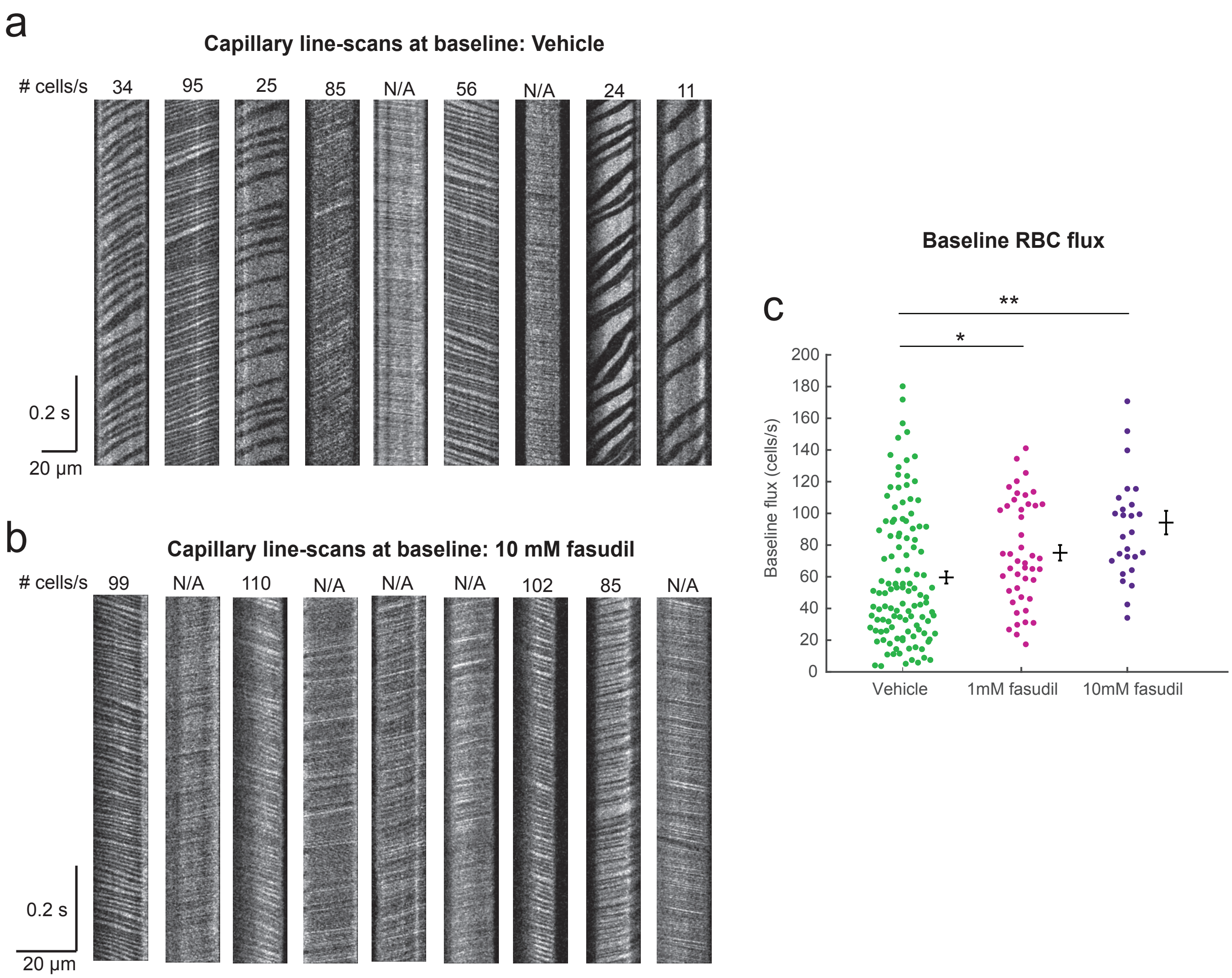


**Supplementary Figure S11. Fasudil increases baseline blood cell flux. (a,b)** Example capillary line-scans from vehicle (panel a) and 10mM fasudil (panel b) treated groups that were selected by a random number generator. Baseline RBC flux values are shown above each image. Note that many could not be counted (N/A) from the Fasudil group because the flux was too high and individual RBCs could not be distinguished. Images represent constant velocity portion of a line-scan from which RBC flux and velocity metrics are extracted. **(c)** The manually-counted baseline RBC flux is significantly greater than vehicle for both the 1 mM fasudil and 10 mM fasudil groups. Kruskal-Wallis test: Chi-square 21.38, df 2, overall p<0.0001. Adjusted *p=0.015, **p<0.001 by Tukey post-hoc test, n = 116, 45, 27 for vehicle, 1 mM fasudil and 10 mM fasudil, respectively). All groups are ChR2-YFP vessels at branch orders 5-9. Note that scatter plots contain only data points with RBC flux that could be counted, which excludes many high flux vessels.

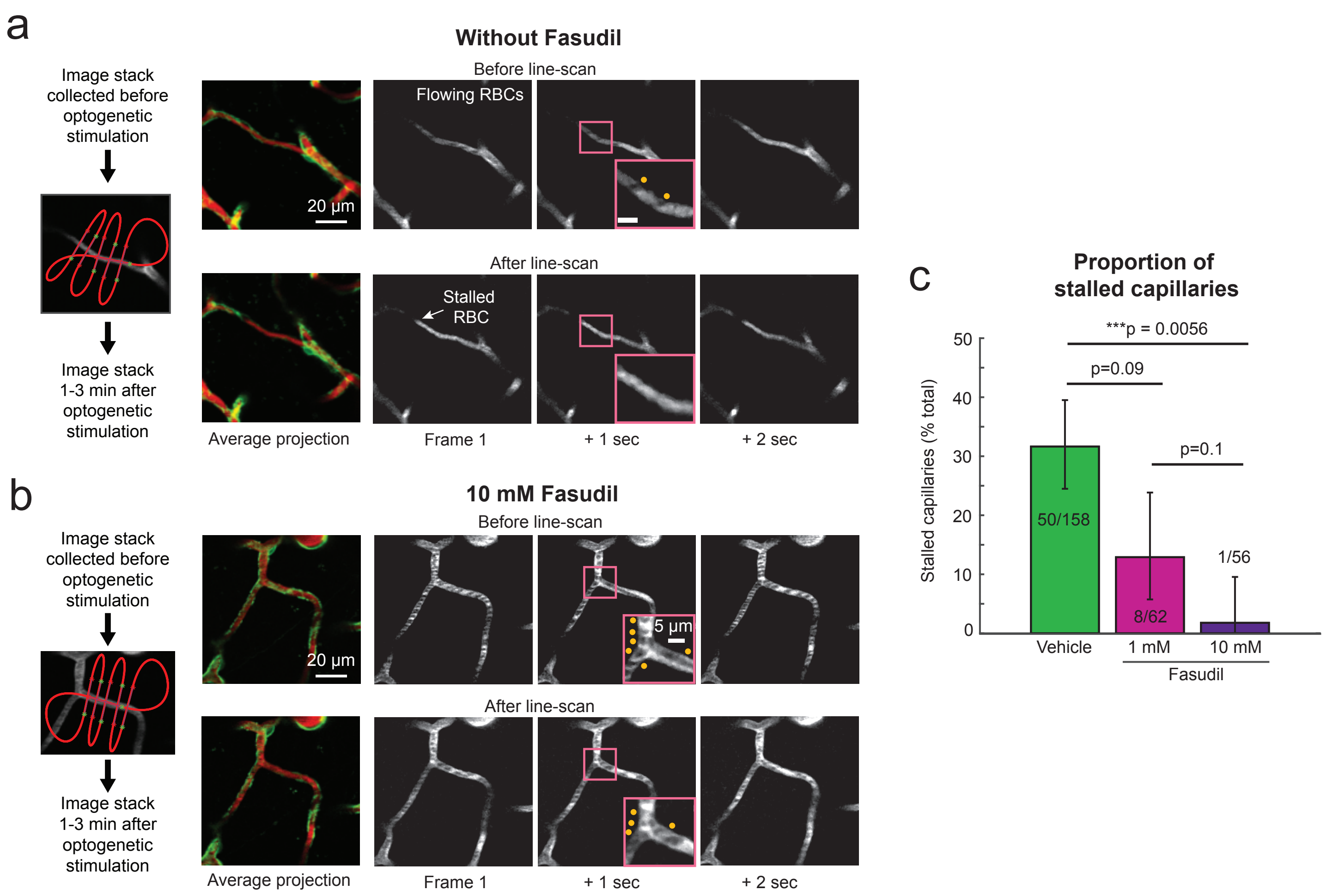


**Supplementary Figure 12: Fasudil prevents flow stalls after capillary pericyte stimulation.** **(a)** Single slices (1 µm apart) through a 30 µm image stack collected before line-scan stimulation of a ChR2-YFP pericyte shows blood cells flowing through the target vessel (inset in pink shows moving RBC shadows, denoted with yellow dots. Inset scale 5µm). Images from the same region after line-scan stimulation show a stalled RBC and no flow through the scanned region. **(b)** This image stack collected before line-scan stimulation of a ChR2-YFP pericyte shows flowing blood cells. The mouse recieved 10 mM fasudil in agarose above the brain surface. The same region after line-scan stimulation shows freely-flowing blood cells. **(c)** Fasudil dose-dependently prevented the stalling of blood cell flow after stimulation (mean +/- 95% binomial fit CI). Repeated measures ANOVA, $F(2,267)=4.89$, overall $p=0.0082$. N values (# stalled/ # total vessels) shown, as are p values adjusted after Tukey-Kramer test.

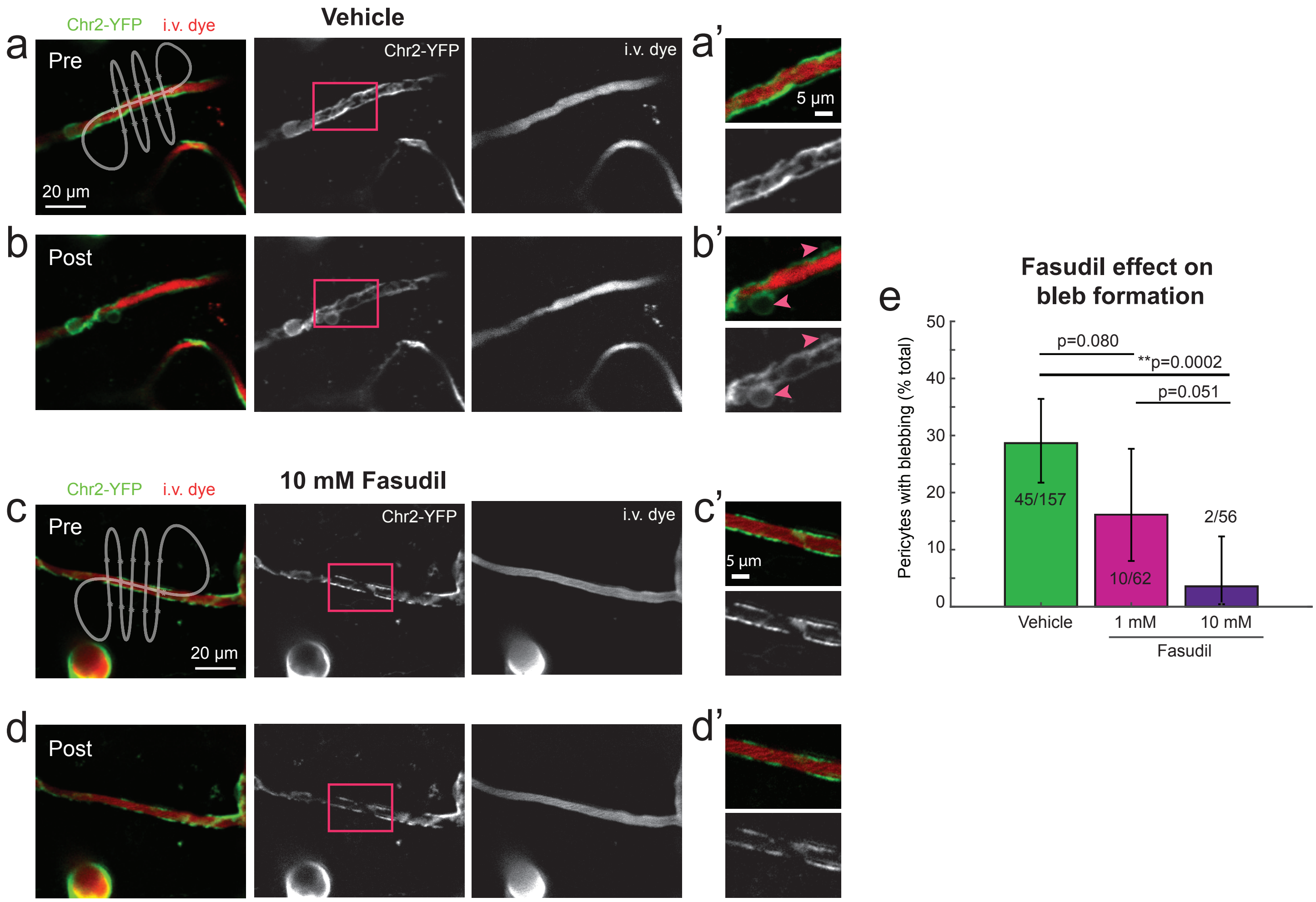


**Supplementary Figure 13: Pericyte blebbing is dose-dependently inhibited by Fasudil. (a,a')** An example of a 7th order ChR2-YFP capillary pericyte never exposed to fasudil scanned along the path shown in white. Panel a' is an inset of the processes of the pericyte. **(b,b')** After 60 seconds of scanning, this pericyte showed blebs on the processes (pink arrowheads). A 24% constriction of lumen diameter was also measured. **(c,c',d,d')** An example of a 6th order ChR2-YFP capillary pericyte exposed to 10 mM fasudil with the absence of blebbing. A 4% constriction was measured. **(e)** Bar graph showing the effect of fasudil on incidence of pericyte blebbing after optogenetic stimulation, mean +/- 95% CI. Blebbing in vehicle-treated capillaries is significantly different than 10 mM fasudil-treated capillaries by Chi-Squared test. Data collected from 10, 3, and 3 animals for vehicle, 1mM and 10mM, repsectively. N values (# with blebs/ # total vessels) are shown.

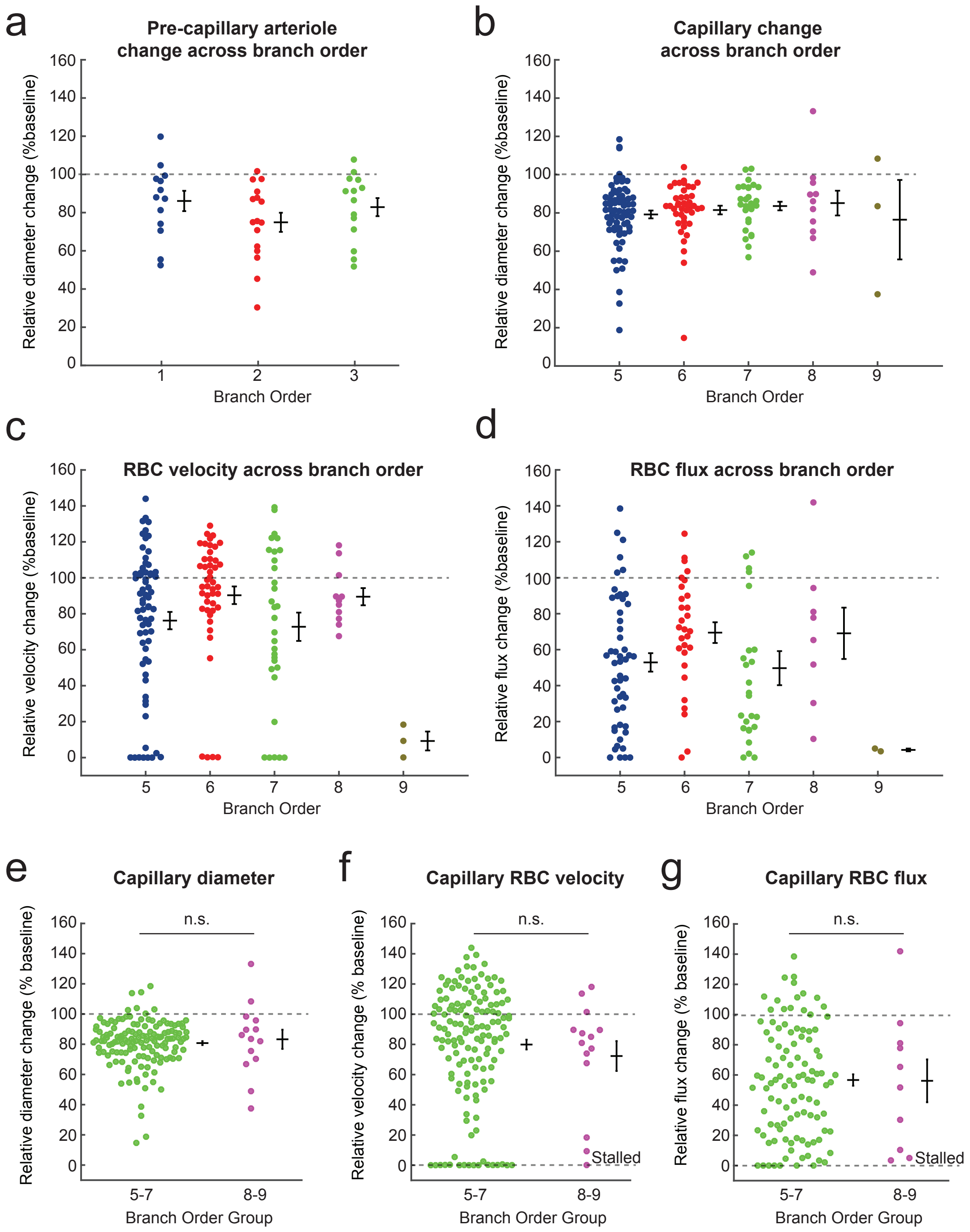


**Supplementary Figure S14. The relative vasodynamic response to ChR2 stimulation is similar across branch orders 1-9. (a)** Each branch order with greater likelihood of ensheathing pericytes (branch orders 1-3) have a similar relative change in diameter at 60 seconds of stimulation; n = 13, 16, 14 vessels from 10 mice for branch orders 1, 2, 3 respectively. **(b-d)** Each branch order harboring capillary pericytes (branch order 5-9) has a similar relative change in *(b)* diameter, *(c)* RBC velocity, and *(d)* RBC flux at 60 seconds of stimulation. For branch orders 5, 6, 7, 8, 9, respectively: *(b)* n = 70, 42, 28, 11, 3 vessels; *(c)* n = 67, 45, 31, 11, 3; *(d)* n = 51, 29, 26, 8, 2, collected over a total of 10 mice. **(e-g)** Capillary branch orders 5-7 and 8-9 exhibit similar changes in *(e)* diameter, *(f)* RBC velocity, and *(g)* RBC flux at 60 seconds of ChR2 stimulation. No difference was detected between groups for these hemodynamic parameters; n.s.means p>0.1 by Wilcoxon rank-sum test. All scatter plots also show Mean +/- SEM.

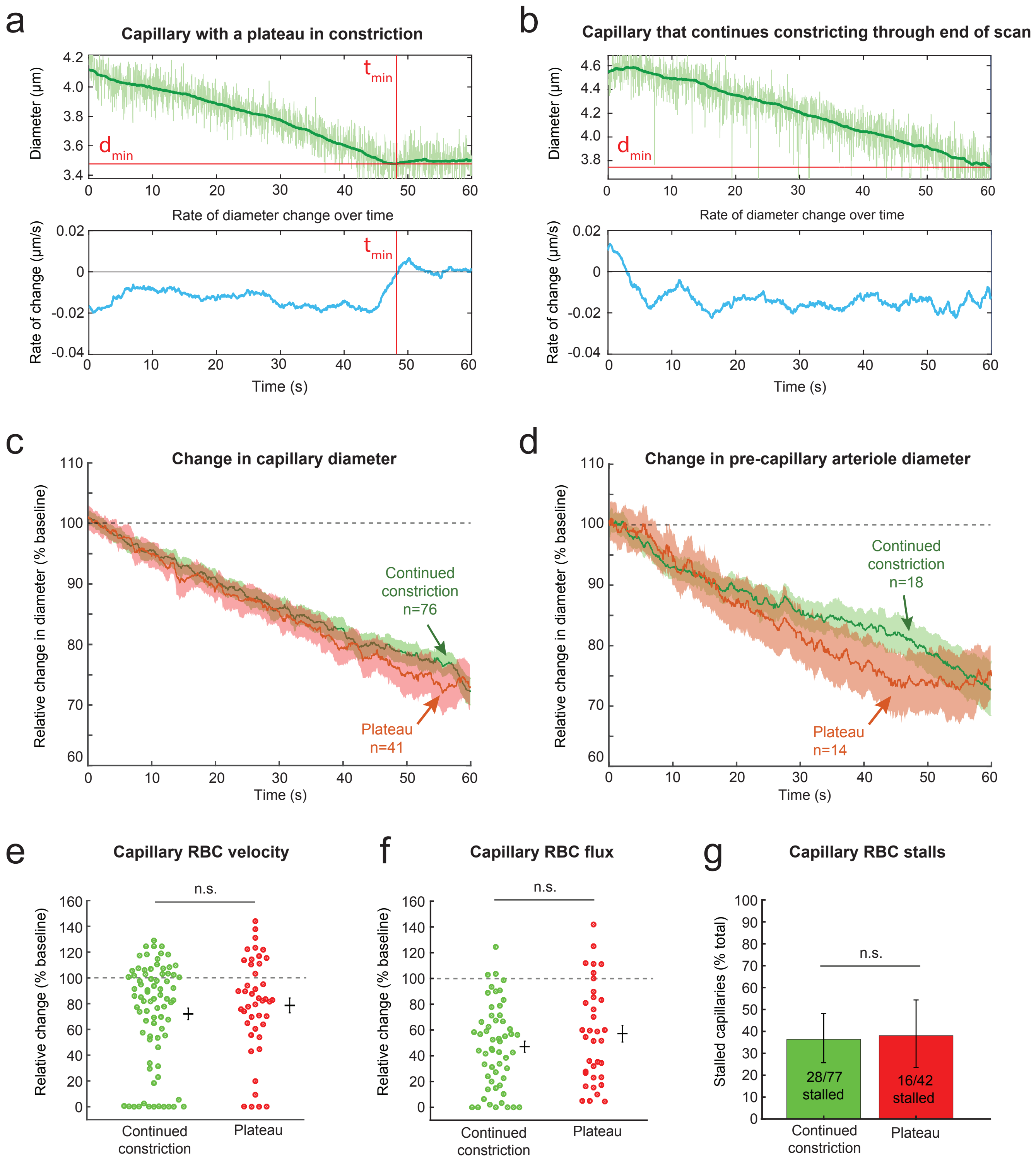


**Supplementary Figure 15: A subset of capillaries in ChR2-YFP mice reach maximal constriction by 60 seconds, but their vasodynamic parameters do not differ from capillaries that continue constricting. (a)** Example of a capillary that constricts during the scan, but reaches a minimum diameter ($d_{min}$) at ~ 49 s ($t_{min}$). Light green shows raw diameter and dark green line shows a median filter of the diameter. Bottom panel: The rate of diameter change crosses from negative to positive to confirm that a minimum diameter had been reached. This is a 5 second moving average of the calculated rate. **(b)** An example of a capillary that did not reach a maximal constriction during the period of stimulation. **(c)** The Mean ± SEM of ChR2-YFP capillaries that reached a plateau (red, n = 41), compared to capillaries that continued to constrict (green, n = 76). Note a slight bend in the curve of the red trace, whereas the green trace continues to decrease (arrows). **(d)** Pre-capillary arterioles (branch orders 1-3) showed similar plateau (n = 14) or continued constriction (n = 18). **(e,f,g)** The capillaries (branch orders 5-9) that did or did not plateau were similar in terms of vasodynamic consequences of constriction; *(panel e)* n=79/44, Rank-sum, p>0.1; *(panel f)* n=55/35, Rank-sum, p>0.1; *(panel g)* n=77/42 total capillaries, Chi-square, p>0.9. mean ± 95% CI shown in *panel g*. Mean ± SEM elsewhere. Vessels with less than a 10% constriction were excluded.

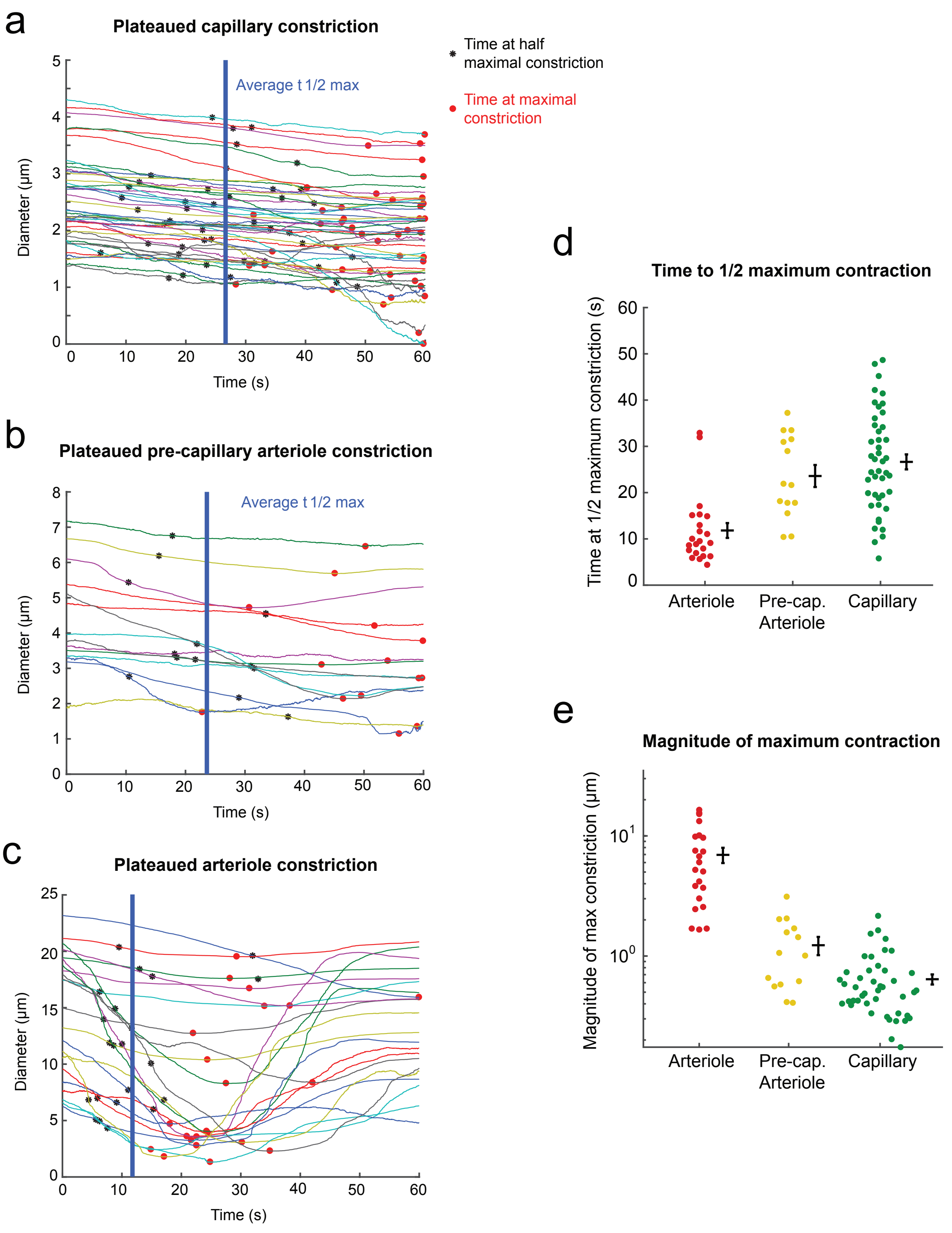


**Supplementary Figure 16: Comparison of contraction kinetics among vessel types in ChR2-YFP mice (line-scan data).** **(a)** A subset of stimulated ChR2 capillaries (branch orders 5-9) had lumen diameters that reached a plateau prior to the end of the scan. Each line is an individual capillary diameter trace that has been median filtered over 10 seconds. The asterisk indicates the time at 1/2 maximum constriction for an individual vessel. The red dot indicates the time at which the minimum diameter (maximum constriction) occurred. The vertical blue line is the average time of 1/2 maximum constriction. **(b,c)** Same as (a) for pre-capillary arterioles (b), and arterioles (c). **(d)** Speed to half maximum constriction follows a trend of arterioles > precapillary arterioles > capillaries. **(e)** The difference between baseline and minimum diameters, representing the magnitude of maximum constriction, follows a trend of arterioles > precapillary arterioles > capillaries. N=22 arterioles, 14 pre-capillary arterioles, and 44 capillaries across 10 mice.

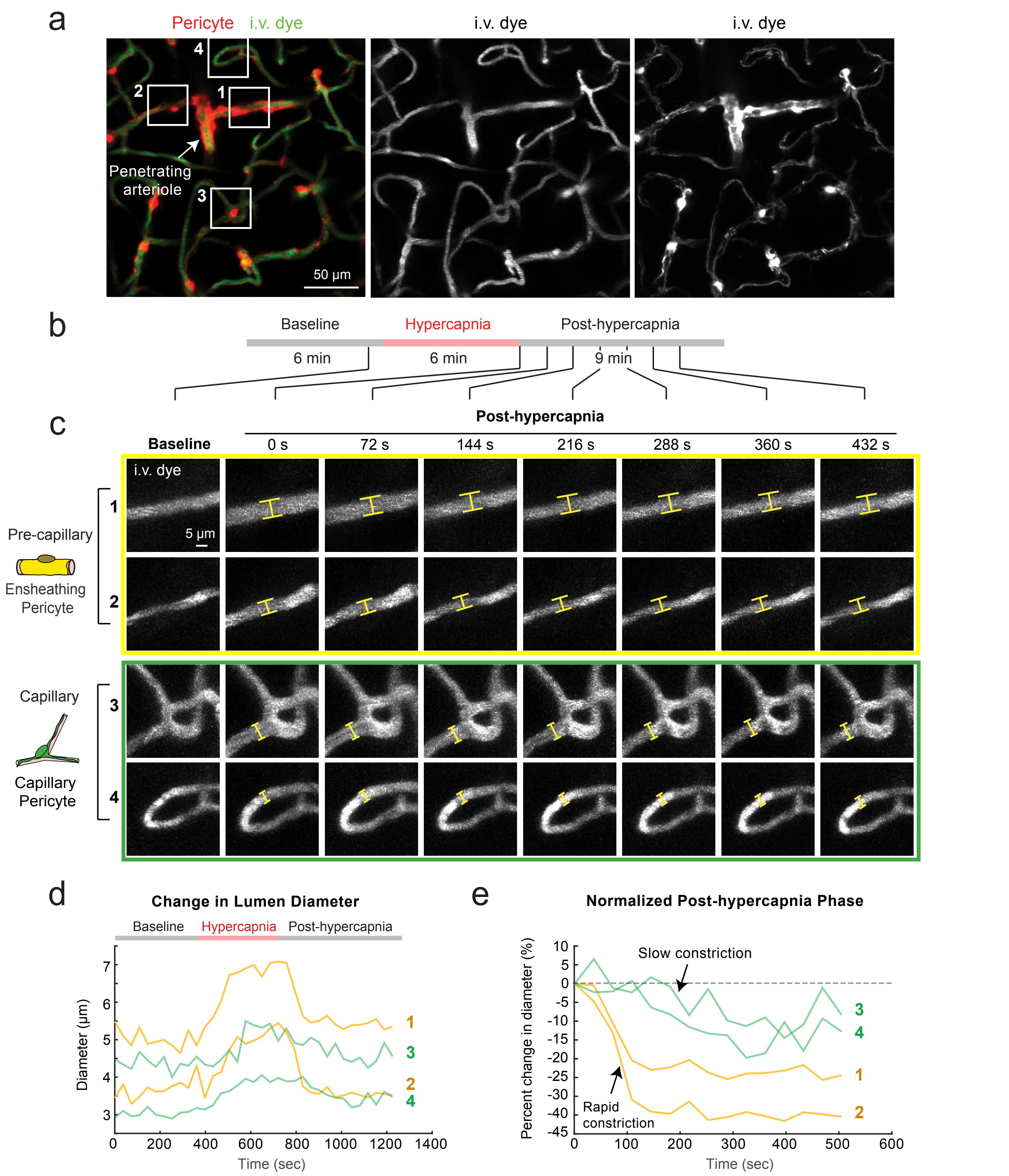


**Supplementary Figure 17: Microvessel contriction following hypercapnia-induced vasodilation.** **(a)** Maximal projection of a 50 μm thick image stack from PDGFRβ-tdTomato mouse cortex, through a thinned-skull window. Image stack were repeatedly collected every 36 seconds before, during and after hypercapnic challenge. **(b)** Time-course and phases of the hypercapnia experiment. **(c)** Magnified views of two 1st branch order pre-capillary arterioles and two capillary regions, covered by ensheathing and capillary pericytes, respectively. Capillary 3 is 7th branch order and capillary 4 is 5th branch order. Insets in panel a depict the regions selected for magnified views. **(d)** Absolute lumen diameter plotted as a function of time for each microvessel. **(e)** Normalized diameter change for each microvessel during the post-hypercapnia phase.

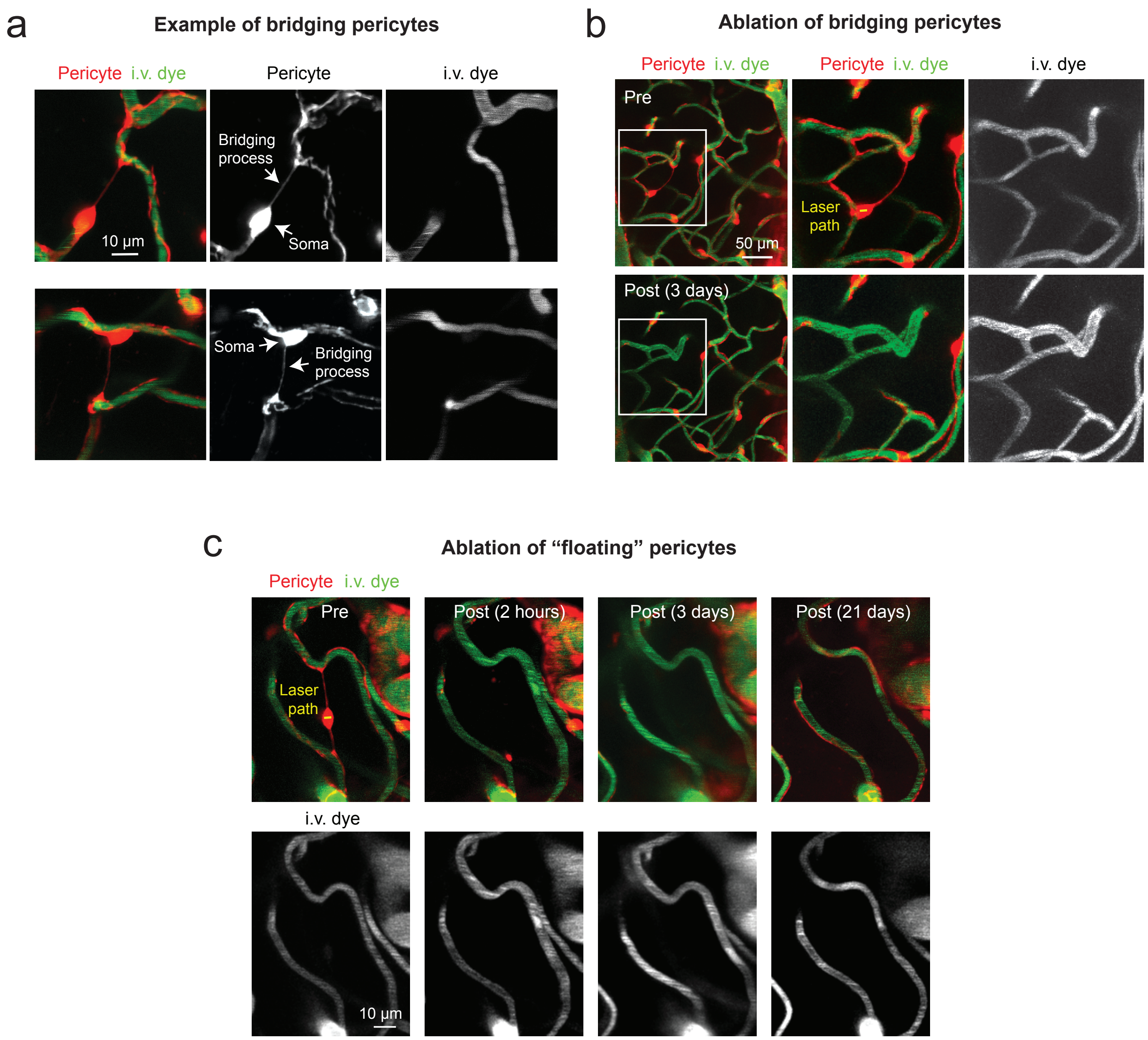


**Supplementary Figure 18: Additional examples of bridging pericyte ablations.** **(a)** Two examples of bridging pericytes. Somata are adjacent to a proximal capillary, but bridging processes extend through the parenchyma to contact a distal capillary. **(b)** Ablation of a bridging pericyte. Inset in pre-ablation period shows target cell with laser scan path restricted to the soma (yellow line). Three days post-ablation, both proximal and distally contacted capillary become dilated due to loss of pericyte contact. **(c)** Ablation of a rarer “floating” pericyte, where the soma is located within the parenchyma and bridging processes extend from either end to contact nearby capillaries. In the acute timeframe of 2 hours, the pericyte is lost and there is a clear dilation in the upper capillary, and modest dilation in the lower capillary. At 3 days post-ablation, capillary dilation persists. At 21 days, pericyte contact is regained by growth of neighboring pericytes. Capillary tone is regained. Note that the structure of the upper capilary becomes less tortuous with coverage by neighboring processes.

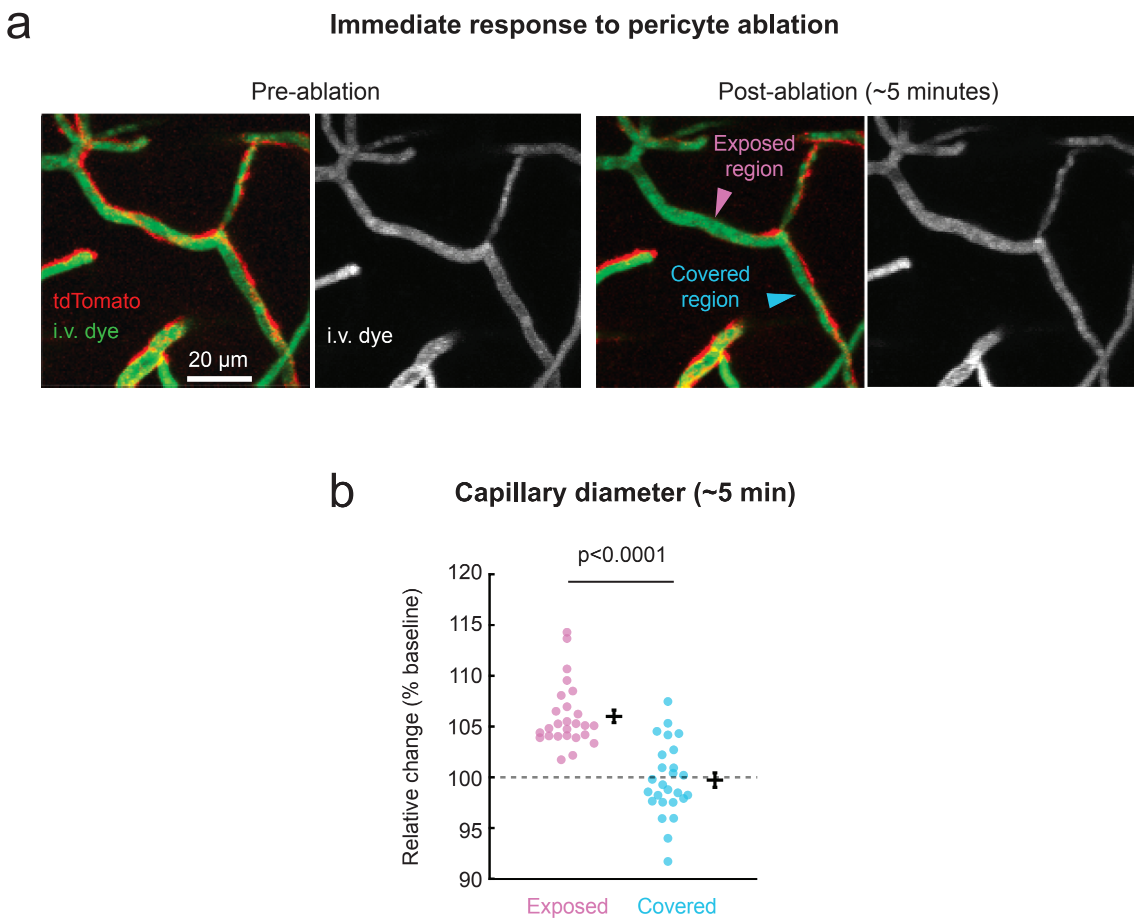


**Supplementary Figure 19. Acute dilation of capillaries following capillary pericyte ablation.** **(a)** A region of capillary bed covered by capillary pericyte processes (the pericyte soma targeted for ablation is outside the field of view). The same capillary region is then re-imaged within 5 minutes after cell ablation. Lumen diameter of the capillary region uncovered by pericyte ablation (exposed region) is compared to diameters in unperturbed regions covered by neighboring pericytes. **(b)** Relative change in capillary diameter measured ~5 minutes after capillary pericyte ablation, comparing regions exposed by the ablation procedure versus neighboring covered regions. $F(1,46)=46.48$, $P < 0.0001$ using repeated measures ANOVA. N = 26 (5 mice) capillaries.

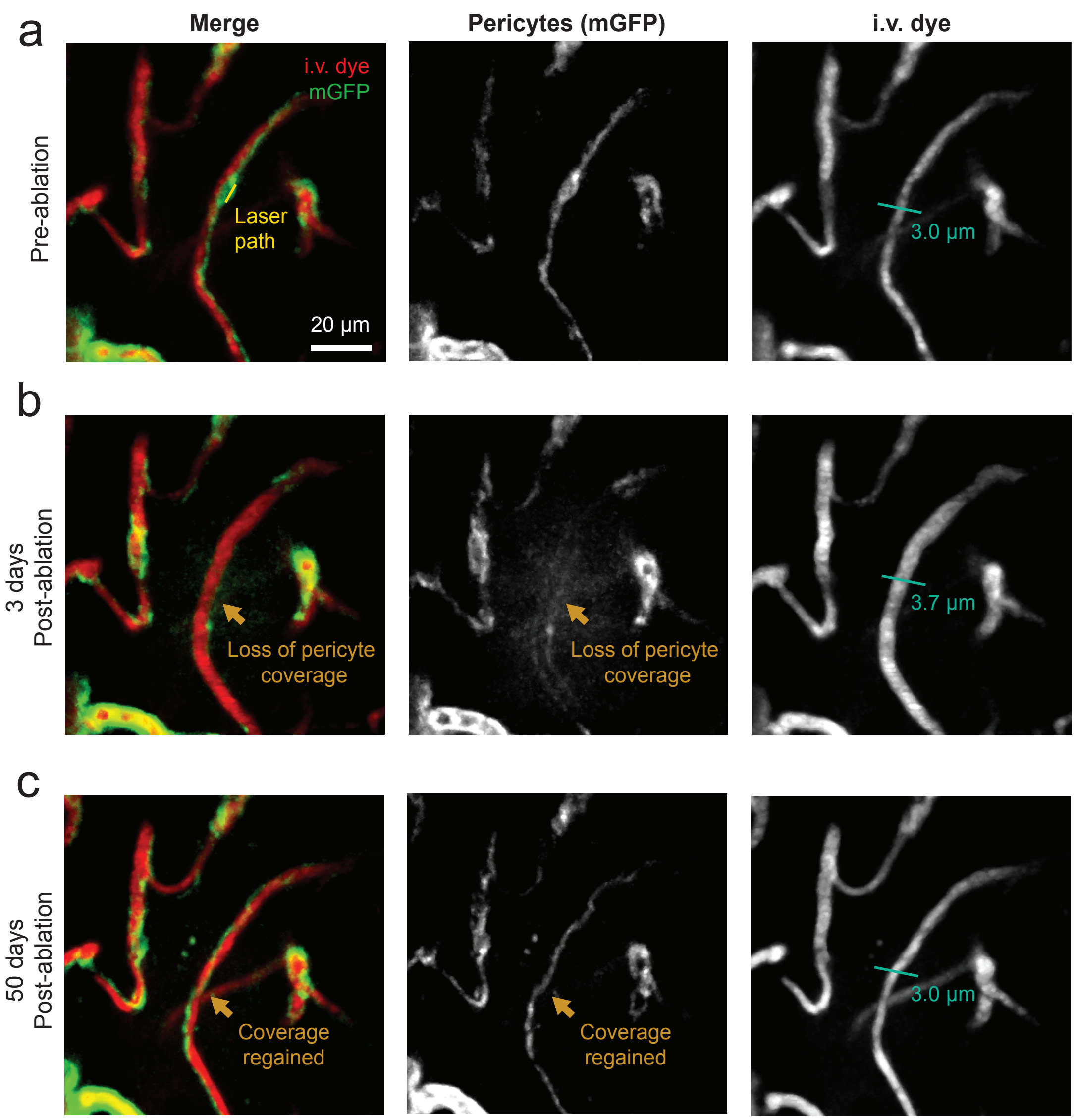


**Supplementary Figure 20. Capillary diameter returns to baseline with restoration of pericyte coverage. (a)** Before ablation, mGFP-positive pericyte processes contact the capillary endothelium. The path of the ablation laser is shown, targeting the pericyte soma (left panel). Before ablation, the full-width at half-max diameter of the target capillary is 3.0 µm (right panel). **(b)** Three days after ablation, the targeted pericyte soma and processes are absent (arrow). Pericyte contact is lost in the central portion of the image, but is maintained elsewhere (left panel). Capillary dilation is seen in the area left un-covered by the ablation (right panel). **(c)** Fifty days after ablation, the vessel has re-gained pericyte contact. Note that there is no pericyte soma where there once was in panel a, but processes of neighboring pericytes have grown into the territory to contact the capillary endothelium (left panel). Capillary diameter returns to the pre-ablation level of 3.0 µm (right panel). Surrounding capillaries are largely unchanged in comparison to the capillary experiencing pericyte ablation. This example is representative of all ablations in this study.

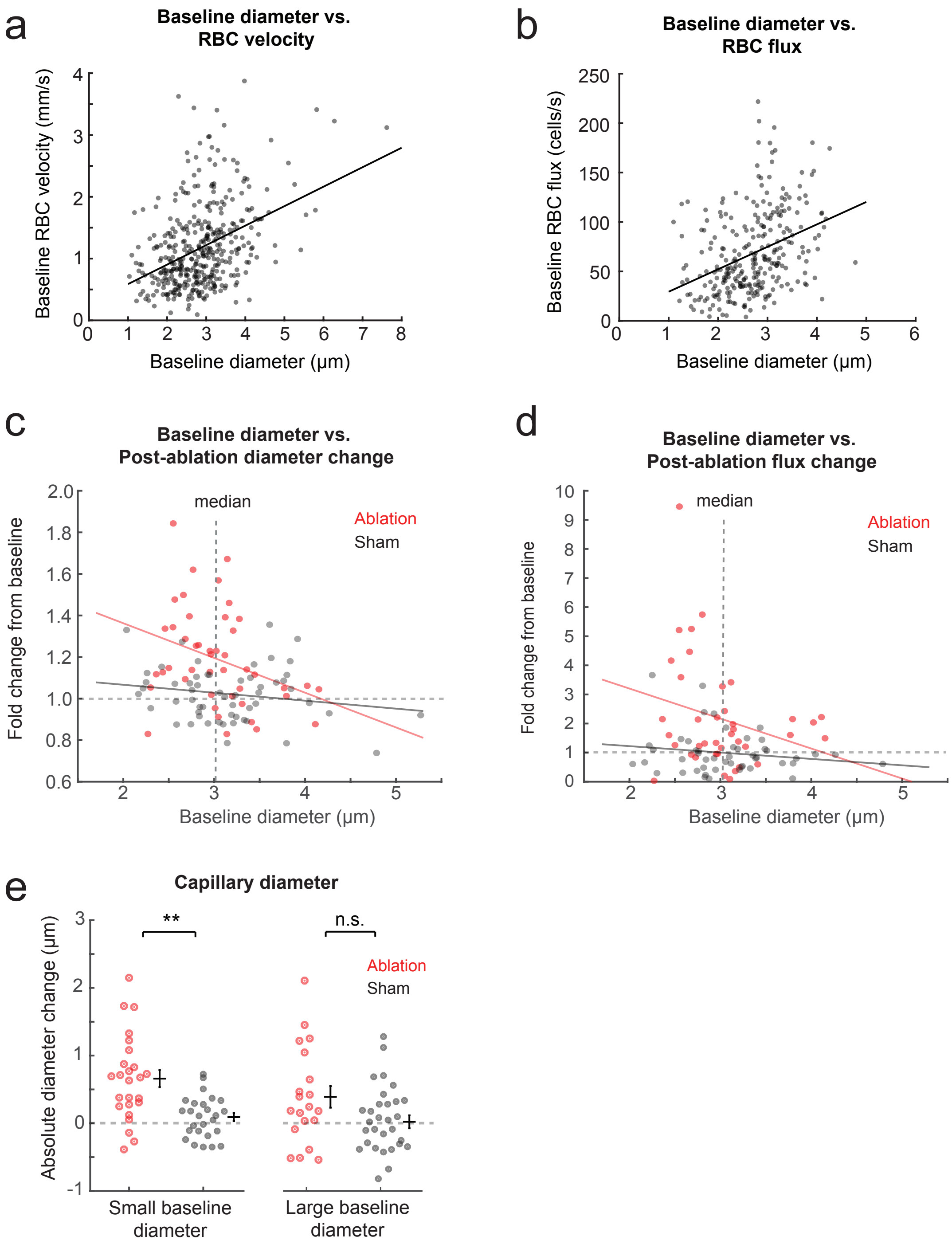


**Supplementary Figure 21. Capillary vasodynamics before and after capillary pericyte ablation.** **(a)** Baseline capillary RBC velocity plotted as a function of the diameter of the same capillary segment. All vessels are capillaries (5th - 9th branch order) compiled from ablation and optogenetic studies. Pearson's correlation: $R^2$ = 0.16, p<0.0001; n = 393 capillaries. Note that FWHM diameter calculations under-estimate the true capillary diameter. **(b)** Baseline capillary RBC flux plotted as a function of the diameter of the same capillary segment. Pearson's correlation: $R^2$ = 0.14, p<0.0001; n = 277 capillaries. **(c,d)** Fold change in lumen diameter (c) and RBC flux (d) following capillary pericyte ablation, plotted as a function of baseline capillary diameter (ablation-red, control-gray). Correlation analysis outcome: Diameter for ablation group ($R^2$ = 0.11, p = 0.03), diameter for control group ($R^2$ = 0.03, p = 0.17), flux for ablation group ($R^2$ = 0.07, p = 0.12), flux for control group ($R^2$ = 0.03, p = 0.25). Dotted vertical line shows median of ~3 µm for all data points, which was used to separate "small" and "large" baseline diameter capillaries. **(e)** Absolute diameter change after ablation was greater than sham for small, but not large capillaries. **p=0.0003, n=25 small ablation capillaries (mean 0.66 µm increase), n=26 small control; n.s. p>0.1, n=20 large ablation (mean 0.39 µm increase), 31 large control. Wilcoxon rank-sum test.
